# Siglec-G on B cells restrains the germinal center response by controlling T cell help during positive selection

**DOI:** 10.64898/2026.05.06.723371

**Authors:** Jhon R. Enterina, Sung-Yao Lin, Liany Luna-Dulcey, Susmita Sarkar, Edward N. Schmidt, Zeinab Jame-Chenarboo, Kennedy Chisholm, Arbab Ul Haq, Shu Luo, Chris D. St. Laurent, Theo Ataei, Fabrizio Giuliani, Olivier Julien, Matthew S. Macauley

## Abstract

The germinal center (GC) reaction requires tight regulation of B cell and T follicular helper (Tfh) cell interactions to ensure B cell expansion and antibody affinity maturation, while preventing oncogenesis. However, regulatory mechanisms fine-tuning B-T cell interactions within the GC to prevent aberrant activation and proliferation remain incompletely understood. Here, we identify Siglec-G, the mouse ortholog of human Siglec-10, as an immune checkpoint that restrains the GC by dampening B-T cell interactions. Selective and temporal ablation of Siglec-G on B cells after immunization triggers GC hyperplasia and enhanced plasma cell and antibody output. While Siglec-G is dispensable in B cell receptor (BCR)-mediated processes, it acts as an intrinsic inhibitory receptor of B-T cell interactions in the GC, ultimately limiting Myc and mTORC activation within positively selected GC B cells. *Trans* interactions of Siglec-G and its glycan ligands on Tfh likely contribute in fine-tuning the strength of bidirectional signaling following contact between GC B cells and Tfh cells. This interaction is further reinforced by glycan remodeling that occurs in the GC, resulting in concurrent decreased in glycan ligands on GC B cells and increased in glycan ligands on Tfh. This augmented binding of Siglec-G/10 on Tfh is mainly due to the upregulation of α2-6 linked sialic acid ligands. Moreover, APEX2-based proximity labeling revealed several candidate Siglec-G/10 binding partners on T cells, including BTLA, CD6, and Slamf6, which are known negative regulators of Tfh cell activation. Taken together, our findings identified that Siglec-G acts as a GC checkpoint receptor, restricting B cell proliferation by tuning T cell help following B-T cell interactions.

## INTRODUCTION

High-affinity antibodies are the product of the germinal center (GC) reaction(*1*). Cycles of somatic hypermutation in the dark zone (DZ) and affinity-based selection in the light zone (LZ), drive the progressive refinement of antibody affinity, a process known as affinity maturation. Positive selection of GC B cells with a B cell receptor (BCR) having higher affinity for its antigen is driven by two key signals: the first is activation of the BCR through antigen displayed on follicular dendritic cells and the second is cognate interactions with T follicular helper (Tfh) cells. These two signals synergize and converge on Myc and mTORC pathways to drive GC B cell proliferation and differentiation(*2–4*). Fine-tuning these signals is critical for not only fate determination of GC B cells into plasma cells (PC) and memory B cells (MBC), but also for safeguarding against the potential formation of malignant or autoreactive B cells. While regulatory T follicular helper (Tfr) cells are an important component(*5, 6*), immune checkpoints that regulate B-T cell interactions in the GC play an important role.

Previous studies have demonstrated that B cell activation via the BCR is reprogrammed in the GC to fine-tune the selection of affinity matured B cells(*7, 8*). Specifically, BCR activation to soluble antigens is significantly attenuated in MBC compared with their naive counterparts. This is in part due to reduced surface BCR expression as well as an altered AKT signaling network, leading to enhanced enzymatic activities of known BCR negative regulators such as CSK, HPK1, and SHP-1(*9*). Recently, we discovered that changes in cell surface glycosylation that occurs upon differentiation of naive B cells to GC B cells participates in the modulation of BCR signaling(*10*). One change in glycosylation relates to the emergence of the glycan epitope detected by the GL7 antibody(*11*). GL7 recognizes *N*-glycans on GC B cells that are capped with α2-6 linked sialic acid (*N*-acetylneuraminic acid; Neu5Ac)(*12*). Appearance of Neu5Ac on GC B cells is accompanied by the disappearance of *N*-glycolylneuraminic acid (Neu5Gc), which stems from suppression of the enzyme CMP sialic acid hydroxylase (CMAH). This switch in sialic acid elicits an effect through CD22, an inhibitory B cell co-receptor, as CD22 has a strong binding preference for Neu5Gc over Neu5Ac. As a consequence, this remodeling blunts BCR signaling in GC B cells through CD22(*12*). Mice that lack CD22 or block the appearance of the GL7 epitope on GC B cells result in aberrant BCR activation, which was associated with reduced GC B cell number, increased apoptotic cell death, and delayed maturation of antigen-specific antibodies following immunization(*10, 13*)

In addition to CD22, B cells express another sialic acid-binding immunoglobulin-type lectin (Siglec) known as Siglec-G. Siglec-G and its human ortholog Siglec-10 have well described roles as inhibitory co-receptors on B-1a B cells(*14*). Specifically, Siglec-G can recruit the phosphatases SHP-1 and SHP-2, following phosphorylation of its cytoplasmic immunoreceptor tyrosine-based inhibition motifs (ITIMs), to dampen BCR signaling uniquely in B1a B cells(*15*). The inhibitory role of Siglec-G in promoting B cell tolerance has been extensively characterized in the context of Siglec-G^KO^ mice, where a role for Siglec-G in B-1a cells has been demonstrated(*16*). Siglec-G-deficient mice displayed a spontaneous autoimmune phenotype characterized by autoantibody production, hyperactivated CD4^+^ T cells, and spontaneous GC formation(*17*). However, it is currently unknown whether a Siglec-G plays a functional role in conventional B-2 B cells. Unlike CD22, which binds exclusively to α2-6 linked sialic acid-containing *N*-glycans, the glycan ligands of Siglec-10 and Siglec-G are not as well understood, with some experimental support for recognition of glycans terminated by either α2-3 or α2-6 linked sialic acid(*18–20*). Notably, Siglec-G/10 have enhanced affinity towards Neu5Gc, although Neu5Gc is not naturally found in humans(*21*). Overall, the switch from Neu5Gc to Neu5Ac on GC B cells as well as the loss of core1 α2-3 linked sialic acid on *O*-glycans, which is evident from enhanced PNA staining, has the potential to impact Siglec-G. Nevertheless, a role for Siglec-G in the GC has not been explored.

Here, we demonstrate that Siglec-G plays a crucial role in regulating the GC response in mice following immunization. Temporal deletion of Siglec-G on B cells following immunization resulted in a significant expansion of the GC compartment, including both the GC B cells, Tfh cells, and an augmented antibody response due to increased PC output. While Siglec-G is dispensable for modulating BCR signaling in GC B cells, we identified that Siglec-G controls the activation of Myc and mTORC signaling pathways by restricting B-T cell interaction in the GC. Investigating Siglec-G/10 ligands in the GC revealed that GC B cells downregulate *cis* ligands, while there is an overall increase in ligands Tfh CD4^+^ T cells compared to naive CD4^+^ T cells. This remodeling promotes *trans* interactions between Siglec-G on B cells and its α2-6 and α2-3 linked sialoglycan ligands on Tfh cells. Overall, our findings highlight a new regulatory mechanism for Siglec-G in restraining T cell help within the GC to prevent aberrant GC B cell proliferation.

## RESULTS

### An expanded GC in Siglec-G^KO^ mice

BALB/c mice deficient for *Siglecg* display splenic hyperplasia with age due to significant expansion of GC B cells, CD4^+^ T cells, and PC, which were associated with the production of IgM and IgG autoantibodies(*17*). To assess whether a similar phenotype is observed on a C57BL6 (B6) genetic background, we examined the spleens of young and aged Siglec-G^KO^ B6 mice and their WT counterparts. While young male and female Siglec-G^KO^ B6 mice showed comparable splenic weight with their WT counterparts, aged Siglec-G^KO^ B6 mice exhibited splenic enlargement, and this was not sex-dependent (**Figure S1A,B**). Although there was no significant difference in splenic size, young Siglec-G^KO^ B6 mice displayed increased levels of follicular CD4^+^ T cells and GC B cells (**Figure S1C-K**). To assess whether expansion of GC B cells in Siglec-G^KO^ B6 mice is T cell-dependent, we disrupted the Tfh master regulator Bcl6(*22*), by crossing Siglec-G^KO^ and CD4^Cre^×*Bcl6*^f/f^ mice (**Figure S2A**). On a Siglec-G^KO^ background, GC B cells were completely abrogated in the absence of Bcl6 in CD4^+^ T cells as similarly seen in WT mice, demonstrating that formation of Siglec-G^KO^ GC B cells retain a requirement for Tfh cells (**Figure S2B,C**). Collectively, these data suggest that Siglec-G expression in mice is involved in regulation of the GC.

### Aged Siglec-G^KO^ mice have a shorter lifespan

An early study shows that expression of Siglec-G limits the formation of B cell-mediated lymphoproliferative disease in BALB/c mice(*23*). Given the significant increase in the splenic mass of Siglec-G^KO^ mice (**Figure S1B**), we aged WT and Siglec-G^KO^ mice to investigate signs of morbidity or mortality. Siglec-G^KO^ B6 mice had a shortened lifespan with a median survival of 776 days, whereas the majority of aged WT B6 mice survived the monitoring period (**Figure S3A-B**). Furthermore, Siglec-G^KO^ mice showed a steady expansion of spontaneously formed GC B and follicular CD4^+^ T cells with age (**Figure S3C-F**). These findings highlight the crucial role that Siglec-G plays in the overall survival of aged B6 mice, as well as its potential involvement in controlling the formation or maintenance of the GC compartment.

Given the splenomegaly and abnormal expansion of GC B cells in Siglec-G^KO^ B6 mice, we wondered if these B cells have tumorigenic potential. To test this, B cells were purified from spleens of aged WT and Siglec-G^KO^ B6 mice and adoptively transferred into immunodeficient Rag-1^-/-^ mice. Recipient mice transplanted with B cells from aged Siglec-G^KO^ mice displayed greatly reduced survival rate compared to Rag-1^-/-^ mice that had been transplanted with B cells from aged WT mice, suggesting that B cells from Siglec-G^KO^ mice are transplantable and potentially malignant (**Figure S3G-H**). Although detailed immunohistochemical analysis of malignant B cells were not carried out for these studies, the results demonstrate that aged B cells from Siglec-G^KO^ mice are transplantable.

### *SIGLEC10* is downregulated in Diffuse Large B cell Lymphoma (DLBCL) tumors

B cell-associated cancer is a heterogeneous form of hematological malignancy that can arise from different stages of B cell development and display distinct phenotypes and therapeutic challenges(*24*). Among the varied subgroups of the disease, diffuse large B cell lymphoma (DLBCL) constitutes over half of all B cell malignancy incidence in the world(*25*). DLBCL is an aggressive form of cancer that arises from aberrant mature B cells in the periphery. Given that aged Siglec-G^KO^ mice develop GC hyperplasia and signs of malignancy, we examined whether expression of the *SIGLEC10* gene is dysregulated in human DLBCL. Using publicly an available transcriptome dataset(*26*), we found that DLBCL tumors downregulate the expression of *SIGLEC10* compared to B cell subsets from healthy controls (**Figure S3I**). Notably, *SIGLEC10* expression levels were comparable between the two main subgroups of DLBCL: GC DLBCL and activated B cell (ABC) DLBCL (**Figure S3J**). To assess whether this dysregulated expression of *SIGLEC10* can predict survival outcome in DLBCL patients, we stratified the DLBCL tumors from patients enrolled in the International DLBCL Rituximab-CHOP Consortium Program Study based on their *SIGLEC10* expression levels(*27*). Overall (OS) and event-free (EFS) survival were measured from tumors with low (lower quartile; *n*=118) and high (upper quartile; *n*=118) *SIGLEC10* expression. Kaplan Meier plots revealed that patients with *SIGLEC10*^lo^ DLBCL tumors have more unfavorable survival outcomes than patients with *SIGLEC10*^hi^ DLBCL tumors (**Figure S3K-L**). These findings suggest that downregulation of *SIGLEC10* in lymphoma may also play a causative role in the development and progression of DLBCL.

To further explore the impact of *SIGLEC10* in the biology of DLBCL tumors, a gene set enrichment analysis (GSEA) was performed to identify any gene networks and pathways associated with *SIGLEC10* expression (**Figure S3M-O**). Here, we used another publicly available transcriptome dataset from Reddy *et al*. (*28*), consisting of 562 DLBCL tumors (**Figure S3M**). Tumors were categorized based on *SIGLEC10* expression and the transcriptome data from tumors in the lower and upper tertiles were assessed for gene signature enrichment using GSEA 4.0. *SIGLEC10*^lo^ DLBCL were significantly enriched for gene signatures relevant to oxidative phosphorylation and Myc signaling, which are both critical drivers for lymphoma (**Figure S3N**). On the other hand, *SIGLEC10*^hi^ DLBCLs showed significant enrichment for gene signatures associated with apoptotic cell death, JAK-STAT signaling, interferon response, and p53 pathway (**Figure S3O**). Overall, these findings highlight the potential regulatory function that Siglec-10 plays in human B cell DLBCL development and progression, implicating Siglec-10 as a tumor suppressor.

### Siglec-G expression on B cells contributes to restraining the GC compartment

Consistent with its established role in the B-1 lineage(*20, 29*), we observed Siglec-G expression on B-1a and B-1b cells, as well as on conventional B-2 cells and plasmacytoid dendritic cells (pDC) (**Figure S4**). To confirm whether the regulation of the GC response is strictly B cell-intrinsic, we sought to isolate its B cell contribution to an expanded GC. To do so, we generated a mixed bone marrow chimera by transplanting a 1:1 ratio of bone marrow cells from CD45.1^+^ WT and CD45.2^+^ Siglec-G^KO^ mice into lethally irradiated B6 recipient mice. At seven weeks post-reconstitution, chimeric mice were immunized with Nitrophenyl-Ovalbumin (NP-OVA) decorated liposomes via intravenous injection, and GC B cells were tracked at different timepoints after immunization (**Figure 1A**). Flow cytometric analyses of splenocytes showed that Siglec-G^KO^ GC B cells outcompeted WT GC B cells from the early phase of the GC response (**Figure 1B,C**). This finding suggests that spontaneous GC expansion in Siglec-G^KO^ mice is driven by loss of Siglec-G on B cells.

**Figure 1.**
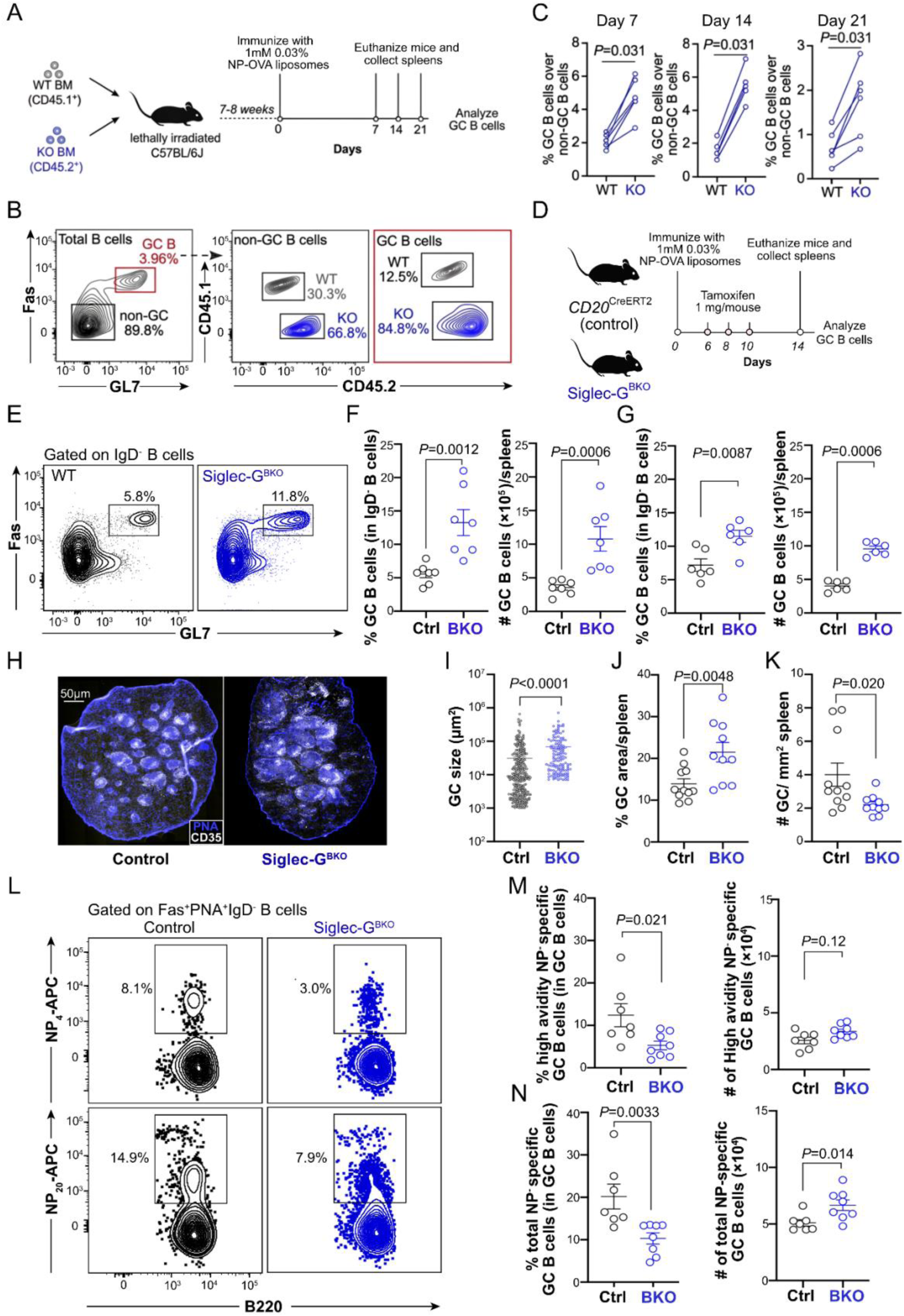
Expression of Siglec-G on B cells controls the GC reaction. (**A**) Experimental scheme for the TD immunization of mixed BM chimera mice reconstituted with 1:1 ratio of BM cells derived from CD45.1^+^ WT and CD45.2^+^ Siglec-G^KO^ B6 mice. Following immunization, mice were euthanized at different time points, and spleens were collected for flow analysis of GC B cells. **(B**) Flow cytometric gating strategy for the identification of WT and Siglec-G^KO^ non-GC and GC B cells from immunized mixed BM chimera mice. (**C**) Quantification of percent GC B cells over non-GC B cells on days 7, 14, and 21 post-immunization. Data are presented as paired line plots. (**D**) Experimental scheme for the immunization of control and Siglec-G^BKO^ mice with NP-OVA liposomes via tail vein injection. Mice were treated with 1 mg of tamoxifen/mouse on days 6, 8, and 10 post-immunization to induce a Cre-recombinase mediated deletion of *Siglecg* in the B cells. Mice were euthanized on days 14 and 21. Spleens were harvested for GC B cell analysis using flow cytometry. (**E-G**) Flow cytometric plots (**E**) and quantifications of percentages and absolute numbers of GC B cells in control and Siglec-G^BKO^ mice after 14 (**F**) and 21 (**G**) days post-immunization. (**H**) Representative immunofluorescence images of GC clusters in spleens derived from control and Siglec-G^BKO^ mice, 14 days post-immunization. (**H-K**) Quantification of individual GC cluster sizes (**I**), percent GC area per spleen section (**J**), and GC number per mm^2^ spleen section (**K**) from the experiment in (**H**). (**L-N**) Flow cytometric plots (**L**) and quantifications of percentages and absolute numbers of high avidity NP^+^ GC B cells (**M**) and total NP^+^ GC B cells (**N**) from control and Siglec-G^BKO^ mice, after day 14 post-immunization with NP-OVA liposomes. Data plots are presented as mean±SEM. Statistical analysis was performed using Mann-Whitney *U* test and Wilcoxon matched-pairs signed rank test.

While the data above suggest a B cell intrinsic function of Siglec-G in regulating the GC, it is still unclear whether this is due to the loss of an immunomodulatory function in immunogen-activated B cells or altered B cell functionality due to defects during early B cell development. To address this, we developed a Siglec-G floxed (*Siglecg*^f/f^) mouse model on a B6 genetic background by inserting two *loxp* sites before exon 2 and after exon 5 on the *Siglecg* allele. This mouse was crossed with *CD20*^CreERT2^ mouse to generate a tamoxifen drug-inducible *CD20*^CreERT2^×*Siglecg*^f/f^ mouse, denoted hereafter as Siglec-G^BKO^ mice, to enable temporal deletion of Siglec-G on B cells (**Figure S5A-C**). To investigate the impact of Siglec-G in the GC response using this new mouse model, we immunized male Siglec-G^BKO^ mice and their control counterparts (*CD20*^CreERT2^) with NP-OVA decorated liposomes. Tamoxifen was administered on days 6, 8, and 10 to delete Siglec-G on B cells, and GC B cells were analyzed from the splenocytes of immunized mice on days 14 and 21 post-immunization (**Figure 1D**). Siglec-G^BKO^ mice displayed a significantly expanded number of splenic GC B cells after days 14 and 21 post-immunization (**Figure 1E-G**). This GC expansion was not mediated by an increase in the number of GC clusters but due to an enlargement of individual GCs in Siglec-G^BKO^ mice relative to the controls (**Figure 1H-K**). When antigen-specific GC B cells were tracked, a relatively lower percentage of high affinity and total NP-specific GC B cells binding to NP_4_-APC and NP_20_-APC probes, respectively, were found in spleens of immunized Siglec-G^BKO^ than the controls (**Figure 1L-N**). However, the absolute numbers of antigen-specific GC B cells per spleen remained comparable between immunized control and Siglec-G^BKO^ mice, implying that loss of Siglec-G either enhanced the exit of antigen-specific GC B cells or expanded GC B cells with low or no apparent affinity to the immunizing antigen. Overall, these findings show that Siglec-G on GC B cells restricts the expansion of GC B cells.

### Siglec-G on B cells controls PC and antibody output but not MBC differentiation

A key function of the GC reaction is to select affinity matured B cell clones for MBC and PC differentiation. To interrogate whether an altered GC response in Siglec-G^BKO^ mice impacted antibody response to immunizing antigen, we compared the splenic PC and antibody responses between Siglec-G^BKO^ and control mice after immunization with NP-OVA liposomes (**Figure 2A and S6A**). We observed an increase in GC-derived plasma cells (B220^+^CD19^+^IgD^+^Fas^+^GL7^+^TAC-I^+^CD138^+^) on Days 14 and 21 post-immunization (**Figure S6B-D**). Immunofluorescence (IF) imaging further confirmed the augmented presence of CD138^+^ plasma cells within the GC area (**FigureS6E-F**). Specifically, spleens of Siglec-G^BKO^ mice produced more CD138^+^TACI^+^ PCs than controls (**Figure 2B,C**). Likewise, using an enzyme-linked immunosorbent spot (ELISPOT) assay, we identified that Siglec-G^BKO^ mice displayed an enhanced production of both splenic IgM^+^ and IgG^+^ PCs that secretes high affinity antibodies to NP hapten (**Figure 2D-G**). This increased PC output correlated with serum antibodies as titers of high affinity and total anti-NP polyclonal IgG1 antibodies were substantially higher in immunized Siglec-G^BKO^ than control mice (**Figure 2H-K**). However, antibody affinity maturation and BCR somatic hypermutation in NP^+^ GC B cells were similar between control and Siglec-G^BKO^ mice (**Figure 2L-N**). MBC output was also examined by flow cytometry on day 21 post-immunization. CD38^hi^IgD^-^B220^+^CD19^+^ MBCs were gated for high expression of both CD80 and PD-L2 receptors to identify a subset of MBCs that are believed to be mostly generated by the GC (**Figure S7A,B**)(*30*), which revealed comparable numbers of high affinity and total NP^+^ CD80^+^PD-L2^+^ MBCs based on their binding to NP_4_-APC and NP_20_-APC probes, respectively, between immunized Siglec-G^BKO^ and control mice (**Figure S7C-D**). Overall, these data highlight Siglec-G as a negative regulator for PC differentiation, while Siglec-G appears dispensable for MBC formation after primary challenge.

**Figure 2.**
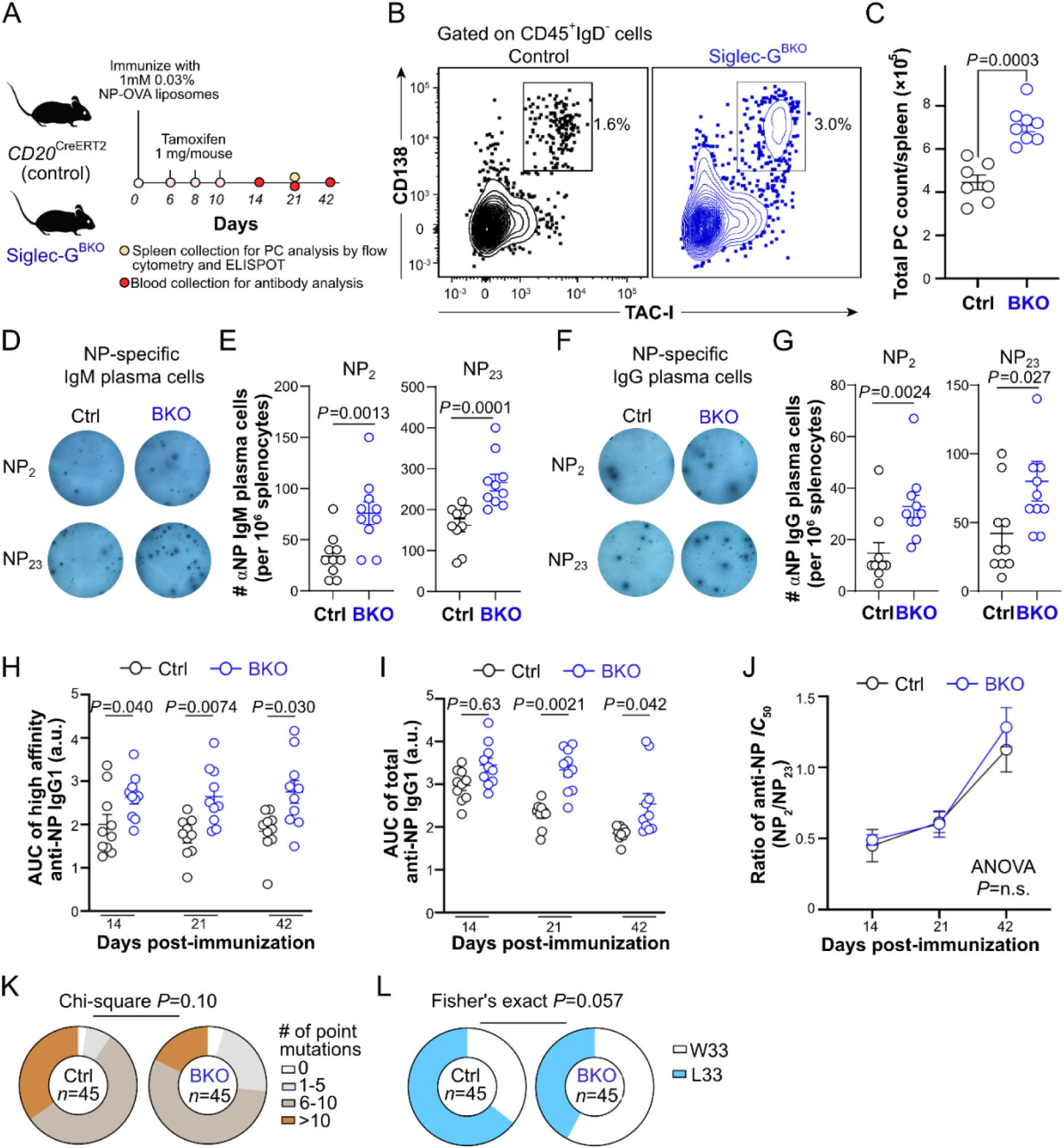
Siglec-G controls plasma cell and antibody production. (**A**) Experimental scheme for immunization of control and Siglec-G^BKO^ mice with NP-OVA liposomes. (**B**) Flow cytometric gating strategy for detecting plasma cells (CD138^hi^TACI^+^IgD^-^CD45^+^ cells) (**C**) Quantification of absolute plasma cell number in spleens of immunized control and Siglec-G^BKO^ mice after day 21 post-immunization. (**D**, **E**) Representative ELISPOT images (**D**) and quantifications of anti-NP_2_ and anti-NP_23_ IgM^+^ plasma cells (**E**) from spleens of control and Siglec-G^BKO^ mice, on day 21 post-immunization with NP-OVA liposomes. (**F**, **G**) Representative ELISPOT images (**F**) and quantifications of anti-NP_2_ and anti-NP_23_ IgG^+^ plasma cells (**G**) from spleens of control and Siglec-G^BKO^ mice, on day 21 post-immunization with NP-OVA liposomes. (**H**) ELISA of anti-NP IgG1 antibodies in sera of immunized control and Siglec-G^BKO^ mice binding to NP_2_-BSA. (**I**) Quantification of area under the curve (AUC) of ELISA plots in (**H**) as estimate of high avidity anti-NP IgG1 antibody titers. (**J**) ELISA of anti-NP IgG1 antibodies in sera of immunized control and Siglec-G^BKO^ mice binding to NP_23_-BSA (**K**). Quantification of AUC of ELISA plots in (**J**) as estimate of total anti-NP IgG1 antibody titers. (**L**) Estimation of affinity maturation of anti-NP IgG1 antibodies from sera of immunized control and Siglec-G^BKO^ mice. (**M**) Assessment of the number of mutations per clone in VH186.2 region of the BCR from control and Siglec-G deficient GC B cells, at day 21 post-immunization. (**N**) Distribution of high affinity mutation (W33L) in the VH186.2 region of the BCR from control and Siglec-G deficient GC B cells, at day 21 post-immunization. Data plots are presented as mean±SEM. Statistical analysis was performed using Mann-Whitney *U* test for **C**, **E**, and **G**, two-way ANOVA for **H**, **J**, and **L**, Kruskal-Wallis *H* test with post-hoc Dunn’s test for multiple comparison for **I** and **K**, and chi-square probability test and Fisher’s exact test for **I** and **J**, respectively.

### Siglec-G does not regulate the BCR in GC B cells

The observed expansion of GC B cells in immunized Siglec-G^BKO^ mice, supported by increased proliferation, suggests a B cell-intrinsic role for Siglec-G in regulating cell cycle entry. This is because sustained cell cycle entry and extensive cell division are intrinsically linked to driving plasma cell differentiation(*31, 32*). Therefore, we analyzed the effect of Siglec-G deficiency on a critical proliferative signal: BCR activation following antigen engagement. Expansion of GC B cells in immunized Siglec-G^BKO^ mice is, in part, due to enhanced proliferation as measured by *in vivo* BrdU labeling (**Figure S8**). As BCR signaling following antigen engagement is a critical signal for B cell proliferation in the GC and given the prior evidence that Siglec-G can repress the BCR in B1 lineage cells(*30*), we hypothesize that the enhanced proliferation of GC B cells lacking Siglec-G is mediated by increased activation downstream of the BCR. To test this, intracellular Ca^2+^ mobilization as well as phosphorylation of downstream proteins of the BCR were compared between control and Siglec-G-deficient GC B cells following *ex vivo* BCR crosslinking (**Figure 3A**). Intracellular Ca^2+^ levels showed no significant difference between control and Siglec-G-deficient GC B cells. Non-GC B cells from control and Siglec-G^BKO^ mice also displayed similar levels of intracellular Ca^2+^ post-stimulation (**Figure 3B,C**) Likewise, probing the phosphorylation of downstream proteins of the BCR complex (Syk, Akt, Erk1/2, and ribosomal protein S6) using phosFlow revealed a comparable phosphorylation profile between control and Siglec-G-deficient GC B cells at different timepoints post-stimulation (**Figure 3D-G**). These findings indicate that Siglec-G does not control BCR signaling in B2 and GC B cells, consistent with the previous findings in B2 cells(*20*).

**Figure 3.**
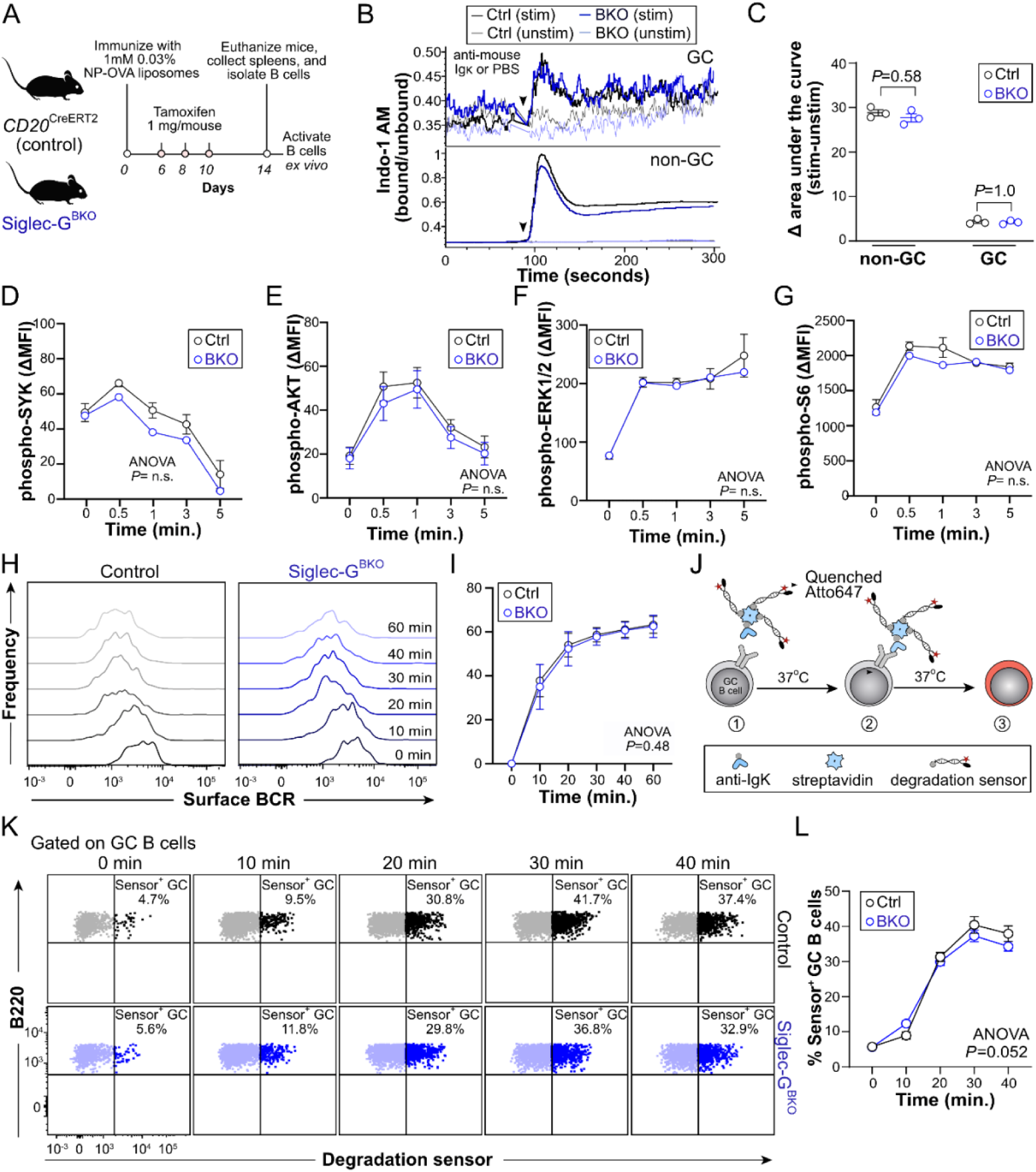
Siglec-G does not impact BCR signaling, BCR internalization, or antigen degradation following BCR crosslinking *ex vivo*. (**A**) Experimental scheme for immunization of control and Siglec-G^BKO^ mice with NP-OVA liposomes. On day 14 post-immunization, mice were euthanized, and spleens were harvested for mouse B cell isolation. Purified B cells were stimulated with anti-Igκ to assess BCR signaling, BCR internalization, and antigen degradation *ex vivo*. (**B**) Representative plots of time-dependent Ca^2+^ flux monitoring in GC and non-GC B cells from immunized control and Siglec-G^BKO^ mice using INDO-1 AM dye. (**C**) Quantification of AUCs from plots in (**B**) as a measure of the degree of Ca^2+^ mobilization between control and Siglec-G deficient B cells. (**D-G**) Phospho-flow analysis of pSYK (**D**), pAKT (**E**), pERK1/2 (**F**), and pS6 (**G**) at different timepoints (0, 0.5, 1, 3, and 5 minutes) following BCR stimulation. (**H**) Representative flow cytometric histograms of surface BCR levels on control and Siglec-G deficient GC B cells at different timepoint post-BCR stimulation. (**I**) Quantification of BCR internalization rates from BCR stimulated control and Siglec-G deficient GC B cells in (**H**). (**J**) Scheme describing the utilization of dsDNA-based degradation sensor to measure antigen degradation in B cells: 1) BCR ligation by surrogate antigen composed of biotin anti-Igκ, streptavidin, and biotinylated dsDNA-based degradation sensor with quenched Atto647, 2) incubation of B cells at 37 °C to internalize the degradation sensor, and 3) unquenching and increase fluorescence of Atto647 following antigen degradation. (**K**,**L**) Flow cytometric plots (**K**) and quantification (**L**) of percent control and Siglec-G deficient G C B cells positive for the degraded dsDNA sensor, which is a surrogate marker for antigen degradation. Data plots are presented as mean±SEM. Statistical analysis was performed using Kruskal-Wallis *H* test with post-hoc Dunn’s test for multiple comparison for **C** and two-way ANOVA for **D**, **E**, **F**, **G**, **I** and **K**.

We also tested whether enhanced expansion of GC B cells in immunized Siglec-G^BKO^ mice is a result of altered BCR internalization or antigen processing. To test BCR internalization, surface BCR were measured at different timepoints after B cell *ex vivo* stimulation with goat F(ab’)_2_ anti-mouse Igκ, which is a surrogate for a soluble antigen. Both the control and Siglec-G^BKO^ GC B cells internalized the BCR at similar rates, implying that Siglec-G does not impact proper uptake of the BCR-antigen complex (**Figure 3H,I**). To evaluate antigen processing, we used a surrogate soluble antigen that is composed of a 1:1:3 complex of biotinylated goat F(ab’)_2_ anti-mouse Igκ, streptavidin, and a biotinylated dsDNA-based degradation sensor(*33*), respectively. This dsDNA degradation sensor is fluorescently quenched in its undegraded state. Following BCR internalization, the BCR-bound complex is degraded in the lysosome and the dsDNA sensor is destroyed, leading to release and increase in fluorescence of the Atto647 fluorochrome(*33*). Control and Siglec-G^BKO^ GC B cells displayed similar ability to degrade antigen at different timepoints post-stimulation, suggesting that Siglec-G does not influence antigen degradation (**Figure 3J,K**). These findings rule out that Siglec-G expression on GC B cells impacts loading and presentation of peptide epitopes.

### Siglec-G on B cells regulates CD4^+^ T cell help during B-T cell interaction in the GC

Given that BCR-mediated processes are intact in Siglec-G-deficient GC B cells, we speculated that the enhanced GC response observed in immunized Siglec-G^BKO^ mice may be due to changes in the ability of GC B cells to process co-stimulatory signals delivered by cognate Tfh cells. To test this, tested acute T cell help elicited by αDEC205^OVA^ (*34*). Specifically, this was carried out in the context of a mixed BM chimera of CD45.1/2^+^ control mice and CD45.2^+^ Siglec-G^BKO^ mice. Chimera mice were immunized with NP-OVA liposomes followed by treatment with tamoxifen to delete Siglec-G on B cells derived from Siglec-G^BKO^ BM cells. On day 13, chimera mice were injected with either αDEC205^OVA^ antibody or PBS and analyzed after 15 hours (**Figure 4A**). In chimera mice treated with PBS, there was no significant difference in the distribution between control and Siglec-G^BKO^ in both DZ (CXCR4^hi^CD86^lo^) and LZ (CXCR4^lo^CD86^hi^) GC B cells. However, following treatment with αDEC205^OVA^ antibody, Siglec-G^BKO^ GC B cells considerably outcompeted their control counterparts in both DZ and LZ (**Figure 4B,C**).

**Figure 4.**
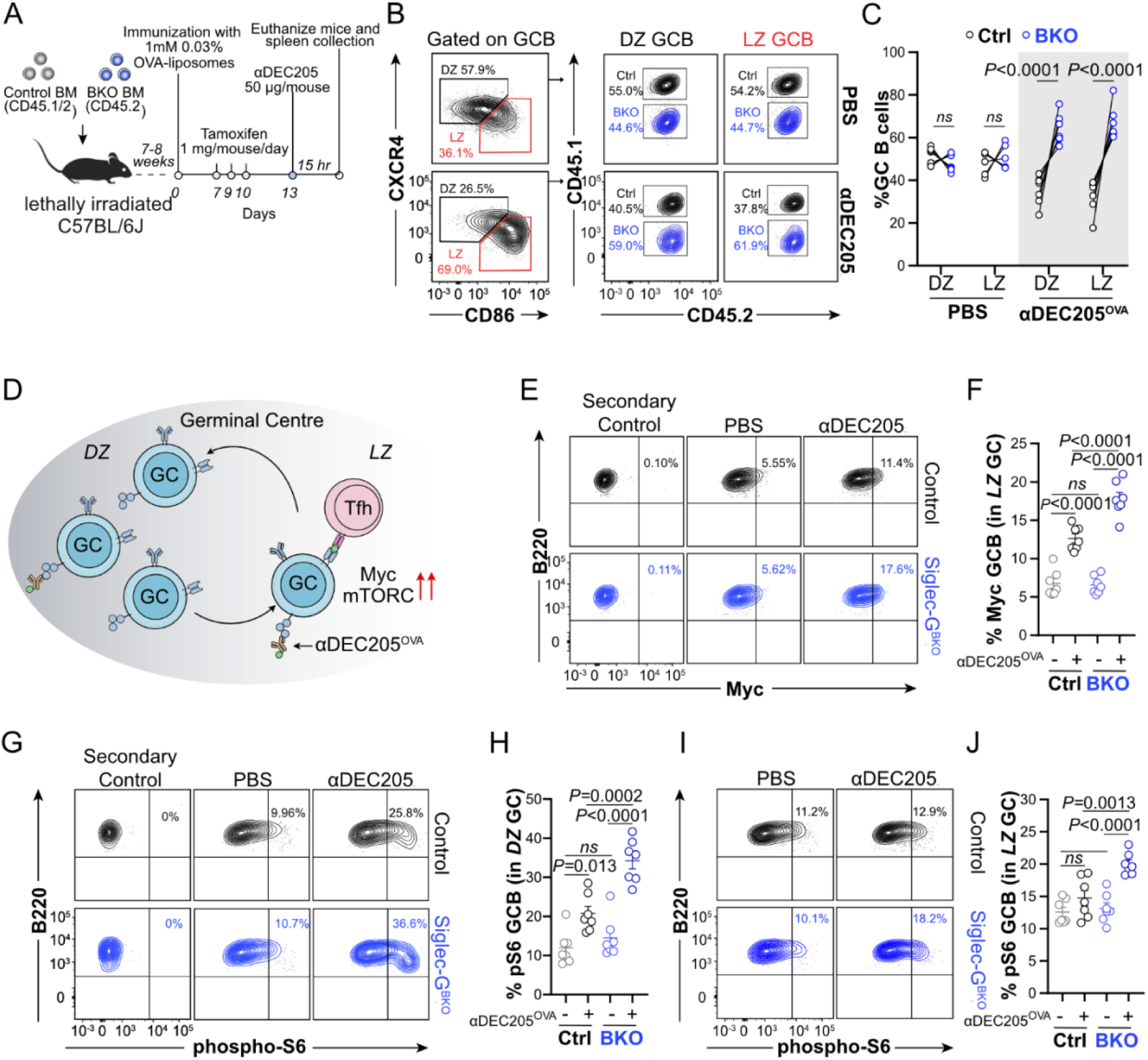
Siglec-G restricts Myc and mTORC signaling in GC B cells following B-Tfh cell interaction. (**A**) Experimental scheme for the induction of B-T cell interaction in vivo in immunized mixed bone marrow chimera mice transplanted with *CD20*^CreERT2^ (control) and Siglec-G^BKO^ bone marrow cells. (**B**) Flow cytometric gating strategy for the identification of control and Siglec-G^BKO^ GC B cells in the DZ and LZ compartments. (**C**) Quantification of percent control and Siglec-G^BKO^ GC B cells in spleens of chimera mice in **A** at day 14 post-immunization and 15 hours after treatment with either sterile PBS or αDEC205^OVA^. (**D**) Diagram showing the upregulation of Myc and mTORC pathways in GC B cells following αDEC205^OVA^-induced B-T cell interaction in the GC. (**E**, **F**) Flow cytometric plots (**E**) and quantification (**F**) of control and Siglec-G^BKO^ LZ GC B cells with high level expression of Myc transcription factor. (**G**, **H**) Flow cytometric plots (**G**) and quantification (**H**) of control and Siglec-G^BKO^ DZ GC B cells expressing high levels of phosphorylated ribosomal protein S6. (**I**, **J**) Flow cytometric plots (**I**) and quantification (**J**) of control and Siglec-G^BKO^ LZ GC B cells expressing high levels of phosphorylated ribosomal protein S6. Data plots are presented as paired line plots in **C** and mean±SEM in **F**, **H**, and **J**. Statistical analysis was performed using two-way ANOVA with post-hoc Sidak’s multiple comparisons test for **C** and one-way ANOVA with post-hoc Tukey’s multiple comparisons tests for **F**, **H**, and **J**.

We also examined whether Siglec-G impedes the delivery of co-stimulatory signals from Tfh cells following B-T cell interaction. To do this, we assessed the upregulation of the Myc transcription factor as well as the activation of the mTORC pathway, two of the critical signaling pathways activated in GC B cells during a productive B-Tfh cell interaction (**Figure 4D**)(*7*). Treatment with αDEC205^OVA^ antibody increased the population of LZ GC B cells with high Myc expression, however, the percentage of Myc^hi^ LZ GC B cells was significantly higher in Siglec-G^BKO^ GC B cells compared to their control counterparts, strongly suggesting the involvement of Siglec-G in dampening the Myc signaling pathway (**Figure 4E,F**). To evaluate mTORC signaling in GC B cells, we tracked the phosphorylation of ribosomal protein S6 (RPS6) as a surrogate marker for mTORC activation. RPS6 is a component of the 40S ribosomal subunit and is downstream of the mTORC1-ribosomal protein S6 kinase (S6K) axis(*4*). Administration of αDEC205^OVA^ in immunized chimera mice resulted in a significantly expanded Siglec-G^BKO^ GC B cell populations, in both DZ and LZ compartments, with higher levels of phosphorylated RPS6 than their control counterparts (**Figure 4G-J**). Overall, these findings highlight the role of Siglec-G in restraining T cell help in GC B cells by directly or indirectly modulating Myc and mTORC signaling.

### Loss of Siglec-G on B cells alters the distribution of Tfh and Tfr cells in the GC

The *in vivo* experiments above strongly suggest the involvement of Siglec-G in B-T cell interactions in the GC. Given that Siglec-G is dispensable in BCR-mediated signaling, it is plausible that Siglec-G controls Myc and mTORC pathways by either restricting activation signals of B cell co-stimulatory receptor such as CD40 or by altering the phenotype of follicular T cells through *trans* interactions with immunomodulatory receptors on CD4^+^ T cells. To test the latter hypothesis, we began by interrogating the general distribution of Tfh and Tfr cells, two specialized subsets of CD4^+^ T cells critical for mounting a robust GC response against T-dependent antigens(*35–37*). Following OVA-immunization, the frequency and total number of Tfh cells (CD4^+^CD44^+^PD-1^+^CXCR5^+^Foxp3^−^ T cells) in the spleen of Siglec-G^BKO^ mice were comparable with their control counterparts. By contrast, Tfr cells (CD4^+^CD44^+^PD-1^+^CXCR5^+^Foxp3^+^) displayed a substantial increase in number within Siglec-G^BKO^ mice (**Figure 5B-F**). To examine whether this observation is correlated with the frequency of antigen-specific follicular T cells, we used a fluorescently labeled I-Ab tetramer loaded with the OVA_329-337_ peptide to detect OVA-specific CD4^+^ T cells. GC B cells from Siglec-G^BKO^ mice had a higher frequency of both OVA-specific Tfh and Tfr cells than control mice, suggesting that B cell Siglec-G is involved in regulating expansion, recruitment, or retention of follicular CD4^+^ T cells in the GC.

**Figure 5.**
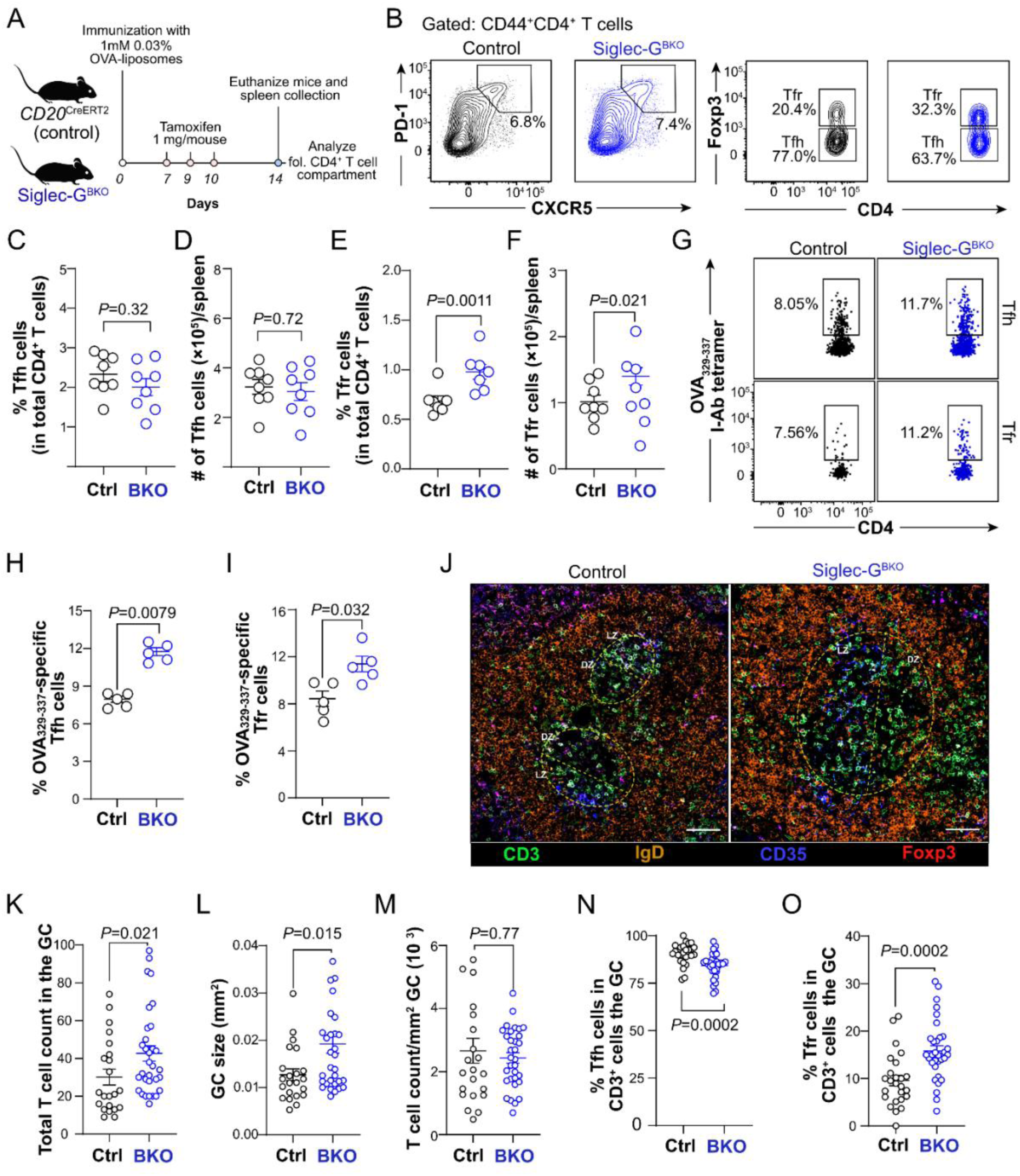
Loss of Siglec-G on B cells alters the balance between Tfh and Tfr cells in the GC. (A) Experimental scheme for immunization of control and Siglec-G^BKO^ mice with NP-OVA liposomes. (**B-F**) Flow cytometric plots (**B**) and quantification of percent and absolute numbers of Tfh cells (**C, D**) and Tfr cells (**E,F**). Flow cytometric plots for Tfh cells (**G**) and quantification of percent Tfh (**H**) and OVA-Tfr (**I**) cells specific for OVA_329-337_ I-A^b^-tetramer. (**J-O**) Confocal microscopic images (**J**) and quantifications of total CD3^+^ cells in the GC (**K**), GC size in mm^2^ (**L**), normalized CD3^+^ T cell count/mm^2^ of GC (**M**), percent Foxp3^-^CD3^+^ GC-Tfh cells (**N**), and percent Foxp3^+^CD3^+^ GC-Tfr cells (**O**) between immunized control and Siglec-G^BKO^ mice. Data plots are presented as mean±SEM. Statistical analysis was performed using Mann-Whitney *U* test. The scale bar in J represents 50 µm.

To evaluate the spatial distribution of follicular CD4^+^ T cells within the GC, we imaged splenic tissue sections from immunized control and Siglec-G^BKO^ mice by confocal microscopy (**Figure 5J**). GCs were visualized as IgD^-^ sites within the B cell follicle. DZ and LZ GC compartments were identified by CD35^-^ and CD35^+^ staining, respectively, while GC Tfh and Tfr cells were marked as CD3^+^Foxp3^-^ and CD3^+^Foxp3^+^ T cell populations, respectively. Microscopic analysis revealed that immunized Siglec-G^BKO^ mice harbored more CD3^+^ T cells per GC than control counterparts, and this observation was correlated with larger GC size in Siglec-G^BKO^ mice (**Figure 5K-M**). Intriguingly, this increase in GC-resident T cells was mainly attributed to increased frequency of GC Tfr cells in spleens of Siglec-G^BKO^ mice (**Figure 5N,O**). When the distribution of follicular T cells in different GC compartments was assessed, we observed that the number of Tfh cells in the DZ was comparable between groups, LZ Tfh cells were significantly higher in Siglec-G^BKO^ mice than their control counterparts. Likewise, Tfr cell numbers were increased in both DZ and LZ in Siglec-G^BKO^ mice (**Figure S9A,B**). When normalized against individual GC sizes, Tfr abundance/GC area in the DZ was higher in Siglec-G^BKO^ mice than controls while Tfh and LZ Tfr were similar between groups (**Figure S9C**). Collectively, these imaging data reveal that Siglec-G on B cells controls Tfh and Tfr numbers in the GC.

### Siglec-G inhibits activation of follicular CD4^+^ T cells

Entry of follicular CD4^+^ T cells partly depends on the strength of B-T cell interaction(*38*). It is possible that loss of Siglec-G on B cells enhanced the signaling strength and prolonged the duration of the cognate interactions between B cells and CD4^+^ T cells in the GC. We interrogated whether Siglec-G on B cells impacts the activation of follicular T cells. First, we characterized the expression of activation-induced marker CD69 on follicular CD4^+^ T cells from immunized control and Siglec-G^BKO^ mice (**Figure S10A,B**). We used the marker CD90.2 to distinguish between GC and non-GC resident Tfh cells by flow cytometry, with CD90.2^low^ Tfh cells representing those inside the GC(*^39^*) (**Figure S10C**). Our results showed that while majority of Tfh cells in both control and Siglec-G^BKO^ mice expressed CD69, their distribution was altered in Siglec-G^BKO^ mice. The frequency of CD69^+^CD90.2^lo^ Tfh cells was significantly higher in Siglec-G^BKO^ than control mice while an opposite trend was observed in CD69^+^CD90.2^hi^ Tfh cells (**Figure S10D,E**). Likewise, Tfr cells from Siglec-G^BKO^ mice showed the same trend as the Tfh compartment. Quantification of CD69 expression revealed that CD90.2^lo^ Tfh cells from Siglec-G^BKO^ mice showed a modest, yet significant, increase in CD69 expression compared to CD90.2^lo^ Tfh cells from control mice (**Figure S10F**). On the other hand, CD90.2^hi^ Tfh and Tfr cells from both control and Siglec-G^BKO^ mice showed comparable CD69 protein levels. This finding indicates that Siglec-G on B cells may regulate activation and GC localization of both Tfh and Tfr cells. To investigate this enhanced activation state at the intracellular level, we evaluated the expression of critical transcription factors associated with TCR signaling. Consistent with the surface activation profile, Tfh cells from Siglec-G^BKO^ mice exhibited significantly increased protein levels of Bcl6, the master transcriptional regulator of Tfh differentiation, as well as NFATc1, a key transcription factor driven by sustained TCR engagement (**Figure S10G-J**).

We also tested expression of selected immunomodulatory co-receptors critical for Tfh cell function (**Figure S10K-M**). We found that both Tfh and Tfr cells from Siglec-G^BKO^ mice showed significantly increased expression of PD-1, CD84 (SLAMF5), ICOS, CD73, and CD120b (TNFR2) on CD90.2^hi^ Tfh cells; CD84, CTLA4, ICOS, CD73, and CD120b on CD90.2^lo^ Tfh cells; and PD-1, ICOS, and CD73 on Tfr cells. Given that sustained expression of key costimulatory receptors requires T cell activation through constant B-T cell interactions(*39, 40*), this observed upregulation of distinct Tfh-associated proteins implies that follicular CD4^+^ T cells in Siglec-G^BKO^ mice displayed a more activated phenotype than their control counterparts.

Lastly, we assessed whether Siglec-G expression on B cells dampen the activation of Tfh cells following B-T contact formation cell *ex vivo*. To test this, isolated WT and Siglec-G^KO^ B cells were pulsed with OVA_323-339_ peptide. These cells were then co-culture with purified OT-II Tfh cells, which possess a transgenic T cell receptor specific for the OVA antigen. To generate these antigen-specific Tfh cells, B6 mice were adoptively transferred with OT-II T cells and subsequently immunized with OVA-liposomes (**Figure S11A**). After two hours of co-culture, we quantified the frequency of B-T cell pairs and levels of CD40L on the Tfh cells. Our results show that the percentage of B-T conjugates was comparable between genotypes (**Figure S11B,C**), indicating that Siglec-G does not impact the physical ability of B and T cells to form stable contacts. Conversely, we observed that OT-II Tfh cells cultured with Siglec-G^KO^ B cells induced higher surface expression of CD40L than OT-II Tfh cells cultured with WT B cells (**Figure S11D,E**), suggesting that Siglec-G controls the stimulation of cognate Tfh cells potentially through interaction with glycosylated immunomodulatory receptors present on Tfh cells. These data suggest that Siglec-G fine-tunes follicular CD4^+^ T cell activation independent of the strength of B-T cell contacts.

### Single cell transcriptome of follicular CD4^+^ T cell compartments between Ctrl and Siglec-G^BKO^ mice

To further investigate the role that B cell Siglec-G plays on Tfh and Tfr cells, we performed single cell RNA sequencing (scRNA-seq) on FACS-sorted follicular CD4^+^T cells from the spleens of immunized control and Siglec-G^BKO^ mice. After stringent quality control and data processing, Tfh (Foxp3^-^) and Tfr (Foxp3^+^) cells were subdivided for unsupervised clustering (**Figure S12A**). Within the Tfh compartment, 9 distinct clusters were identified and visualized using the uniform manifold approximation and projection (UMAP) (**Figure S12B**). All clusters were populated by Tfh cells from both control and Siglec-G^BKO^ mice, 5 clusters (0, 1, 2, 3, and 4) displayed frequencies greater than 10% from either group (**Figure S12C**). Cluster 0 was characterized by the high expression of *Tnfrsf4* (OX40), *Tnfrsf18* (GITR), and Ikaros family transcription factors *Ikzf2* and *Ikzf4* (**Figure S12D**). Gene ontology (GO) enrichment analysis of cluster 0 differentially expressed genes (DEGs) revealed an enrichment for biological processes involved in cilium organization, NF-kappaB signaling, and GTPase-mediated signaling (**Figure S13**). Cluster 1, defined by the upregulation of *Lmo4*, *Asb2, Il4,* and *Pde3b*, was associated with MAPK signaling as well as phosphatidylinositol phosphate and branched chain amino acid metabolism (**Figure S13**). Clusters 2 and 3 were expanded in Siglec-G^BKO^ mice, populated by 25.1% and 31.1% of the Siglec-G^BKO^ Tfh cells, respectively, compared to only 16.4% and 8.3% of Tfh cells from control mice. Cluster 2 Tfh cells displayed the highest expression for *Gpm6b*, *Cacna1d*, *Tnfsf8* (CD30L) and *Il21* (**Figure S12D**), with DEGs converging to pathways associated with ribosome biogenesis and the CD40 signaling (**Figure S13**). On the other hand, cluster 3 showed upregulated for *Rflnb*, *H2-Q2*, *Gm4814*, and *Tubb2a* (**Figure S12D**), with significant DEGs enriched for for cytoplasmic translation and ribosome biogenesis (**Figure S13**). Strikingly, cluster 4 displayed the most substantial difference in cell frequency between groups, constituting only 0.9% of the Siglec-G^BKO^ Tfh compared to around 13% of the control Tfh cells. This cluster was uniquely characterized by genes crucial for T cell cytotoxicity, including *Nkg7, Gzmk, Eomes*, and *Ccl5* (**Figure S12D**), and was enriched for genes related Type II interferon signaling, T cell activation, and integrin signaling (**Figure S13**).

Within the Tfr compartment, unsupervised clustering identified 6 clusters, 3 of which (0, 1, and 2) showing frequencies >10% from either group (**Figure S14A,B**). Tfr cluster 0 represented the majority of control cells (62.1%). It was characterized by the highest expression of *Cd226*, *Cd96*, and *St6galnac3* (**Figure S14C**); and DEG gene enrichment for pathways involved in IL-18 signaling and T cell activation (**Figure S15**). Meanwhile, Tfr clusters 1 and 2 are populated by 31.2% and 32.4% of total Siglec-G^BKO^ Tfr cells, respectively, but only represented by 19.5% and 12.1% of control Tfr cells (**Figure S14B**). Cluster 1 showed upregulated expression for *Hif1a*, *Tbc1d4, and Pou2af* (**Figure S14C**). Cluster 2 was defined by increased expression of Neb, Prdm1, Capg, and Arl5a, while also expressing the highest levels of Il10 compared to all other clusters (**Table S1**). Furthermore, DEG gene enrichment for pathways involved in the negative regulation of JAK-STATsignaling and apoptotic processes (**Figures S14C** and **S15**). Altogether, results from scRNA-seq of follicular CD4^+^ T cells reinforced our *in vivo* data showing that expression of Siglec-G on B cells can partly control the phenotypic states of both Tfh and Tfr cells, potentially shaping the overall response of these T cell populations in the GC reaction.

### Siglec-G/10 glycan ligands are remodeled in the GC

Alterations in CD22 ligands regulate the GC response(*8*), therefore, we wondered if there may also be changes to Siglec-G ligands with the GC. As all Siglec-G-Fc proteins explored to date function poorly as probes, we used Siglec-10-Fc because Siglec-G and Siglec-10 have a similar glycan ligand binding specificity and the ability of Siglec-10-Fc to recognize sialoglycan ligands on cells is well established(*20, 41, 42*). For these studies, we employed a Siglec-10-Fc mutant (R119A), which we refer to as Siglec-10R-Fc, as a control; this arginine is critical for sialic acid recognition in the Siglec family(*43*). On B cells, Siglec-10-Fc binding was significantly lower to both LZ and DZ GC B cells compared to naive B cells (**Figure 6A**). To assess if this downregulation is conserved in humans, we assessed Siglec-10 ligands on human naive and GC B cells from tonsils (**Figure S16A**). Here, we used our recently developed Siglec-liposomes for enhanced sensitivity and to avoid detection of by adsorbed human IgG from primary cells(*44*), and found that GC B cells have lower Siglec-10 ligands (**Figure S16B**). Therefore, downregulation of Siglec-G/10 ligands on GC B cells is conserved between mice and humans.

**Figure 6.**
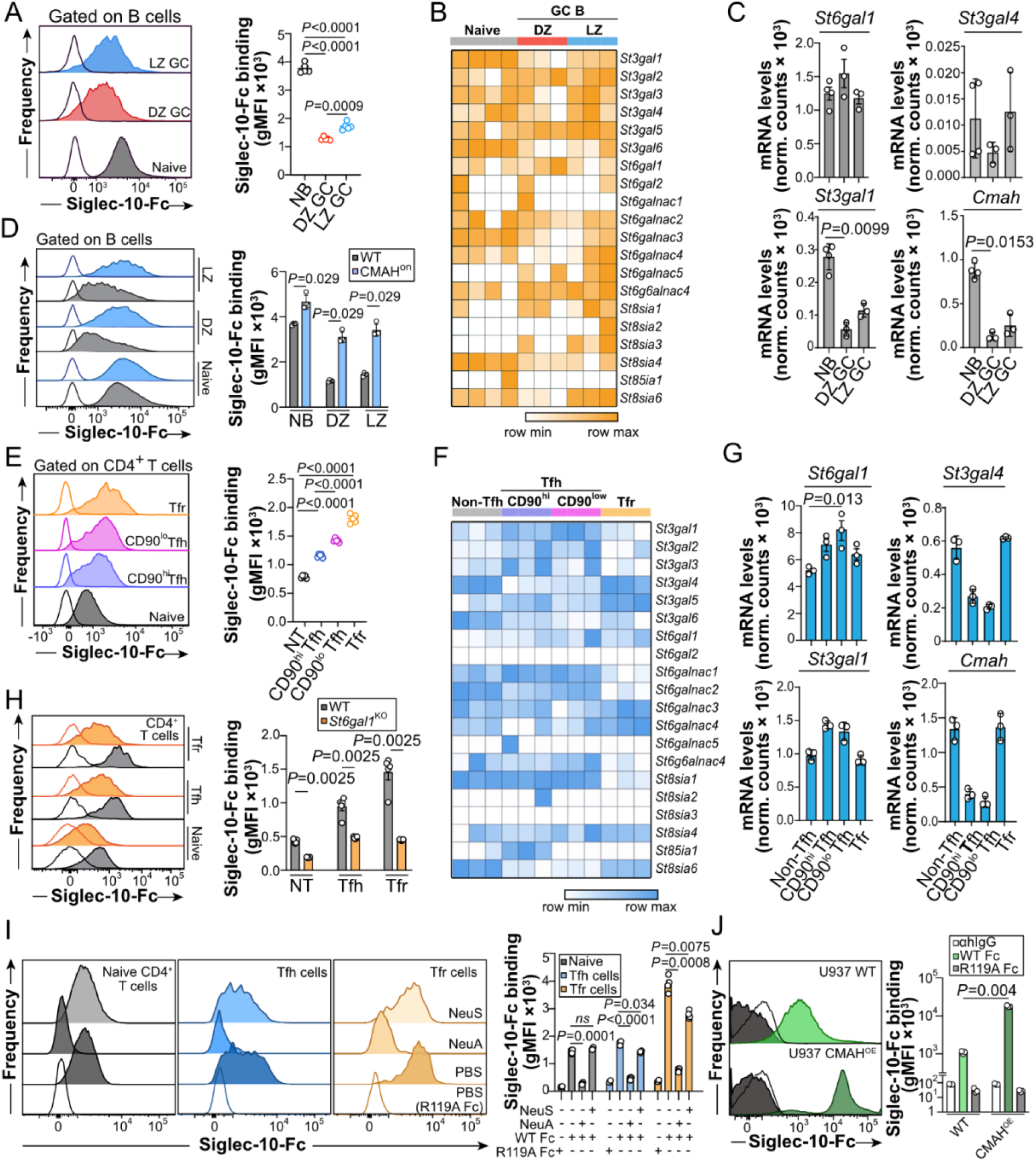
Siglec-10 glycan ligands are reprogrammed on GC B cells and follicular CD4^+^ T cells. (**A**) Flow cytometric histograms and quantification of Siglec-10-Fc binding on GC B cells. (**B**) Heatmaps of sialyltransferase transcript expression in mouse GC B cells derived from publicly available datasets. (**C**) Normalized transcript levels of *St6gal1*, *St3gal1*, *St3gal4* and *Cmah* genes in B cells (naive and GC)(*118*). (**D**) Flow cytometric histograms and quantification of Siglec-10-Fc binding on naive and GC B cells in CMAH^on^ mice. (**E**) Flow cytometric histograms and quantification of Siglec-10-Fc binding on follicular CD4^+^ T cells. (**F**) Heatmaps of sialyltransferase transcript expression in mouse follicular CD4^+^ T cells derived from publicly available datasets(*39*). (**G**) Normalized transcript levels of *St6gal1*, *St3ga1*, *St3ga4,* and *Cmah* genes in follicular CD4^+^ T cells. (**H**) Flow cytometric histograms and quantification of Siglec-10-Fc binding on control (CD4^Cre^) and St6Gal1-deficient mouse T cells. Shaded and line histogram plots represent binding of WT and their sialic acid non-binding mutants Fc, respectively. (**I**) Flow cytometric histograms and quantification of Siglec-10-Fc binding on NeuA and NeuS treated mouse CD4^+^ T cells. (**J**) Flow cytometric histograms and quantification of Siglec-10-Fc binding on WT and CMAH-overexpressing U937 cell lines. Filled histograms represent Siglec-10-Fc binding. Line histograms represent the sialic acid non-binding mutant (R119A) control. Data plots are presented as mean±SEM. Statistical analysis was performed using Mann-Whitney *U* test and Friedman test.

To identify potential enzymes involved in downregulation of Siglec-G/10 ligands, we analyzed transcript levels of sialyltransferases in naive and GC B cells (**Figure 6B**). Focusing on *St6gal1*, *St3gal1*, *St3gal4*, and *Cmah*, given their potential to create Siglec-G/10 ligands, we find that *St3gal1* and *Cmah* expression was decreased in DZ and LZ GC cells relative to naive B cells, while *St6gal1* and *St3gal4* levels remained unchanged (**Figure 6C**). We tested these candidates by using B cells from *St3gal1*^KO^, *St3gal4*^KO^, and CMAH^ON^ mice(*10, 45–47*). While Siglec-10-Fc binding to St3gal1^KO^ and St3gal4^KO^ cells was unaffected (**Figure S17A,B**), CMAH^ON^ B cells, which do not allow for downregulation of CMAH on GC B cells(*10*), exhibited a significant higher Siglec-10-Fc binding of Siglec-10-Fc (**Figure 6D**). These data indicate that the downregulation of CMAH, and the subsequent reduction of Neu5Gc-containing glycans, is the primary driver of the reduced Siglec-10-Fc staining profile observed on GC B cells.

In CD4^+^ T cells, Siglec-10-Fc binding was significantly increased on CD90^hi^ Tfh, CD90^low^ Tfh, and Tfr cells relative to naive cells (**Figure 6E**). This trend was conserved in human tonsillar CD4^+^ T cell subsets (**Figure S16C,D**). To analyze the enzymatic basis of upregulation, we profiled sialyltransferase expression in GC CD4^+^ T cell subsets (**Figure 6F**). *St6gal1* was significantly upregulated across follicular subsets compared to non-Tfh cells, *St3gal1* expression increased specifically in Tfh cells, while *St3gal4* expression was reduced (**Figure 6G**). Consistent with the increased *St6gal1* expression in Tfh and Tfr cells, CD90^hi^ Tfh and Tfr cells displayed higher SNA staining, which reflects higher levels of α2-6 sialosides (**Figure S18**). Expression of *Cmah* gene was modestly reduced in follicular CD4^+^ T cells, ruling out altered *Cmah* expression as being responsible for enhanced Siglec-G/10 ligands on Tfh cells. Validation of these results was performed using knockout mice and neuraminidase. A significant decrease in Siglec-10-Fc binding was observed in St6gal1^KO^ CD4^+^ T cells across naïve, Tfh, and Tfr populations (**Figure 6H**), whereas no change was seen in St3gal1^KO^ or St3gal4^KO^ T cells (**Figure S17C-E**), implicating α2-6-linked sialoglycans as important Siglec-10/G ligands on CD4^+^ T cells. We examined Siglec-10-Fc staining after pre-treated with Neuraminidase A (NeuA) or Neuraminidase S (NeuS), which removed all sialic acid linkages or only α2-3 sialic acid linkages, respectively. Siglec-10-Fc staining was reduced in all three NeuA-treated T cell subsets, whereas NeuS only caused a decrease in Siglec-10-Fc binding to Tfh and Tfr subsets (**Figure 6I**). It is noteworthy that NeuS did not diminish Siglec-10-Fc binding to the same extent as NeuA, suggesting that both α2-3 and α2-6 sialosides contribute to Siglec-10/-G ligands on Tfh cells, but that α2-6 sialosides dominate.

To further investigate Siglec-10/-G ligands, we used U937 cells as we have previously used glyco-engineered U937 cells successfully to make insights into the biosynthesis of Siglec ligands(*48*). Specifically, we used Cosmc^KO^ and Mgat1^KO^ cells, as these will lack complex type mucin-type *O*-glycans and *N*-glycans, respectively. Siglec-10-Fc binding remained intact on Cosmc^KO^ cells but was significantly reduced on Mgat1^KO^ cells, indicating that Siglec-10 glycan ligands are primarily presented on complex *N*-glycans (**Figure S19**). Furthermore, we confirmed that overexpressing CMAH, the enzyme that converts Neu5Ac to the non-human Neu5Gc sialic acid, led to higher Siglec-10-Fc binding (**Figure 6J**). Altogether, our results demonstrate that Siglec-10 recognizes α2-3- and α2-6-linked sialic acid linkages in the context of *N*-glycans, and that this recognition is enhanced by the presence of Neu5Gc. Overall, the opposing regulation of Siglec-10/-G ligands on GC B cells and follicular T cells suggests a model whereby the loss of *cis* ligands on GC B cells unmasks Siglec-10/-G, while the upregulation of ligands on Tfh cells promotes *trans* interactions (**Figure S20**).

### Proximity labeling to identify binding partners of Siglec-10 on CD4^+^ T cells

We next aimed to identify the specific cell surface proteins that carry the Siglec-10 ligands using an APEX2-based proximity labeling assay. Follicular CD4^+^ T cells were enriched from OVA-immunized mice using magnetic beads (**Figure 7A**) and these isolated cells were sequentially incubated with the Siglec-10-Fc followed by a protein-A-APEX2 fusion protein (**Figure 7B**)(*49, 50*). APEX2 catalyzes the oxidation of biotin-tyramide to highly reactive biotin-phenoxyl radicals, which covalently bind to proteins within 20 nm proximity(*51*). Biotinylated proteins were purified using streptavidin magnetic beads and identified via mass spectrometry-based proteomics analysis. To confirm the activity of APEX2 enzyme, biotinylated surface proteins were detected using an anti-biotin antibody and analyzed by flow cytometry and imaging flow cytometry (**Figure 7C,D**). Biotin staining only showed an increase on Siglec-10-Fc-labeled follicular CD4^+^ T cells, and not with Siglec-10R-Fc, by flow cytometric and imaging flow cytometry.

**Figure 7.**
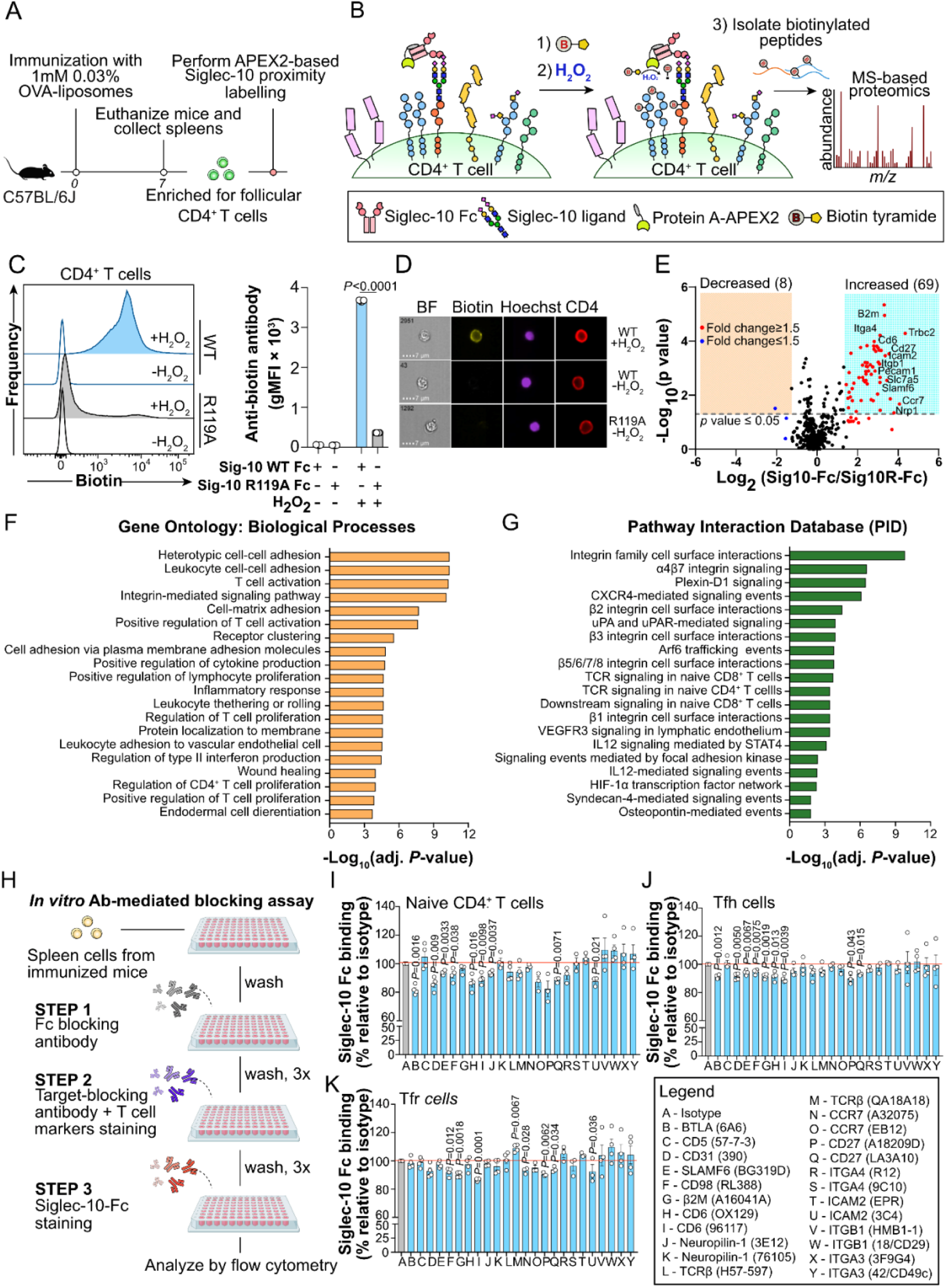
APEX2-based proximity labeling assay detected putative binding partners for Siglec-10 on follicular CD4^+^ T cells. (**A**) Schematic representation of the experimental timeline for follicular CD4^+^ T cell isolation and the APEX2-based proximity labeling assay performed using spleen cells from WT. (**B**) Diagram illustrating the principle of the APEX2 labeling strategy, where the protein A-APEX2 fusion protein links to Siglec-10-Fc and biotinylates adjacent Siglec-10 partners. (**C, D**) Flow cytometry (**C**) and imaging flow cytometry (**D**) quantification of biotinylation on CD4^+^ T cells stained with either Siglec-10-Fc or the non-binding Siglec-10R-Fc control, assayed with or without H_2_O_2_. (**E**) Volcano plot summarizing the MS-based proteomic analysis comparing Siglec-10-Fc binding to the IgG Fc control. The y-axis (-log_10_, *p*-value) represents statistical significance, and the x-axis (log_2_, Fold Change) represents the degree of enrichment for potential Siglec-10-Fc partners. (**F, G**) Gene Ontology (GO) enrichment analysis (**F**) and Pathway Interaction Database analysis (**G**) of the significantly enriched proteins identified in the proteomic analysis. (**H**) Schematic diagram illustrating the experimental design of the *in vitro* antibody-mediated blocking assay. (**I-K**) Quantification of Siglec-10-Fc staining after blocking with antibodies targeting specific T cell surface proteins on naive CD4^+^ T cells (**I**), Tfh cells (**J**), and Tfr cells (**K**). Data are normalized to the 100% anti-Rat-IgG-isotype control group. Data are presented as mean ± SEM. Statistical analysis was performed using the one sample t test.

Proteomic analysis of the biotinylated proteins successfully identified 69 proteins significantly enriched as putative Siglec-10-Fc binding partners, including the following proteins as the top 15 highest scoring hits: *Trbc2, Ccr7, Nrp1, Gmfg, TVB4, Cd27, Atp1b3, Icam2, B2m, Sell, Cd6, Scamp3, Slc7a5, Itga4,* and *Slamf6* (**Figure 7E; Table S2**). GO enrichment analysis revealed that the 69 enriched proteins are associated with cell-cell adhesion, T cell activation, and the integrin-mediated signaling pathway (**Figure 7F**). Likewise, Pathway Interaction Database (PID) identified several proteins from the dataset to be involved in integrin interactions and signaling, T cell activation, and IL-12 signaling (*52*) (**Figure 7G**). Taken together, result from the APEX2-based proximity labeling highlights the potential interaction of Siglec-10 and key immunoregulatory proteins on follicular CD4^+^ T cells, specifically identifying a network of receptors critically involved in T cell activation.

We next sought to validate which of these identified candidates function as Siglec-10 ligand carriers by measuring the reduction of Siglec-10-Fc binding in the presence of specific blocking antibodies. To explore the ability of specific cell surface proteins to carry the Siglec-10 ligands, we used an *in vitro* blocking assay to test 15 of the highest-scoring Siglec-10-Fc partners identified by the proximity labeling, as well as other proteins of interest on the T cell surface (**Figure 7H**). This strategy relied on the ability of specific blocking antibodies to bind and sterically hinder Siglec-10-Fc binding. First, spleen cells were treated with an Fc blocking antibody to inhibit endogenous Fc receptor binding, followed by incubation with the specific blocking antibodies and Tfh cell markers, and staining with Siglec-10-Fc. The blocking assay, which normalized Siglec10-Fc staining to the IgG isotype control group, revealed a significant decrease in Siglec-10-Fc binding when using the blocking antibodies against BLTA, CD5 CD31, SLAMF6, CD98, CD6, Neuropilin-1, CD27 and ICAM2 on naive CD4^+^ T cells (**Figure 7I**). On Tfh cells, a significant decrease in Siglec-10-Fc binding was observed with BLTA, CD31, SLAMF6, CD98, CD6, and CD27 antibodies (**Figure 7J**). On Tfr cells, a significant decrease was observed using antibodies against CD98, CD6, and CD27 (**Figure 7K**). Notably, antibodies targeting CD98, CD6, and CD27 consistently reduced binding across all three subsets, identifying them as potential universal carriers. The antibody blocking assay supported the APEX2 findings by suggesting that Siglec-10 binds to a network of multiple low-affinity ligands carriers across Tfh and Tfr subsets. Interestingly, the reported Siglec-10 ligand, CD24, did not show up as a hit. As four other studies have recently failed to capture CD24 as a Siglec-10 ligand(*18, 44, 52, 53*), we followed up these results by overexpressing CD24 in U937 cells, which did not enhance Siglec-10-Fc binding (**Figure S21**).

## DISCUSSION

The current model of B cell positive selection in the GC requires signals derived from the BCR through engagement of antigens presented by follicular dendritic cells (Signal 1), followed by T cell help via physical contact with cognate Tfh cells (Signal 2). This stepwise process results in sustained survival and expansion of positively selected GC B cells in the DZ compartment without the need of constant stimulation(*4, 54–56*). Mechanistically, this enhanced proliferative capacity is due to the synergistic of signals derived from BCR activation, CD40/40L, and Il-21R/IL-21 interactions(*57–61*). Engagement of these receptors activate critical downstream mediators, including NF-κB and STAT3, which ultimately converge to upregulate the Myc and mTORC pathways, triggering sustained survival and cell cycle entry(*57, 60, 61*). This process often results in fixed proliferation cycles, protecting the host from overt B cell expansion that could potentially lead to autoimmunity or cancer(*62–65*). How the GC inherently tunes B cell selection to prevent aberrant B cell proliferation while maintaining robust SHM and selection of affinity matured antibodies has yet to be fully elucidated. In this study, we identified Siglec-G, a mouse ortholog of Siglec-10 in humans, as an intrinsic B cell checkpoint receptor that restrains the strength of the signals delivered to GC B cells during positive selection.

Previous work suggested that the inhibitory function of Siglec-G is largely restricted to B-1 cells(*14, 16, 23, 66–68*). However, spontaneous GC hyperplasia observed in aged Siglec-G^KO^ mice implied that Siglec-G may regulate a specific B-2 lineage cells, such as GC B cells, either by dampening BCR activation or through other unknown mechanisms. To accurately test the function of Siglec-G on B cells and minimize the impact of Siglec-G depletion during B cell development, we developed a Siglec-G floxed mouse line. Crossing it with *CD20*^CreERT2^ mouse(*69*) produced a temporal and B cell-specific deletion of Siglec-G. Depletion of Siglec-G on B cells post-immunization triggered an expanded GC, augmented PC output, and enhanced antibody responses, establishing Siglec-G as a negative regulator of the GC reaction. Nevertheless, the number of antigen-specific MBCs remained unaffected in Siglec-G-deficient mice. Current models establish that PC selection requires a strong Signal 1 and 2: high-affinity BCR engagement combined with robust T cell help from strong CD40 and IL-21 signaling, respectively, to drive the expression of key PC differentiation transcription factors, IRF4 and Blimp-1(*70–75*). Hence, the skewed differentiation of Siglec-G-deficient GC B cells towards PCs argues that Siglec-G is involved in dampening the strong activating signals coming from either the BCR or B-T cell interactions.

Consistent with foundational studies on Siglec-G(*16, 76*), our analysis confirmed that Siglec-G is dispensable in regulating BCR activation in B-2 lineage cells, such as follicular and GC B cells. Kinetics of BCR-antigen internalization and antigen degradation, which are both downstream events following BCR activation, were also intact in GC B cells lacking Siglec-G. Despite Siglec-G being structurally related to CD22, these two Siglecs on B cells appear to play opposite roles in the GC. While both receptors contain ITIMs capable of recruiting SHP-1/2 when phosphorylated and have overlapping glycan recognition(*77, 78*), deficiency for CD22 on B cells in mice results in blunted GC and antibody responses following a T-dependent (TD) antigen immunization. This was pinpointed to be partly due to BCR hyperactivation, leading to increase activation-dependent cell death of GC B cells(*10*). However, mice that lack Siglec-G on B cells displayed an overall enhancement of GC reaction. Our *ex vivo* BCR stimulation assay using anti-Igκ as a surrogate soluble antigen showed comparable BCR function between WT and Siglec-G^KO^ GC B cells. However, this does not intrinsically imply that the inhibitory function of Siglec-G is impaired on B-2 cells because forced recruitment of Siglec-G close to the BCR complex via high affinity Siglec-G ligands effectively dampened BCR activation and TD antibody responses(*20*). We speculate that the low cell surface levels and distinct spatial organization of Siglec-G compared to CD22 may have contributed to its inability to control BCR signaling in B-2 cells, such as the GC B cells(*67, 79, 80*). Instead, our findings support a model wherein Siglec-G enacts its inhibitory function by restricting the strength of T cell help following B-T cell interactions in the GC, effectively limiting the induction of Myc and mTORC pathways required for GC B cell proliferation.

The ability of *trans* glycan ligands to drive biological activities of Siglecs is an emerging concept(*81–86*). Specifically, studies demonstrate that the interaction between *trans* ligands and Siglecs can recruit and concentrate Siglecs at the immunological synapse(*87–90*). Sequestration of Siglec-G by *trans* ligands on Tfh cells is orchestrated by the remodeling of sialic acid landscape within the GC. In this study, we observed downregulation of Siglec-G/10 ligands on GC B cells in both mice and humans. This reduced ligand expression unmasks Siglec-G/10, making it more accessible for binding with *trans* glycan ligands present on other cells. Our data suggest that unmasking of Siglec-G on mouse GC B cells likely involves a switch from Neu5Gc to Neu5Ac sialic acid, driven by the downregulation of CMAH enzyme. However, humans do not make Neu5Gc due to the inactivating mutation found in the *CMAH* gene(*91*). Thereby, the mechanism resulting in the loss of Siglec-10 ligands on human GC B cells is yet to be determined. It is worth noting that the same scenario was previously observed for CD22 on GC B cells. Both mice and humans show downregulation of CD22 ligands on GC B cells albeit the mechanisms involved in the unmasking are distinct between species(*10, 12, 92, 93*). Interestingly, in contrast to the downregulation of Siglec-G/10 ligands on GC B cells, we observed an upregulation of ligands on Tfh compared to their naive counterparts. Our results suggest that this upregulation is largely driven by the increased levels of α2-6 linked sialic acids glycans on Tfh cells, as evident by the enhanced SNA lectin staining and higher transcript expression of *St6gal1* compared to non-follicular CD4^+^ T cells. Altogether, these glycosylation changes on GC B cells and Tfh cells likely induce an efficient recruitment of Siglec-G at the immunological synapse formed at the interface between these two cells.

Once recruited at the synapse, there are several mechanistic possibilities on how Siglec-G could limit the signal strength delivered via B-T cell interactions. One possibility is that Siglec-G could sequester inhibitory co-receptors to the synapse of cognate Tfh cells; thereby, placing these receptors in close proximity to the TCR complex and effectively dampening TCR-mediated signaling. Using an APEX2-based proximity labeling assay, we identified several inhibitory co-receptors on CD4^+^ T cells that potentially bind with Siglec-G/10. These receptors include CD6, BTLA, and SLAMF6, which have been shown to inhibit Tfh activation and function (*94–99*). Consistent with this concept, OT-II Tfh cells co-cultured with OVA_323-339_ pulsed Siglec-G deficient B cells induced more CD40L surface mobilization than their WT B cell counterpart, highlighting the role of Siglec-G in tuning Tfh stimulation potentially by sequestering inhibitory co-receptors at the synapse. Another possible mechanism on how Siglec-G could modulate T cell help through *trans* glycan interactions is via recruitment of Siglec-G at the synapse by ligand-carrying counter receptors on Tfh cells. This recruitment brings Siglec-G closer to B cell stimulatory co-receptors such as CD40, which is upstream of Myc and mTORC pathways(*4, 7*). While there is no direct evidence linking Siglec and CD40 inhibition, signals downstream of CD40 can be regulated indirectly by SHP-1(*100*). Specifically, SHP-1 has been shown to interact with TRAF3, a known negative regulator of CD40(*100–102*), and that this interaction promotes TRAF3 dephosphorylation which inhibits protein ubiquitination and degradation(*100*). Further experiments are needed to fully elucidate whether the latter mechanism also contributes to restraining the GC reaction.

Besides expansion of GC B cells and plasma cells, Siglec-G deficiency on B cells also impacted the follicular CD4^+^ T cell compartment, with an overall increase in the abundance of both Tfh and Tfr inside the GC. Given that loss of Siglec-G did not alter B-T cell conjugation *in vitro*, it is highly likely that confinement of more Tfh cells within the GC of Siglec-G^BKO^ mice is mainly due to enhanced T cell activation following contact between GC B cells and Tfh cells. The increased amplitude of Tfh activation in Siglec-G^BKO^ mice, as observed by high expression of activation-induced markers CD40L, CD69, and ICOS, may result in better Tfh retention in the GC than their control counterparts. Congruent with this idea, T cells lacking the expression of SAP, an adaptor protein that promotes TCR signaling, displayed limited recruitment and retention of Tfh cells in the GC(*103*). Moreover, the maintenance of Tfh program, including receptors required for GC retention such as PD-1 and CXCR5 (*104–106*), relies on the robust expression of BCL6, which by itself depends partly on TCR signaling and NFAT transcription factors(*107, 108*). Likewise, the increase in frequency and localization of Tfr cells within the GC of Siglec-G^BKO^ mice may be caused by similar cues that governs Tfh cell maintenance and retention. It is also feasible that a subset of this expanded Tfr compartment arose from Tfh cells that upregulated Foxp3 transcription factor and acquired Tfr-like phenotypes(*109*). However, a detailed clonal analysis is required to confirm this hypothesis. Given the multifaceted nature of Tfr cells, it is not surprising that increased Tfr cell numbers were not sufficient to broadly suppress the GC in immunized Siglec-G^BKO^ mice. In fact, we speculate that Tfr cells may have promoted GC reaction to some extent in our Siglec-G deficient model. Our scRNA-seq analysis revealed that a significant subset Tfr cells from Siglec-G^BKO^ mice upregulated the transcript for *Il10* compared to control. This Tfr-derived cytokine has been shown to support the GC response as mice lacking IL-10 in Tfr cells displayed significant reduction in GC B cells and plasmablasts following LCMV infection (*110, 111*). Additionally, our sequencing data identified a reduction in the proportion of *Eomes^+^* Tfh cells in Siglec-G^BKO^ compared to control mice. This subset of *Foxp3^-^*follicular T cells has recently emerged as a negative regulator of humoral immunity by controlling the Tfh and GC B cells (**REFS**). Collectively, this skewed balance of different follicular T cell subsets highlights the involvement of Siglec-G on B cells in shaping the functionality Tfh and Tfr cells as well as modulating their retention inside the GC.

These findings may offer important insights into pathogenesis of human DLBCL. Building on previous report that the loss of Siglec-G to the development of B cell malignancies(*23*), our studies also further characterize this pathology by analyzing the aberrant expansion of both GC B and Tfh cell compartments and how they correlate with shortened lifespans. Crucially, a B cell-intrinsic malignancy was evidenced by the ability of adoptive transferred Siglec-G-deficient B cells to cause death in immunodeficient Rag1^-/-^ hosts. The clinical significance of these findings are that *SIGLEC10* is downregulated in DLBCL tumors compared to healthy B cells. Moreover, a strong correlation between low *SIGLEC10* expression and poor patient survival suggests that Siglec-10 functions as a vital tumor suppressor, analogous to Siglec-G in mice(*23*). Therefore, within the highly proliferative environment of the GC, loss of Siglec-10/G serves as a critical immune checkpoint that restrains lymphomagenesis.

## CONCLUSION

In conclusion, our work establishes Siglec-G/10 as a glycan-dependent molecular fine tuner of B-T cell interaction. By integrating cell signals, glycan remodeling, and specific protein-protein interactions, Siglec-G ensures that the germinal center reaction produces a focused and high-affinity antibody response while preventing the dysfunctional germinal center response that leads to autoimmunity or cancer. If the receptor lost or its glycan ligands are altered, B cells can escape the proliferative constraints of the checkpoint, leading to lymphomagenesis. Therefore, therapeutic strategies should focus on inhibiting the unconstrained downstream survival signals in Siglec 10 deficient B cell lymphoma. Targeting these hyperactive signaling pathways represents an approach for treating aggressive lymphomas.

## MATERIALS AND METHODS

### Mice

The following mice were used in this study: WT C57BL/6J (#000664), B6-CD45.1 (#002014), *Mb1*^Cre^ (#020505), CD4^Cre^ (#017336), OT-II (#004194), ST6Gal1^flox/flox^ (#006901) and Rag1^KO^ (#002216) were obtained from Jackson Laboratory (Maine, USA). Siglec-G^KO^, *CD20*^CreERT2^, ST3Gal1^KO^, and ST3Gal4^KO^ mice were provided to us by Dr. Lars Nitschke (FAU Erlangen-Nürnberg, Erlangen, Germany), Dr. Mark Shlomchik (University of Pittsburgh, Pennsylvania, USA), Dr.Jamey Marth (Sanford Burnham Prebys Medical Discovery Institute, CA, USA).

Floxed *Siglecg* mice (C57BL/6-Siglecg^tm1c(EUCOMM)Tcp^) were generated by The Centre for Phenogenomics (Toronto, Canada). *Siglecg*^tm1a(EuCOMM)Tcp^ allele was excised by microinjecting 100 ng/μL of FLP recombinase mRNA (Miltenyi Biotec Inc.) into the cytoplasm of *in vitro* fertilization (IVF)-derived zygotes. Successful conversion of tm1a allele into tm1c allele of mouse *Siglecg* was detected by PCR through the presence of distal *loxP* site (primers: 5’-CGCAACGCAATTAATGATAACTTCG-3’ and 5’-GGACTTTGAGACCTGAATTTAATCCG-3’; band size: 355 bp) and absence of WT allele (primers: 5’-GGGAGATGAGTAGGCTGATAAAGAG-3’ and 5’-GAGACAGACATAAGGAAAGGGAGAG-3’; band sizes: 1083 and 876 bps). Mice were subsequently backcrossed in-house onto the C57BL/6J background. Homozygous floxed *Siglecg* mice (*Siglecg*^f/f^) were then crossed with either *Mb1*^Cre^ or *CD20*^CreERT2^ to generate *Siglecg* conditional knockout models, *Mb1*^Cre^×*Siglecg*^f/f^ and *CD20*^CreERT2^×*Siglecg*^f/f^, respectively, that delete Siglec-G specifically on B cells.

*Rosa26^lsl^-Cmah* mice were generated following a previously described protocol(*10*). To confirm the genotype of *Rosa26^lsl^-Cmah* mice, PCR was performed using extracted DNA from ear notch samples. The WT Rosa26 locus was identified using 50-GGAGCGGGAGAAATGGATATG-30 as forward primer and 50-AAAGTCGCTCTGAGTTGTT AT-30 as reverse primer (band size: 600 bp). On the other hand, the *Rosa26^ls^*^l^-*Cmah* gene was determined using 50-ATTCTAGTTGTGGTTTGTCC-30 and 50-ATGGTGCTCACGTCTAACTTC C-30 primers (band size: 370 bp).

All mice were bred and maintained in a specific pathogen-free facility of the Health Sciences Laboratory Animal Services (HSLAS) at the University of Alberta. Animal procedures described in this study were approved by the Health Sciences Animal Care and Use Committee of the University of Alberta, in accordance with the guidelines set by Canadian Council on Animal Care. Both male and female mice between 2 months to 4 months of age were used for all experiments.

### Adoptive transfer of immune cells

Chicken ovalbumin (OVA)-specific CD4^+^ T cells were isolated from naive OT-II transgenic mice. Spleens were dissected from euthanized mice and mashed through a 40 μm cell strainer. Red blood cells were depleted using 1X ACK lysis buffer and the resulting cell suspension was resuspended in 1X HBSS (Gibco^TM^) supplemented with 1% bovine serum albumin (BSA) and 2 mM EDTA. OT-II T cells were isolated by negative selection using a commercially available mouse CD4^+^ T cell isolation kit (Miltenyi Biotec). Isolated cells were then resuspended in sterile 1X PBS (Gibco^TM^) and 2 million cells were injected intravenously into naive C57BL/6J or B6-CD45.1 mice. Age and sex of both donor and recipient mice were matched in all adoptive transfer experiments performed in this study.

For aged B cell transplantation, B cells from 20-24 months old WT C57BL/6J and Siglec-G^KO^ mice were isolated from spleens and lymph nodes (inguinal, brachial, axillary, and cervical) using a pan-mouse B cell isolation kit (Miltenyi Biotec). Approximately 10 million cells were adoptively transferred into 8-10 weeks old and sex matched *Rag1*^KO^ mice via intraperitoneal injection. Recipient mice were monitored daily for survival and morbidities.

### Cell lines

Flp^TM^-In Chinese hamster ovary (CHO) cells (Invitrogen^TM^) were cultured in DMEM/F-12 media (Gibco^TM^) supplemented with 5% heat-inactivated fetal bovine serum (FBS, Gibco^TM^), 10 mM HEPES, and 100 U/mL penicillin and 100 μg/mL streptomycin (Gibco^TM^). Human embryonic kidney (HEK) 293T cells (ATCC) were maintained in DMEM media (Gibco^TM^) with 10% heat-inactivated FBS and 100 U/mL penicillin and 100 μg/mL streptomycin. WT, *MGAT1*^KO^, *COSMC*^KO^, and *MGAT1*^KO^×*COSMC*^KO^ U937 cell lines were maintained in RPMI media (Gibco^TM^) with 10% heat inactivated FBS and 100 U/mL penicillin and 100 μg/mL streptomycin. All cell lines were grown at 37°C, in a 5% CO_2_ incubator.

### Generation of CD24 over-expressing HEK293T and U937 cells

cDNA encoding CD24 was inserted into the RP172 lentiviral vector as described previously(*112*). To make lentivirus, the RP172-CD24 transfer plasmid, and RP18 and RP19 helper plasmids were combined with TransIT-LT1 Transfection Reagent (Mirus Bio) and transfected into HEK293T cells and cultured for 3 days. The cultured medium was then harvested and mixed with Lenti-X Concentrator (Takara Bio) in a 3:1 ratio and incubated for 1 hour at 4 °C. After centrifugation at 1500 rcf. at 4 °C for 45 min, pellets containing virus were resuspended in PBS, aliquoted, and stored at −80 °C. To generate the CD24 overexpressing U937 and HEK293T cell lines, lentivirus was added to 200,000 cells and cultured for 3 days. After 3 days, the transduced cells were cultured with RPMI medium (U937) or DMEM (HEK293T) containing penicillin/streptomycin, 10% fetal bovine serum, and 300 μg/mL Zeocin for selection of positively transduced cells. Selection was carried out for 7 days, changing media and adding fresh Zeocin every 2 days. To ensure cultures were fully selected, cells were analyzed on an LSRFortessa X-20 flow cytometer (Becton Dickenson) for mAmatrine expression.

### Antigen-lipid conjugation

OVA protein (Millipore Sigma) was conjugated to pegylated 1,2-Distearoyl-sn-glycero-3-phosphoethanolamine (PEG-DSPE) using maleimide chemistry. First, reactive thiol groups were introduced to OVA using a heterobifunctional crosslinker *N*-Succinimidyl 3-(2-pyridyldithio)-propionate (SPDP; Thermo Scientific^TM^). OVA (in 1X PBS) and SPDP (in DMSO) were mixed in a 1:2.5 molar ratio, respectively, and the reaction mixture was incubated for 1 hour at room temperature with gentle agitation. Protein was then desalted by passing it through a Sephadex-50 column equilibrated with 1X PBS (pH = 6.5). Flow through fractions containing proteins were combined and treated with 25 mM DTT for 10 minutes at room temperature. The release of thiol 2-pyridyl was monitored at absorbance 343 nm, using a NanoDrop^TM^ (Thermo Scientific^TM^). After desalting, the thiol-derivatized OVA (in 1X PBS; pH = 6.5) was reacted with maleimide PEG_2000_-DSPE (in DMSO; Avanti Polar Lipids) in a 1:5 molar ratio. The reaction mixture was incubated overnight at room temperature with constant stirring and in the presence of N_2_ gas. PEG-DSPE-modified OVA was separated from the unmodified OVA using a Sephadex-100 column and stored in 4°C prior to use. SDS-PAGE was used to verify the purity of OVA-lipid conjugation.

Lipid conjugated NP-OVA was produced by adding 20 molar equivalents of 4-Hydroxy-3-nitrophenyl hapten (NP)-ε-Aminocaproyl-OSu (in DMF; LGC Biosearch Technologies) for every 1 molar equivalent of OVA-PEG_2000_-DSPE (in 3% NaHCO_3_). The reaction was performed for 2 hours at room temperature, with constant agitation. The solution was then dialyzed overnight in 1X PBS (pH = 7.4) using a SnakeSkin^TM^ dialysis tubing (3500 MWCO; Thermo Scientific^TM^) to remove excess NP-ε-Aminocaproyl-OSu. Dialyzed NP-OVA-PEG_2000_-DSPE was stored in 4°C prior to use.

### Antigen-decorated liposome preparation

Working solutions of 1,2-Distearoyl-sn-glycero-3-phosphocholine (DSPC; Avanti Polar Lipids), Cholesterol (Millipore Sigma), and PEG_2000_-1,2-Distearoyl-sn-glycero-3-phosphoethanolamine (PEG_2000_-DSPE; Avanti Polar Lipids) were prepared by solvating the lipids in chloroform. An appropriate volume of each lipid solution was transferred into a glass test tube to reach 57:38:4.7 mol % of DSPC, Cholesterol, and PEG_2000_-DSPE, respectively. Chloroform was removed by flushing N_2_ gas to yield a thin film of lipids. Dry lipids were resuspended in 100 μL DMSO and stored in -80°C freezer. Once completely frozen, DMSO was removed via lyophilization. Lyophilized lipids were then stored in -80°C prior to use. To prepare for OVA- and NP-OVA-decorated liposomes, an appropriate amount of OVA-PEG_2000_-DSPE or NP-OVA-PEG_2000_-DSPE was added to the lyophilized lipids to achieve a final concentration of 0.03 mol %. The lipid mixture was immediately hydrated in sterile 1X PBS and sonicated for 5-10 minutes. After sonication, antigen-decorated liposomes were extruded at least 20 times through an 800 nm and then a 100 nm polycarbonate membrane filters (Whatman), using a hand-held mini-extruder device (Avanti Polar Lipids). Extruded liposomes were purified over Sepharose CL-4B column and then reconstituted in sterile 1X PBS to a final concentration of 1 mM.

### Mouse immunization and treatments

GCs were induced in mice by immunization with 200 μL of freshly prepared 1 mM NP-OVA-decorated liposomes via tail vein injection. C57BL/6J or B6-CD45.1 mice adoptively transferred with OT-II T cells were immunized with 200 μL of 1 mM OVA-decorated liposomes via tail vein injection, 1 day after cell transfer. Where indicated, mice were treated with 50 μg/mouse of anti-Dec205^OVA^ or 1X PBS on day 13 post-immunization to acutely induce B-T cell interaction. For Siglec-G antibody-based blocking experiment, mice were treated with 50 μg/mouse of either Rat anti-Siglec-G monoclonal antibody (clone: 4A6) or Rat IgG2a isotype control (clone: RTK2758; BioLegend) on days 7 and 9 post-immunization.

For temporal depletion of Siglec-G, *CD20*^CreERT2^×*Siglecg*^f/f^ and control (*CD20*^CreERT2^) mice were treated with 100 μL of 10 mg/mL Tamoxifen (Combi-Blocks) in corn oil (Millipore Sigma) via intraperitoneal injection on days 6, 8, and 10 post-immunization.

### Flow cytometry

Splenocytes were harvested from mouse spleens for flow cytometric analysis. Spleens were dissociated in complete RPMI media (Gibco^TM^) by mashing the tissue through a 40-μm cell strainer, followed by RBC depletion using 1 mL of 1X ACK lysis buffer for 1 min on ice. Cells were spun down, washed, and resuspended in appropriate volume of 1X HBSS supplemented with 2% FBS (flow buffer). Prior to staining, cells were incubated with 1 μg/mL on anti-mouse CD16/32 antibody (clone: S17011E; BioLegend) for 10 min on ice to block for non-specific immunoglobulin binding to mouse Fc receptors. After washing, cells were stained with commercially available fluorochrome-conjugated monoclonal antibodies (BioLegend and BD Bioscience) against mouse surface receptors CD19 (clone: 6D5), B220 (clone: RA3-6B2), IgD (clone: 11-26c.2a), CD38 (clone: 90), CD95 (clone: Jo2), GL7 glycan epitope (clone: GL7), CD86 (clone: GL1), CXCR4 (clone: L276F12), Siglec-G (clone: SH1), CD138 (clone: 281-2), TAC-I (clone: 8F12), EpCAM (clone: G8.8), CXCR3 (clone: CXCR3-173), CD80 (clone: 16-10A1), PD-L2 (clone: TY25), CCR6 (clone: 29-2L17), CD4 (clone: GK1.5), CD3 (clone: 17A2), CD62L (clone: MEL-14), CD44 (clone: IM7), CXCR5 (clone: L138D7), PD-1 (clone: 29F.1A12), CD90.2 (clone: 30-H12), CD69 (clone: H1.2F3), ICOS (clone: 7E.17G9), CD45 (clone: I3/2.3), CD45.1 (clone: A20), CD45.2 (clone: 104), CD40L (clone: MR1), GITR (clone: DTA-1), OX40 (clone: OX-86), 4-1BB (clone: 17B5), CD120b (clone: TR75-89), and CD73 (clone: TY1/11.8) in flow buffer for 30 min at 4°C. For detection of antigen-specific B and T cells, splenocytes were incubated with either NP_21_-APC (in-house) and NP_28_-PE (LGC Biosearch) for NP-specific B cells or OVA_329-337_-loaded I-Ab tetramer-PE (NIH Tetramer Facility, GA, USA) for OVA-specific CD4^+^ T cell for 30 min at room temperature prior to cell surface receptor staining. Stained cells were washed and resuspended in flow buffer containing 1 μg/mL of 7-AAD for the detection of live/dead cells. An appropriate volume of CountBright counting beads (Invitrogen^TM^) was also added in cell suspension for absolute cell counting. Flow cytometric analysis was performed using BD LSRFortessa X-20 Cell Analyzer equipped with 5 lasers and 20 parameters. Immune cell subsets were detected using the following combination of markers: naive B cells (B220^+^ CD19^+^ IgD^+^), antigen-specific GC B cells (B220^+^ CD19^+^ IgD^-^ Fas^+^ GL7^+^ NP^+^), memory B cells (B220^+^ CD19^+^ IgD^-^ CD38^+^ CD80^+^ PD-L2^+^ NP^+^), plasma cells (CD45^+^ B220^+^ IgD^-^ CD138^+^ TAC-I^+^), Tfh (B220^-^ CD3^+^ CD4^+^ CD44^+^ CD62L^-^ PD-1^hi^ CXCR5^hi^ Foxp3^-^), Tfr (B220^-^ CD3^+^ CD4^+^ CD44^+^ CD62L^-^ PD-1^hi^ CXCR5^hi^ Foxp3^+^). GC B cells in DZ and LZ compartments were gated as CXCR4^hi^ CD86^lo^ and CXCR4^lo^ CD86^hi^, respectively.

For detecting nuclear and phosphorylated intracellular proteins, cells stained with surface markers were fixed with 2% paraformaldehyde (PFA) at room temperature for 5 min. Fixed cells were washed with flow buffer and permeabilized with 1X True-Nuclear^TM^ Transcription Factor permeabilization buffer (BioLegend). Fixed and permeabilized cells were then washed twice with flow buffer followed by overnight staining at 4°C with antibodies against transcription factors Bcl6 (clone: K112-91), Foxp3 (clone: FJK-16s), Nfatc1 (clone: 7A6), and c-Myc (clone: D84C12), as well as phosphorylated proteins ribosomal S6 (clone: D57.2.3E; pS235/236), Akt (clone: D9E; pS473), Akt (clone: D25E6; pT308), and Syk (clone: 65E4; pY352). Following incubation, cells were washed once with 1X True-Nuclear^TM^ Transcription Factor permeabilization buffer prior to flow cytometric analysis.

For identifying cycling and proliferating GC B cells, immunized mice were injected with 2 mg/mouse Bromodeoxyuridine (BrdU; MilliporSigma) intraperitoneally 1 hour prior to euthanasia and their spleens were harvested, depleted for RBCs, and dissociated into single cell suspensions. Cells were incubated with anti-mouse CD16/32 antibodies and stained with appropriate cell surface markers. After incubation, cells were washed with a flow buffer and fixed with BD Cytofix/Cytoperm^TM^ (BD Biosciences) for 15 min on ice. Cells were then washed once with 1X BD Perm/Wash buffer (BD Biosciences) followed by 10 min incubation with BD Cytoperm^TM^ Permeabilization Buffer Plus (BD Biosciences) at 4°C. Cells were washed with 1X BD Perm/Wash buffer and re-fixed BD Cytofix/Cytoperm^TM^ for 10 min at 4°C. After the second round of fixation, cell pellets were resuspended in 1X PBS supplemented with 1 μM Ca^2+^, 1 μM Mg^2+^ ions, and 400 μg/mL DNAse I (Millipore Sigma), and incubated for 1 hour at 37°C. Following incubation, cells were washed with 1X BD Perm/Wash buffer and stained with anti-BrdU antibody (BioLegend) and 7-AAD dye (Invitrogen^TM^) for 30 min. Proliferating GC B cells were gated as BrdU^+^ cells while cell cycle phases were identified as described previously.

### Evaluating the dependency of the sialic acid linkage for the Siglec-10 ligands on B and T cells

Splenocyte preparation and cell staining were conducted using the methodology described above for flow cytometry. For investigating the sialic acid linkage dependency of Siglec10-Fc ligand binding on T cells and B cells by gene-knockout murine model, spleens were harvested from C57BL/6J and sialyltransferase-deficient mice, *ST6Gal1^fl/fl^* x *CD4cre*^-/+^, *mb1cre^-/+^* x *R26^ISI^-Cmah*, *ST3Gal1*^KO^, and *ST3Gal4*^KO^, and processed into cell suspensions. Spleens were macerated in RPMI-1640 medium containing 2% FBS and filtered through a 40 μm mesh. Red blood cells were lysed using ammonium-chloride-potassium (ACK) buffer for 5 minutes at 4°C. Cells were washed and resuspended in HBSS containing 2% FBS. Cells were stained on ice for 30 minutes using fluorophore-conjugated antibodies listed in the key material table.

For Siglec-10-Fc partner blocking assays, cells were also treated with TruStainX to block non-specific Fc receptor binding before the cell marker staining. The cell suspension was incubated on ice for 10 minutes as the first step. Cells were stained with cocktails specifically including an antibody for the target Siglec-10-Fc binding partner from the top hit of the proximity labeling, alongside a negative control (Mouse IgG2b, κ Isotype Ctrl Antibody). After staining, cells were followed by the Siglec-10-Fc pre-complex staining, discussed above. To quantify the blocking efficiency, the Siglec-10-Fc staining intensity for each candidate protein was normalized to the IgG isotype control group (set to 100%). The final presented data for the 15 candidate proteins represents a compilation of results from four independent experiments.

For assessing the sialic acid linkage dependency of Siglec-10-Fc by Neuraminidase, a subset of cells was pre-treated with neuraminidase. Briefly, cells were incubated with either recombinant enzyme Neuraminidase A or Neuraminidase S. The cells were treated for an hour at 37 ^◦^C, and washed to remove the enzyme. Following this treatment, the cells were stained with Siglec10-Fc, and processed for flow cytometry as described above. Siglec-10-Fc, pre-complexed following a previously described protocol(*113*), was incubated with the cell suspension to assess binding. After staining, cells were washed and resuspended in flow buffer supplemented with 1 µg/mL 7-AAD for viability assessment. Flow cytometric analysis was conducted on a BD Fortessa X-20 cytometer. CD4^+^ T and CD8^+^ T cell populations were gated B220^-^, CD3^+^, CD4^+^ and B220-, CD3^+^, CD8^+^ individually.

### Fluorescence Activated Cell Sorting (FACS) of follicular CD4^+^ T cells

Spleens from immunized mice were dissociated in complete RPMI 1640 media supplemented with 5 μg/mL of actinomycin-D (Millipore Sigma). After RBC depletion, splenocytes were incubated with anti-mouse CD16/32 antibodies and stained for cell surface markers B220, CD3, CD4, CD44, PD-1, and CXCR5 in HBSS (Gibco^TM^) supplemented with 2% FBS, 1 mM EDTA, and 5 μg/mL of actinomycin-D (FACS buffer). Cells were then washed and resuspended in FACS buffer with 1 μg/mL 7-AAD. Follicular CD4^+^ T cells were gated and sorted BS FACS Melody (BD Biosciences) as B220^-^CD3^+^CD4^+^CD44^+^PD-1^hi^CXCR5^hi^ live cells. Sorted cells were collected in complete RPMI 1640 media. Cell count was confirmed using an automated cell counter.

### Enzyme-linked immunosorbent assay (ELISA)

To measure serum antibody titers and affinities from immunized mice, blood samples were collected via tail vein on days 14, 21, and 42 post-immunization. Sera were separated from whole blood cells and stored in -80°C freezer prior to use. To assess antibody titer, 96-well ELISA microplates (Greiner Bio-One) were coated overnight with 50 μL 1X PBS containing 10 μg/mL NP_2_-BSA (prepared in-house) or NP_23_-BSA (LGC Biosearch). Plates were washed 3 times with 1X PBS and blocked with 1% bovine serum albumin (Millipore Sigma) in 1X PBS for 1 hour at room temperature. After blocking step, microplates were washed with 3 times with 1X PBS containing 0.05% Tween-20 (PBS-T) and 50 μL of pre-diluted sera (1:50) in 1% BSA/PBS-T were added into wells and serially diluted 3-folds at least 7 times. After 1 hour incubation, microplates were washed 3 times with PBS-T and incubated with Goat anti-mouse IgG1 conjugated with horseradish peroxidase (HRP) (SouthernBiotech) in 1% BSA/PBS-T for another 1 hour. Microplates were then washed 5 times with PBS-T followed by addition of TMB peroxidase substrate (SeraCare) into each well for colour development. Reaction was stopped by adding equal volume of 1 M phosphoric acid. Optical density was measured using a microplate reader set to 450 nm.

### Enzyme-linked immunospot (ELISPOT) assay for detecting antigen-specific PCs

Multiscreen® 96 well HTS filter plates (MAIPS4510; Milllipore Sigma) were soaked in 35% ethanol for 30 seconds, washed with 1X PBS, and coated overnight at 4°C in 1X PBS containing either 5 μg/mL NP_2_-BSA or NP_23_-BSA. Plates were washed with 1X PBS and blocked with complete RPMI 1640 for 2 hours at room temperature. A specified number of splenocytes from immunized mice were plated and incubated overnight at 37°C in a CO_2_ incubator. Plates were washed with 1X PBS containing 0.01% Tween-20 before incubation for 3 hours with either anti-mouse IgG-HRP (SouthernBiotech) or anti-mouse IgM-HRP (SouthernBiotech). Plates were washed and spots were developed using ELISPOT TMB substrate (Mabtech). Spots were recorded and counted using S6 Immunospot Analyzer (CTL Analyzers, LLC).

### Ig VH 186.2 DNA sequencing and mutation analysis

For VH186.2 sequencing, 20,000-30,000 NP^+^ GC B cells were sorted from spleens of NP-OVA liposome-immunized control and Siglec-G^BKO^ mice. Genomic DNA was extracted using QIAamp DNA micro kit (Qiagen) and stored in - 80 °C prior to use. Nested PCR was used to amplify the VH186.2 region using the following primers: CATGGGATGGAGCTGTATCATGC (F primer) and CTCACAAGAGTCCGATAGACCCTG (R primer) for the 1^st^ PCR reaction, and GGTGACAATGACATCCACTTTGC (F primer) and GACTGTGAGAGTGGTGCCTTG (R primer) for the 2^nd^ PCR reaction (*98*). The final PCR products were inserted into a plasmid using Zero Blunt™ TOPO™ PCR Cloning Kit (Invitrogen^TM^) and transformed into chemically competent DH5α *E. coli* (NEB Biolabs). Plasmids were extracted from *E. coli* clones using a GeneJET plasmid miniprep kit (Thermo Fisher) and DNA inserts were sequenced using ABI 3730 Sanger DNA Sequencer. Mutational analysis of the IgH V region was performed using the IMGT/V-QUEST tool (www.imgt.org).

### Ex vivo BCR internalization and antigen processing assay

Splenocytes from immunized control and Siglec-G^BKO^ mice were incubated with 1 µg/mL of goat F(ab’)_2_ anti-mouse Igκ-biotin (SouthernBiotech) for 30 minutes at 4°C. Cells were washed 3 times with 0.1% BSA supplemented 1X HBSS and resuspended in pre-warmed complete RPMI 1640 media. Cells were then incubated in a 5% CO_2_ incubator at 37°C. At different timepoints, cells were fixed in 1% paraformaldehyde (PFA) for 10 minutes at room temperature to stop BCR internalization. Fixed cells were washed 3 times with 0.1% BSA in 1X HBSS followed by incubation of GC B cell surface marker antibodies and fluorochrome-conjugated streptavidin. Changes in the levels of anti-mouse Igκ-biotin on fixed naive and GC B cells were monitored by flow cytometer.

To test for antigen degradation post-internalization, splenocytes from immunized mice were stained with 1 µg/mL of goat F(ab’)_2_ anti-mouse Igκ-biotin (SouthernBiotech) followed by 1 µg/mL of streptavidin (BioLegend) at 4 °C. Cells were washed 3 times with 1X HBSS containing 0.1% BSA and incubated with 100 nM of biotinylated antigen degradation sensor for 20 minutes at 4°C (*33*). The degradation sensor is comprised of a fluorescent DNA strand sequence: 5 ′ - Atto647N- TCCGGCTGCCTCGCTGCCGTCGCCA-3 ′ -biotin and a quencher DNA strand sequence: 5 ′ -TGGCGACGGCAGCGAGGCAGCCGGA-3′′-Iowa Black RQ. Stained cells were washed 3 times and resuspended in pre-warmed complete RPMI 1640 media. After each timepoint, cells were fixed in 1% paraformaldehyde for 10 minutes at RT. Fixed cells were then washed with 1X HBSS containing 0.1% BSA and stained with GC B cell surface markers. Flow cytometry was used to evaluate changes in the fluorescence of the antigen sensor in GC B cells. Increased Atto647N signal signified increased antigen degradation sensor.

### Intracellular Ca2^+^ mobilization

Around 10 million splenocytes from immunized control and Siglec-G^BKO^ mice were resuspended in 1X HBSS (Ca^2+^/Mg^2+^ free) supplemented with 1% FBS and 1 μM Indo-1 AM (Invitrogen^TM^) and incubated at 37°C for 30 minutes. Cells were washed with 5-fold volume of assay buffer (1% FBS in X HBSS) and centrifuged at 300 rcf for 5 minutes at RT. Pelleted cells were then stained with PNA lectin and antibodies against mouse B220 and CD38 for 20 minutes at 4 C. Cells were washed and resuspended in assay buffer. A 0.5 mL containing 1 million cells were incubated for 5 minutes at 37°C prior to BCR activation and flow cytometry acquisition. After a baseline Indo-1 AM fluorescence was achieved (90 seconds), splenocytes were stimulated with either vehicle (1% FBS in HBSS) or 10 μg/mL goat F(ab’)_2_ anti-mouse Igκ (SouthernBiotech). Indo-1 AM fluorescence ratio (UV379/UV515) was monitored by Fortessa X-20 flow cytometer for 5 minutes. Sample temperature was maintained at 37°C throughout the acquisition. Ca^2+^ mobilization in cells were analyzed using the kinetics function in FlowJo v9.

### Immunofluorescence imaging of GC clusters

Immunized mice were euthanized, and spleens were harvested and immediately fixed in 2% PFA for 5 hours at 4°C. After tissue fixation, spleens were transferred to tubes containing 30% sucrose (MilliporeSigma) in 1X PBS solution and incubated at 4°C overnight. Spleens were then embedded in Cryomatrix^TM^ resin (Epredia^TM^), frozen in a pre-cooled bath of 2-methylbutane chilled in dry ice and stored in -80°C freezer prior to sectioning. Spleen cryosections (9 μm) were blocked in 1X HBSS supplemented with 5% FBS for 2 hours and washed 3 times with PBS-T. To block endogenous biotin, tissue sections were treated Ready probes Streptavidin/Biotin blocking solution (Invitrogen^TM^) following the kit’s recommended protocol. For imaging GC clusters, spleen sections were stained with biotinylated Peanut agglutinin (PNA) lectin (Vector Labs), unlabeled Rat IgG2a anti-mouse CD35 (clone: 8C12; BD Biosciences), and unlabeled Syrian hamster anti-mouse CD3e (clone: 500A2; BD Biosciences) overnight at 4°C. Following incubation, sections were washed 5 times with PBS-T and incubated with AF647 anti-Rat IgG2a (clone: MRG2a-83; BioLegend), AF568 polyclonal anti-Syrian hamster IgG (Invitrogen^TM^), and AF405 streptavidin (Invitrogen^TM^) for 2 hours at room temperature. On the other hand, for enumerating Foxp3^+^ T cells in the GC, spleen sections were stained with AF488 anti-mouse Foxp3 (clone: FJK-16s; Invitrogen^TM^), AF647 anti-mouse IgD (clone: 11-26c.2a; BioLegend), Biotin anti-mouse CD35 (clone: 8C12; BD Biosciences), and unlabeled anti-mouse CD3e overnight at 4°C. After incubation, sections were washed 5 times with PBS-T and stained with AF405 streptavidin (Invitrogen^TM^) and AF568 polyclonal anti-Syrian hamster IgG (Invitrogen^TM^) for 2 hours at room temperature. Stained tissue sections were then washed 5 times with PBS-T and mounted with Fluoro-Gel (EMS). Imaging of whole splenic tissue sections to analyze GC clusters were performed on a Zeiss Axio Observer microscope using a 20X magnification objective. To detect for Foxp3^+^ T cells in the GC, tile scan images of the regions of interest within the tissue section were acquired on the Zeiss LSM 700 laser scanning confocal microscope equipped with Axiocam 702 mono camera and 3 objectives. Individual GC clusters were imaged using 63X magnification oil immersion objective (N.A. 1.4). Images were processed using ZEN 2.6 Blue edition software (Zeiss).

### Single cell RNA sequencing and analysis

Approximately 50,000 follicular CD4^+^ T cells (PD-1^hi^CXCR5^hi^CD4^+^CD3^+^B220^-^7AAD^-^ cells) were sorted by FACS from pooled splenocytes of either immunized control or Siglec-G^BKO^ mice. Sorted cells with >95% viability was resuspended in 1× PBS to achieve a cell suspension containing 1000 cells/mL. cDNA libraries were prepared by UAlberta Advanced Cell Exploration Core. Briefly, barcoded gel beads-cell emulsion was generated by 10X Chromium Controller, and the cDNA libraries were prepared using Chromium Next GEM Single Cell 3ʹ GEM (10X Genomics), Library & Gel Bead Kit v3.1 (10X Genomics), Chromium Next GEM Chip G Single Cell Kit (10X Genomics), and Single Index Kit T Set A (10X Genomics) following the manufacturer’s recommended protocol. Sample cDNA libraries were then sequenced using the Illumina NovaSeq 6000 (Novogene, USA) with an average depth of 100,000 reads per cell.

Paired-end sequencing data were processed with Cell Ranger v7.0 (10X Genomics) for sequence alignment to mm10 mouse genome assembly. Cell Ranger output files were imported into the R program version 3.4 using Seurat v4(*115*) for downstream analysis. Cells with unique feature counts <200 or >4000 were removed for further analysis, along with cells containing mitochondrial gene counts greater than 10%. Data were normalized using the default settings and scaled using ScaleData function regressing out mitochondrial gene expression. RunPCA function was used to run principal component (PC) analysis, and the number of PCs were determined using elbow plot and jack straw analyses. UMAP, a nonlinear dimensional reduction technique was chosen to visualize the cell population. Data were then subjected to clustering using FindNeighbors and FindClusters functions (resolution = 0.3). Follicular regulatory T cells were extracted from the normalized data using the Subset function and a criterion of Foxp3 > 0. Differentially expressed genes (DEGs) in distinct cell clusters from either Tfh or Tfr compartments were identified using the FindMarkers function. Gene Ontology (GO) term enrichment analysis of filtered DEGs was performed using EnrichR(*116*).

### Expression and purification of Neuraminidase A and Neuraminidase S

NeuA/S plasmid were transformed into electrocompetent *E. coli* BL21 (DE3) cells individually. The cells were then cultured overnight into 100 mL of liquid culture (lysogeny broth plus 100 μg/mL ampicillin). In the following day, the culture was added to 1 L of liquid culture (lysogeny broth plus 100 μg/mL ampicillin) and incubated at 37°C, 220 rpm until OD 0.6. Once OD∼0.6, the NeuA expression was induced by a final concentration of 1mM IPTG and incubated at 16 °C, overnight. In the following day, cells were centrifuged at 3400×*g* for 35 minutes, 4°C, and the pellets were resuspended in binding buffer (50 mM Tris, 500 mM NaCl, 5 mM imidazole pH 8) with 300ug/mL lysozyme and incubated for 1 hour on ice. Cells were then sonicated using constant pulse for 15 seconds on and 45 seconds off on ice for ten times. Cells were centrifuged for 35 minutes at 18000 g and 4°C. The supernatant was then filtered and stored at 4°C for purification. A 5 mL Ni^2+^-Affinity chromatography column was equilibrated with the binding buffer for 15 column volumes. The supernatant was then loaded to the column. The column was then washed with 100 mL of wash buffer (50 mM Tris, 500 mM NaCl, 60 mM imidazole pH 8). The NeuA protein was then eluted from the column using an elution buffer (50 mM Tris, 500 mM NaCl, 200 mM imidazole pH 8) into 1 mL of fractions. The fractions containing proteins were then combined and the protein dialyzed against PBS pH 7 overnight at 4 °C. The protein was then concentrated using AMICON Ultra Centrifugal Filters (Thermo Fisher) and the size of the protein was validated by SDS-PAGE. The concentration of the protein was then validated using BCA assay.

### Protein A-APEX2 construct, expression, and purification

pET-11a plasmid was used as the vector backbone to construct the His6-proteinA-APEX2(C32S) expression plasmid. The His6-APEX2(C32S) construct in pET-11a was a gift from Dr. Yun (Kurt) Mou’s laboratory at the Institute of Biomedical Sciences, Academia Sinica. A truncated form of protein A was subsequently inserted into this plasmid to generate the His6-proteinA-APEX2(C32S) construct. The His6-proteinA-APEX2(C32S) plasmid was transformed into BL21 (DE3) pLysS *E. coli* and plated on a LB agar supplemented with 100 µg/mL ampicillin to grow overnight at 37°C. A single colony was inoculated in 100 mL LB (Lysogeny Broth, Miller) medium and grown overnight at 37°C in a shaking incubator at 225 rpm. 20 mL of the starter culture was used to inoculate a 3L culture in LB medium supplemented with 100 µg/mL ampicillin and was incubated at 37°C on a shaker at 250 rpm until O.D. ∼0.6. The culture was then chilled at 4°C for 30 minutes. Expression was induced by adding fresh IPTG (0.50 mM) and incubated at 18°C on a shaker at 200 rpm overnight. The culture was then centrifuged at 6,000 x g and 4°C for 30 minutes. The pellets were stored at -80°C until needed. For protein purification, the frozen pellets were resuspended on ice in 100 mL chilled lysis buffer (50 mM Tris at pH 8, 300 mM NaCl, and 5 mM Imidazole) with 1 mM phenylmethylsulfonyl fluoride (PMSF) and 0.25 mM EDTA. The lysate was emulsified while on ice. The cells were lysed through high-pressure homogenization (Avestin Emulsiflex C3), then centrifuged at 16,000 x g and 4°C for 30 minutes and the soluble fractions were moved to a fresh 200 mL flask. The soluble fraction was passed through a 5 mL HisTrap FF column (Nickel affinity column). The column was washed with 100 mL of wash buffer (50 mM Tris at pH 8, 300 mM NaCl, and 25 mM imidazole) at a rate of 5 mL/minute and protein A-APEX2 was eluted with 30 mL of elution buffer (50 mM Tris at pH 8, 300 mM NaCl, and 250 mM imidazole) with a 10% to 100% linear gradient at a rate of 2 mL/minute. 1.5 mL elution fractions were collected. The elution fractions were analyzed by SDS-PAGE. The pure samples were pooled and dialyzed overnight at 4°C in buffer (100 mM Tris at pH 7.5, 200 mM NaCl, 0.2 mM EDTA, and 2 mM DTT). Protein concentration was determined using a Bradford assay and aliquots of the protein were flash frozen in liquid nitrogen and stored at -80°C in a final glycerol concentration of 20%.

### CD4^+^ T cell surface biotinylation using pre-complexed Siglec-10-Fc and Protein A-APEX2

C57BL/6J mice were immunized with 200 µL of 1mM OVA-liposomes via tail vein injection. Mice were euthanized on day 4 post-immunization, and spleens were harvested and dissociated into single cell suspension. Follicular CD4^+^ T cells were enriched from pooled splenocytes following a custom magnetic bead-based negative selection protocol. Briefly, splenocytes were incubated with biotinylated antibodies at a concentration of 1 µg/mL per antibody that target mouse cell surface markers other than follicular CD4^+^ T cells (B220, CD19, IgD, IgM, CD11c, CD11b, CD162, Ly6C, Gr-1, Ter119, CD49b, TCRγ/δ, and CD8a). After 30 min of incubation, cells were washed and resuspended in 1X PBS containing 0.5% BSA and 2 mM EDTA, and anti-biotin microbeads (Miltenyi) were added to cell suspension at 20 µL per 10^7^ cells. After 20 min incubation on ice, cells were passed through the LS columns (Miltenyi) placed in magnetic separators. Flow-through containing follicular CD4^+^ T cells was collected in cold complete RPMI media. Enriched cells were washed twice with 1X PBS and distributed in a 96-well V-bottom plate, with approximately 1 million cells per well. Cells were incubated with 1X PBS containing 2 µg/mL Siglec-10-Fc (or R119A Siglec-10-Fc) pre-complexed with hemin-containing Protein A-APEX2 (1:2 molar ratio) for 30 min at RT. Following incubation, cells were washed with 1X PBS and incubated with 500 µM biotin-tyramide (MilliporeSigma) in 1X PBS for 30 min at RT. Cells were washed with 1X PBS and incubated with 500 µM H_2_O_2_ for 1 min at RT to initiate protein biotinylation. Reaction was quenched by washing cells 3 times with 1X PBS supplemented with 5 mM Trolox (MilliporeSigma) and 10 mM sodium ascorbate (MilliporeSigma). Finally, cells were then washed once with 1X PBS, pelleted, snapped frozen in liquid N_2_, and stored in -80°C freezer prior to MS proteomic analysis. Conventional and imaging flow cytometric analyses were performed to validate the level and localization of biotinylated proteins.

### Sample preparation for LC-MS/MS

Cell pellets were lysed in 500 µL RIPA lysis buffer (Pierce PI89900) supplemented with Halt Protease and Phosphatase Inhibitor Cocktail (Thermo Fisher 78442), and lysates were clarified by centrifugation at 12,000 x g for 10 min at 4°C. Soluble lysates proceeded to automated streptavidin enrichment using KingFisher Duo Prime and on-bead digestion as previously described(*117*) . In brief, 200 µL of streptavidin magnetic beads (Pierce PI88817) were pre-washed with lysis buffer, binding to lysates at 4°C for 18 h, washed 2X with lysis buffer and 1X with PBS. The beads with biotinylated proteins bound were then transferred for reduction with 300 µL of 10 mM dithiothreitol for 45 min, alkylated with 300 µL of 50 mM iodoacetamide for 45 min at RT, washed 1X with 100 mM ammonium bicarbonate, and proceeded to on-bead trypsin digestion with 600 ng of trypsin at 37°C for 5 h. Peptides were dried using GeneVac, desalted with C18 ziptips (Millipore ZTC18S960) and recovered in buffer A (0.1% formic acid; FA) prior to mass spectrometry analysis.

### Mass spectrometry

The samples were analyzed using Orbitrap Exploris 480 Mass Spectrometer (Thermo Fisher). Reverse phase separation of the peptides was performed with an EASY-Spray HPLC column (15 cm x 75 μm ID, Thermo Fisher). Peptides were eluted with a solvent B gradient (80% acetonitrile and 0.1% FA) for 90 min. The gradient was run at 400 nL/min with analytical column temperature set at 45°C. Data were analyzed using Proteome Discoverer (v3.1 CHIMERYS, prediction model inferys_3.0.0_fragmentation) against the concatenated database of the *mus musculus* proteome (UP000000589, v2024-03-25), with relaxed false discovery rate set at 5% and restricted at 1%. Search parameters included a maximum of two missed trypsin cleavages, a precursor mass tolerance of 20 ppm, a fragment mass tolerance of 0.01 Da, with the constant modification carbamidomethylation (C), and variable modification of oxidation (M) with maximum of three. Only tryptic peptides were searched, and MS1 intensities were normalized using the total peptide amount. Statistical analysis was performed using Proteome Discoverer (v3.1). Protein abundances were calculated using the sum of the peptide peak MS1 intensities. Hypothesis testing was performed using ANOVA (individual proteins) with Protein Abundance Based protein ratio calculation. Missing values were imputed with low abundance resampling using the bottom 5% of all detected values. Mass spectrometry spectrum files are available on MassIVE: MSV000100018.

### Quantification and statistical analysis

All data and statistical analyses were processed and visualized using GraphPad Prism software (version 10.1.2). In all graphs, data are presented as the mean ± standard error of the mean (SEM) unless otherwise indicated. For comparisons between two independent experimental groups, statistical significance was determined using the non-parametric Mann-Whitney *U* test. For experimental designs involving multiple groups, a one-way analysis of variance (ANOVA) was performed followed by Tukey’s post-hoc test for multiple comparisons. For datasets with two or more independent variables, a two-way ANOVA was utilized followed by Sidak’s post-hoc multiple comparisons test. For survival analyses, data were visualized using Kaplan-Meier survival curves, and statistical differences were determined using the Log-rank (Mantel-Cox) test. For proximity labeling and single-cell RNA sequencing datasets, specific statistical cutoffs (-log10(*p*-value) > 1.3 and log2 fold-change > 1.0) were applied as detailed in their respective methods sections.

## ACKNOWLEDGMENTS

We thank Dr. Mark Shlomchik for kindly providing the CD20^CreERT2^ mice, and Dr. Jamey Marth for generously providing the ST3Gal1^KO^ and ST3Gal4^KO^ mice. We are also grateful to Sarah Knight for her excellent technical assistance with mouse breeding and crosses. We thank James C. Paulson for their support in the early stages of this work. This study is funded by grants from NIAID (R21AI128598), CIHR (203698), and Canada Research Chair in Chemical Glycoimmunology to MSM. JRE is supported by fellowships from the Alberta Innovates Graduate Student Scholarship, Canadian Arthritis Society, and Izaak Walton Killam Memorial Scholarship. This work was supported in part by infrastructure CFI awards (JELF: 37833, 39051 and BRIF: 42736) and Striving for Pandemic Preparedness – the Alberta Research Consortium (SPP-ARC).

## REFERENCES

1. G. D. Victora, M. C. Nussenzweig, Germinal Centers. Annu Rev Immunol 40, 413–442 (2022).

2. D. Dominguez-Sola, G. D. Victora, C. Y. Ying, R. T. Phan, M. Saito, M. C. Nussenzweig, R. Dalla-Favera, The proto-oncogene MYC is required for selection in the germinal center and cyclic reentry. Nat Immunol 13, 1083–1091 (2012).

3. A. L. Raybuck, S. H. Cho, J. Li, M. C. Rogers, K. Lee, C. L. Williams, M. Shlomchik, J. W. Thomas, J. Chen, J. V. Williams, M. R. Boothby, B Cell-Intrinsic mTORC1 Promotes Germinal Center-Defining Transcription Factor Gene Expression, Somatic Hypermutation, and Memory B Cell Generation in Humoral Immunity. Journal of immunology (Baltimore, Md. : 1950) 200, 2627–2639 (2018).

4. J. Ersching, A. Efeyan, L. Mesin, J. T. Jacobsen, G. Pasqual, B. C. Grabiner, D. Dominguez-Sola, D. M. Sabatini, G. D. Victora, Germinal Center Selection and Affinity Maturation Require Dynamic Regulation of mTORC1 Kinase. Immunity 46, 1045–1058.e1046 (2017).

5. P. T. Sage, A. H. Sharpe, The multifaceted functions of follicular regulatory T cells. Current opinion in immunology 67, 68–74 (2020).

6. J.-M. Lee, P. L. Raeder, R. B. Rashid, H. Zhang, C. S. Nelson, S. G. Richardson, K. P. Burke, P. Chandrakar, M. A. Podestà, M. G. Gempler, C. Batty, D. Kong, S. J. Gavzy, M. W. Shirkey, C. Yang, J. S. Bromberg, M. C. Haigis, W. A. Marasco, A. H. Sharpe, P. T. Sage, Stability and progressive differentiation of TFR cells are intrinsically and extrinsically controlled by TFH programs. Nature Immunology 27, 336–347 (2026).

7. W. Luo, F. Weisel, M. J. Shlomchik, B Cell Receptor and CD40 Signaling Are Rewired for Synergistic Induction of the c-Myc Transcription Factor in Germinal Center B Cells. Immunity 48, 313–326.e315 (2018).

8. J. R. Enterina, J. Jung, M. S. Macauley, Coordinated roles for glycans in regulating the inhibitory function of CD22 on B cells. Biomed J 42, 218–232 (2019).

9. Y. Wen, Y. Jing, L. Yang, D. Kang, P. Jiang, N. Li, J. Cheng, J. Li, X. Li, Z. Peng, X. Sun, H. Miller, Z. Sui, Q. Gong, B. Ren, W. Yin, C. Liu, The regulators of BCR signaling during B cell activation. Blood Sci 1, 119–129 (2019).

10. J. R. Enterina, S. Sarkar, L. Streith, J. Jung, B. M. Arlian, S. J. Meyer, H. Takematsu, C. Xiao, T. A. Baldwin, L. Nitschke, M. J. Shlomchick, J. C. Paulson, M. S. Macauley, Coordinated changes in glycosylation regulate the germinal center through CD22. Cell reports 38, 110512 (2022).

11. Y. Naito, H. Takematsu, S. Koyama, S. Miyake, H. Yamamoto, R. Fujinawa, M. Sugai, Y. Okuno, G. Tsujimoto, T. Yamaji, Y. Hashimoto, S. Itohara, T. Kawasaki, A. Suzuki, Y. Kozutsumi, Germinal center marker GL7 probes activation-dependent repression of N-glycolylneuraminic acid, a sialic acid species involved in the negative modulation of B-cell activation. Mol Cell Biol 27, 3008–3022 (2007).

12. M. S. Macauley, N. Kawasaki, W. Peng, S.-H. Wang, Y. He, B. M. Arlian, R. McBride, R. Kannagi, K.-H. Khoo, J. C. Paulson, Unmasking of CD22 Co-receptor on Germinal Center B-cells Occurs by Alternative Mechanisms in Mouse and Man. Journal of Biological Chemistry 290, 30066–30077 (2015).

13. S. J. Meyer, M. Steffensen, A. Acs, T. Weisenburger, C. Wadewitz, T. H. Winkler, L. Nitschke, CD22 Controls Germinal Center B Cell Receptor Signaling, Which Influences Plasma Cell and Memory B Cell Output. Journal of immunology (Baltimore, Md. : 1950) 207, 1018–1032 (2021).

14. A. Hoffmann, S. Kerr, J. Jellusova, J. Zhang, F. Weisel, U. Wellmann, T. H. Winkler, B. Kneitz, P. R. Crocker, L. Nitschke, Siglec-G is a B1 cell-inhibitory receptor that controls expansion and calcium signaling of the B1 cell population. Nat Immunol 8, 695–704 (2007).

15. G. Whitney, S. Wang, H. Chang, K. Y. Cheng, P. Lu, X. D. Zhou, W. P. Yang, M. McKinnon, M. Longphre, A new siglec family member, siglec-10, is expressed in cells of the immune system and has signaling properties similar to CD33. European journal of biochemistry / FEBS 268, 6083–6096 (2001).

16. A. Hoffmann, S. Kerr, J. Jellusova, J. Zhang, F. Weisel, U. Wellmann, T. H. Winkler, B. Kneitz, P. R. Crocker, L. Nitschke, Siglec-G is a B1 cell–inhibitory receptor that controls expansion and calcium signaling of the B1 cell population. Nature Immunology 8, 695–704 (2007).

17. J. Müller, B. Lunz, I. Schwab, A. Acs, F. Nimmerjahn, C. Daniel, L. Nitschke, Siglec-G Deficiency Leads to Autoimmunity in Aging C57BL/6 Mice. Journal of immunology (Baltimore, Md. : 1950) 195, 51–60 (2015).

18. R. E. Forgione, C. Di Carluccio, J. Guzmán-Caldentey, R. Gaglione, F. Battista, F. Chiodo, Y. Manabe, A. Arciello, P. Del Vecchio, K. Fukase, A. Molinaro, S. Martín-Santamaría, P. R. Crocker, R. Marchetti, A. Silipo, Unveiling Molecular Recognition of Sialoglycans by Human Siglec-10. iScience 23, 101231 (2020).

19. E. N. Schmidt, D. Lamprinaki, K. A. McCord, M. Joe, M. Sojitra, A. Waldow, J. Nguyen, J. Monyror, E. N. Kitova, F. Mozaneh, X. Y. Guo, J. Jung, J. R. Enterina, G. C. Daskhan, L. Han, A. R. Krysler, C. R. Cromwell, B. P. Hubbard, L. J. West, M. Kulka, S. Sipione, J. S. Klassen, R. Derda, T. L. Lowary, L. K. Mahal, M. R. Riddell, M. S. Macauley, Siglec-6 mediates the uptake of extracellular vesicles through a noncanonical glycolipid binding pocket. Nature communications 14, 2327 (2023).

20. F. Pfrengle, M. S. Macauley, N. Kawasaki, J. C. Paulson, Copresentation of antigen and ligands of Siglec-G induces B cell tolerance independent of CD22. Journal of immunology (Baltimore, Md. : 1950) 191, 1724–1731 (2013).

21. T. Angata, Possible Influences of Endogenous and Exogenous Ligands on the Evolution of Human Siglecs. Front Immunol 9, 2885 (2018).

22. J. Choi, H. Diao, C. E. Faliti, J. Truong, M. Rossi, S. Bélanger, B. Yu, A. W. Goldrath, M. E. Pipkin, S. Crotty, Bcl-6 is the nexus transcription factor of T follicular helper cells via repressor-of-repressor circuits. Nat Immunol 21, 777–789 (2020).

23. G. Simonetti, M. T. Bertilaccio, T. V. Rodriguez, B. Apollonio, A. Dagklis, M. Rocchi, A. Innocenzi, S. Casola, T. H. Winkler, L. Nitschke, M. Ponzoni, F. Caligaris-Cappio, P. Ghia, SIGLEC-G deficiency increases susceptibility to develop B-cell lymphoproliferative disorders. Haematologica 99, 1356–1364 (2014).

24. A. M. Bock, N. Epperla, Therapeutic landscape of primary refractory and relapsed diffuse large B-cell lymphoma: Recent advances and emerging therapies. J Hematol Oncol 18, 68 (2025).

25. M. Pfreundschuh, L. Trümper, A. Osterborg, R. Pettengell, M. Trneny, K. Imrie, D. Ma, D. Gill, J. Walewski, P. L. Zinzani, R. Stahel, S. Kvaloy, O. Shpilberg, U. Jaeger, M. Hansen, T. Lehtinen, A. López-Guillermo, C. Corrado, A. Scheliga, N. Milpied, M. Mendila, M. Rashford, E. Kuhnt, M. Loeffler, CHOP-like chemotherapy plus rituximab versus CHOP-like chemotherapy alone in young patients with good-prognosis diffuse large-B-cell lymphoma: a randomised controlled trial by the MabThera International Trial (MInT) Group. Lancet Oncol 7, 379–391 (2006).

26. K. Dybkær, M. Bøgsted, S. Falgreen, J. S. Bødker, M. K. Kjeldsen, A. Schmitz, A. E. Bilgrau, Z. Y. Xu-Monette, L. Li, K. S. Bergkvist, M. B. Laursen, M. Rodrigo-Domingo, S. C. Marques, S. B. Rasmussen, M. Nyegaard, M. Gaihede, M. B. Møller, R. J. Samworth, R. D. Shah, P. Johansen, T. C. El-Galaly, K. H. Young, H. E. Johnsen, Diffuse large B-cell lymphoma classification system that associates normal B-cell subset phenotypes with prognosis. J Clin Oncol 33, 1379–1388 (2015).

27. E. Frei, C. Visco, Z. Y. Xu-Monette, S. Dirnhofer, K. Dybkær, A. Orazi, G. Bhagat, E. D. Hsi, J. H. van Krieken, M. Ponzoni, R. S. Go, M. A. Piris, M. B. Møller, K. H. Young, A. Tzankov, Addition of rituximab to chemotherapy overcomes the negative prognostic impact of cyclin E expression in diffuse large B-cell lymphoma. J Clin Pathol 66, 956–961 (2013).

28. A. Reddy, J. Zhang, N. S. Davis, A. B. Moffitt, C. L. Love, A. Waldrop, S. Leppa, A. Pasanen, L. Meriranta, M. L. Karjalainen-Lindsberg, P. Nørgaard, M. Pedersen, A. O. Gang, E. Høgdall, T. B. Heavican, W. Lone, J. Iqbal, Q. Qin, G. Li, S. Y. Kim, J. Healy, K. L. Richards, Y. Fedoriw, L. Bernal-Mizrachi, J. L. Koff, A. D. Staton, C. R. Flowers, O. Paltiel, N. Goldschmidt, M. Calaminici, A. Clear, J. Gribben, E. Nguyen, M. B. Czader, S. L. Ondrejka, A. Collie, E. D. Hsi, E. Tse, R. K. H. Au-Yeung, Y. L. Kwong, G. Srivastava, W. W. L. Choi, A. M. Evens, M. Pilichowska, M. Sengar, N. Reddy, S. Li, A. Chadburn, L. I. Gordon, E. S. Jaffe, S. Levy, R. Rempel, T. Tzeng, L. E. Happ, T. Dave, D. Rajagopalan, J. Datta, D. B. Dunson, S. S. Dave, Genetic and Functional Drivers of Diffuse Large B Cell Lymphoma. Cell 171, 481–494.e415 (2017).

29. C. Ding, Y. Liu, Y. Wang, B. K. Park, C. Y. Wang, P. Zheng, Y. Liu, Siglecg limits the size of B1a B cell lineage by down-regulating NFkappaB activation. PloS one 2, e997 (2007).

30. G. V. Zuccarino-Catania, S. Sadanand, F. J. Weisel, M. M. Tomayko, H. Meng, S. H. Kleinstein, K. L. Good-Jacobson, M. J. Shlomchik, CD80 and PD-L2 define functionally distinct memory B cell subsets that are independent of antibody isotype. Nat Immunol 15, 631–637 (2014).

31. A. D. Gitlin, Z. Shulman, M. C. Nussenzweig, Clonal selection in the germinal centre by regulated proliferation and hypermutation. Nature 509, 637–640 (2014).

32. W. Ise, K. Fujii, K. Shiroguchi, A. Ito, K. Kometani, K. Takeda, E. Kawakami, K. Yamashita, K. Suzuki, T. Okada, T. Kurosaki, T Follicular Helper Cell-Germinal Center B Cell Interaction Strength Regulates Entry into Plasma Cell or Recycling Germinal Center Cell Fate. Immunity 48, 702–715.e704 (2018).

33. C. R. Nowosad, K. M. Spillane, P. Tolar, Germinal center B cells recognize antigen through a specialized immune synapse architecture. Nat Immunol 17, 870–877 (2016).

34. G. D. Victora, T. A. Schwickert, D. R. Fooksman, A. O. Kamphorst, M. Meyer-Hermann, M. L. Dustin, M. C. Nussenzweig, Germinal center dynamics revealed by multiphoton microscopy with a photoactivatable fluorescent reporter. Cell 143, 592–605 (2010).

35. C. G. Vinuesa, M. A. Linterman, D. Yu, I. C. MacLennan, Follicular Helper T Cells. Annu Rev Immunol 34, 335–368 (2016).

36. S. Crotty, T Follicular Helper Cell Biology: A Decade of Discovery and Diseases. Immunity 50, 1132–1148 (2019).

37. M. A. Linterman, W. Pierson, S. K. Lee, A. Kallies, S. Kawamoto, T. F. Rayner, M. Srivastava, D. P. Divekar, L. Beaton, J. J. Hogan, S. Fagarasan, A. Liston, K. G. C. Smith, C. G. Vinuesa, Foxp3+ follicular regulatory T cells control the germinal center response. Nature medicine 17, 975–982 (2011).

38. D. Liu, J. Yan, J. Sun, B. Liu, W. Ma, Y. Li, X. Shao, H. Qi, BCL6 controls contact-dependent help delivery during follicular T-B cell interactions. Immunity 54, 2245–2255.e2244 (2021).

39. C. H. Yeh, J. Finney, T. Okada, T. Kurosaki, G. Kelsoe, Primary germinal center-resident T follicular helper cells are a physiologically distinct subset of CXCR5(hi)PD-1(hi) T follicular helper cells. Immunity 55, 272–289.e277 (2022).

40. H. Xu, X. Li, D. Liu, J. Li, X. Zhang, X. Chen, S. Hou, L. Peng, C. Xu, W. Liu, L. Zhang, H. Qi, Follicular T-helper cell recruitment governed by bystander B cells and ICOS-driven motility. Nature 496, 523–527 (2013).

41. J. Munday, S. Kerr, J. Ni, A. L. Cornish, J. Q. Zhang, G. Nicoll, H. Floyd, M. G. Mattei, P. Moore, D. Liu, P. R. Crocker, Identification, characterization and leucocyte expression of Siglec-10, a novel human sialic acid-binding receptor. The Biochemical journal 355, 489–497 (2001).

42. B. H. Duong, H. Tian, T. Ota, G. Completo, S. Han, J. L. Vela, M. Ota, M. Kubitz, N. Bovin, J. C. Paulson, D. Nemazee, Decoration of T-independent antigen with ligands for CD22 and Siglec-G can suppress immunity and induce B cell tolerance in vivo. J Exp Med 207, 173–187 (2010).

43. E. Rodrigues, J. Jung, H. Park, C. Loo, S. Soukhtehzari, E. N. Kitova, F. Mozaneh, G. Daskhan, E. N. Schmidt, V. Aghanya, S. Sarkar, L. Streith, C. D. St Laurent, L. Nguyen, J. P. Julien, L. J. West, K. C. Williams, J. S. Klassen, M. S. Macauley, A versatile soluble siglec scaffold for sensitive and quantitative detection of glycan ligands. Nature communications 11, 5091 (2020).

44. K. Sobczak, A. Antoñana-Vildosola, P. Valverde, M. A. Travecedo, Z. Jame-Chernaboo, E. N. Schmidt, S. D’Andrea, E. Valdaliso-Díez, I. Oyenarte, M. E. Laugieri, M. Joe, F. Mozaneh, S.-Y. Lin, A. Bosch, M. J. Moure, A. Franconetti, S. Y. Lee, J. E.-D. de Durana, L. Pérez-Gutierrez, A. Palazón, F. Marcelo, E. Fadda, F. Corzana, A. Gimeno, M. S. Macauley, J. Jiménez-Barbero, J. Ereño-Orbea, The unique molecular recognition features of Siglec-10: structural insights into sialoglycan and antibody interactions. bioRxiv, 2025.2006.2010.658867 (2025).

45. T. Hennet, D. Chui, J. C. Paulson, J. D. Marth, Immune regulation by the ST6Gal sialyltransferase. Proc Natl Acad Sci U S A 95, 4504–4509 (1998).

46. L. G. Ellies, D. Ditto, G. G. Levy, M. Wahrenbrock, D. Ginsburg, A. Varki, D. T. Le, J. D. Marth, Sialyltransferase ST3Gal-IV operates as a dominant modifier of hemostasis by concealing asialoglycoprotein receptor ligands. Proc Natl Acad Sci U S A 99, 10042–10047 (2002).

47. J. J. Priatel, D. Chui, N. Hiraoka, C. J. Simmons, K. B. Richardson, D. M. Page, M. Fukuda, N. M. Varki, J. D. Marth, The ST3Gal-I sialyltransferase controls CD8+ T lymphocyte homeostasis by modulating O-glycan biosynthesis. Immunity 12, 273–283 (2000).

48. J. Jung, E. N. Schmidt, H. C. Chang, Z. Jame-Chenarboo, J. R. Enterina, K. A. McCord, T. E. Gray, L. Kageler, C. D. St Laurent, C. Wang, R. A. Flynn, P. Wu, K. H. Khoo, M. S. Macauley, Understanding the Glycosylation Pathways Involved in the Biosynthesis of the Sulfated Glycan Ligands for Siglecs. ACS Chem Biol 20, 386–400 (2025).

49. X. Li, J. Zhou, W. Zhao, Q. Wen, W. Wang, H. Peng, Y. Gao, K. J. Bouchonville, S. M. Offer, K. Chan, Z. Wang, N. Li, H. Gan, Defining Proximity Proteome of Histone Modifications by Antibody-mediated Protein A-APEX2 Labeling. Genomics Proteomics Bioinformatics 20, 87–100 (2022).

50. J. Zhou, X. Li, Q. Wen, H. Gan, Antibody-Mediated Protein A-APEX2 Labeling (AMAPEX) for Proximity Proteome Exploration. Methods in molecular biology (Clifton, N.J.) 2953, 295–310 (2025).

51. H. W. Rhee, P. Zou, N. D. Udeshi, J. D. Martell, V. K. Mootha, S. A. Carr, A. Y. Ting, Proteomic mapping of mitochondria in living cells via spatially restricted enzymatic tagging. Science 339, 1328–1331 (2013).

52. P. Saini, G. Mirji, S. M. S. Islam, L. M. Simons, S. A. Bhat, A. P. Bonfanti, K. Muthumani, P. Agrawal, J. Cassel, H. Y. Tang, H. Tateno, R. Zhang, J. F. Hultquist, R. S. Shinde, M. Abdel-Mohsen, Targeting Interactions Between Siglec-10 and α3β1 Integrin Enhances Macrophage-Mediated Phagocytosis of Pancreatic Cancer. Cancer Res 86, 99–115 (2026).

53. J. Daly, L. Piatnitca, M. Al-Seragi, V. Krishnamoorthy, S. Wisnovsky, CRISPR activation screens map the genomic landscape of cancer glycome remodeling. Cell Genom, 101139 (2026).

54. R. Nakagawa, D. P. Calado, Positive Selection in the Light Zone of Germinal Centers. Frontiers in Immunology 12, (2021).

55. J. Pae, J. Ersching, T. B. R. Castro, M. Schips, L. Mesin, S. J. Allon, J. Ordovas-Montanes, C. Mlynarczyk, A. Melnick, A. Efeyan, A. K. Shalek, M. Meyer-Hermann, G. D. Victora, Cyclin D3 drives inertial cell cycling in dark zone germinal center B cells. Journal of Experimental Medicine 218, e20201699 (2020).

56. G. D. Victora, M. C. Nussenzweig, Germinal Centers. Annual Review of Immunology 40, 413–442 (2022).

57. A. J. Fike, S. B. Chodisetti, N. E. Wright, K. N. Bricker, P. P. Domeier, M. Maienschein-Cline, A. M. Rosenfeld, S. A. Luckenbill, J. L. Weber, N. M. Choi, E. T. Luning Prak, M. Mandal, M. R. Clark, Z. S. M. Rahman, STAT3 signaling in B cells controls germinal center zone organization and recycling. Cell reports 42, 112512 (2023).

58. W. Luo, L. Conter, R. A. Elsner, S. Smita, F. Weisel, D. Callahan, S. Wu, M. Chikina, M. Shlomchik, IL-21R signal reprogramming cooperates with CD40 and BCR signals to select and differentiate germinal center B cells. Sci Immunol 8, eadd1823 (2023).

59. S. Finkin, H. Hartweger, T. Y. Oliveira, E. E. Kara, M. C. Nussenzweig, Protein Amounts of the MYC Transcription Factor Determine Germinal Center B Cell Division Capacity. Immunity 51, 324–336.e325 (2019).

60. K. Hlavac, P. Pavelkova, L. Ondrisova, M. Mraz, FoxO1 signaling in B cell malignancies and its therapeutic targeting. FEBS letters 599, 2911–2931 (2025).

61. J. G. Cyster, C. D. C. Allen, B Cell Responses: Cell Interaction Dynamics and Decisions. Cell 177, 524–540 (2019).

62. B. Röder, L. Nitschke, The role of Siglec-G on B cells in autoimmune disease and leukemia. Seminars in arthritis and rheumatism 64, 152328 (2024).

63. K. A. Brzezicka, J. C. Paulson, Impact of Siglecs on autoimmune diseases. Molecular Aspects of Medicine 90, 101140 (2023).

64. M. Garcia-Lacarte, S. C. Grijalba, J. Melchor, A. Arnaiz-Leché, S. Roa, The PD-1/PD-L1 Checkpoint in Normal Germinal Centers and Diffuse Large B-Cell Lymphomas. Cancers 13, 4683 (2021).

65. M. Stebegg, S. D. Kumar, A. Silva-Cayetano, V. R. Fonseca, M. A. Linterman, L. Graca, Regulation of the Germinal Center Response. Frontiers in Immunology 9, (2018).

66. L. Nitschke, Siglec-G is a B-1 cell inhibitory receptor and also controls B cell tolerance. Annals of the New York Academy of Sciences 1362, 117–121 (2015).

67. S. Hutzler, L. Özgör, Y. Naito-Matsui, K. Kläsener, T. H. Winkler, M. Reth, L. Nitschke, The ligand-binding domain of Siglec-G is crucial for its selective inhibitory function on B1 cells. Journal of immunology (Baltimore, Md. : 1950) 192, 5406–5414 (2014).

68. L. Nitschke, CD22 and Siglec-G regulate inhibition of B-cell signaling by sialic acid ligand binding and control B-cell tolerance. Glycobiology 24, 807–817 (2014).

69. A. M. Khalil, J. C. Cambier, M. J. Shlomchik, B cell receptor signal transduction in the GC is short-circuited by high phosphatase activity. Science 336, 1178–1181 (2012).

70. D. T. Avery, E. K. Deenick, C. S. Ma, S. Suryani, N. Simpson, G. Y. Chew, T. D. Chan, U. Palendira, J. Bustamante, S. Boisson-Dupuis, S. Choo, K. E. Bleasel, J. Peake, C. King, M. A. French, D. Engelhard, S. Al-Hajjar, S. Al-Muhsen, K. Magdorf, J. Roesler, P. D. Arkwright, P. Hissaria, D. S. Riminton, M. Wong, R. Brink, D. A. Fulcher, J. L. Casanova, M. C. Cook, S. G. Tangye, B cell-intrinsic signaling through IL-21 receptor and STAT3 is required for establishing long-lived antibody responses in humans. J Exp Med 207, 155–171 (2010).

71. L. Mesin, J. Ersching, G. D. Victora, Germinal Center B Cell Dynamics. Immunity 45, 471–482 (2016).

72. A. R. Dvorscek, C. I. McKenzie, M. J. Robinson, Z. Ding, C. Pitt, K. O’Donnell, D. Zotos, R. Brink, D. M. Tarlinton, I. Quast, IL-21 has a critical role in establishing germinal centers by amplifying early B cell proliferation. EMBO Rep 23, e54677 (2022).

73. D. Suan, N. J. Kräutler, J. L. V. Maag, D. Butt, K. Bourne, J. R. Hermes, D. T. Avery, C. Young, A. Statham, M. Elliott, M. E. Dinger, A. Basten, S. G. Tangye, R. Brink, CCR6 Defines Memory B Cell Precursors in Mouse and Human Germinal Centers, Revealing Light-Zone Location and Predominant Low Antigen Affinity. Immunity 47, 1142–1153.e1144 (2017).

74. M. A. ElTanbouly, V. Ramos, A. J. MacLean, S. T. Chen, M. Loewe, S. Steinbach, T. Ben Tanfous, B. Johnson, M. Cipolla, A. Gazumyan, T. Y. Oliveira, M. C. Nussenzweig, Role of affinity in plasma cell development in the germinal center light zone. J Exp Med 221, (2024).

75. N. J. Kräutler, D. Suan, D. Butt, K. Bourne, J. R. Hermes, T. D. Chan, C. Sundling, W. Kaplan, P. Schofield, J. Jackson, A. Basten, D. Christ, R. Brink, Differentiation of germinal center B cells into plasma cells is initiated by high-affinity antigen and completed by Tfh cells. J Exp Med 214, 1259–1267 (2017).

76. J. Jellusova, U. Wellmann, K. Amann, T. H. Winkler, L. Nitschke, CD22 × Siglec-G Double-Deficient Mice Have Massively Increased B1 Cell Numbers and Develop Systemic Autoimmunity. The Journal of Immunology 184, 3618–3627 (2010).

77. P. R. Crocker, J. C. Paulson, A. Varki, Siglecs and their roles in the immune system. Nature Reviews Immunology 7, 255–266 (2007).

78. S. Y. Lin, E. N. Schmidt, K. Takahashi-Yamashiro, M. S. Macauley, Roles for Siglec-glycan interactions in regulating immune cells. Seminars in immunology 77, 101925 (2024).

79. F. Gasparrini, C. Feest, A. Bruckbauer, P. K. Mattila, J. Müller, L. Nitschke, D. Bray, F. D. Batista, Nanoscale organization and dynamics of the siglec CD22 cooperate with the cytoskeleton in restraining BCR signalling. The EMBO journal 35, 258–280 (2016).

80. L. Wasim, F. H. M. Buhari, M. Yoganathan, T. Sicard, J. Ereno-Orbea, J. P. Julien, B. Treanor, N-Linked Glycosylation Regulates CD22 Organization and Function. Front Immunol 10, 699 (2019).

81. J. Wang, J. Sun, L. N. Liu, D. B. Flies, X. Nie, M. Toki, J. Zhang, C. Song, M. Zarr, X. Zhou, X. Han, K. A. Archer, T. O’Neill, R. S. Herbst, A. N. Boto, M. F. Sanmamed, S. Langermann, D. L. Rimm, L. Chen, Siglec-15 as an immune suppressor and potential target for normalization cancer immunotherapy. Nature Medicine 25, 656–666 (2019).

82. G. Murugesan, V. G. Correia, A. S. Palma, W. Chai, C. Li, T. Feizi, E. Martin, B. Laux, A. Franz, K. Fuchs, B. Weigle, P. R. Crocker, Siglec-15 recognition of sialoglycans on tumor cell lines can occur independently of sialyl Tn antigen expression. Glycobiology 31, 44–54 (2021).

83. K. Boelaars, Y. van Kooyk, Targeting myeloid cells for cancer immunotherapy: Siglec-7/9/10/15 and their ligands. Trends in Cancer 10, 230–241 (2024).

84. A. Alborzian Deh Sheikh, S. Gomaa, X. Li, M. Routledge, K. Saigoh, N. Numoto, T. Angata, Y. Hitomi, H. Takematsu, M. Tsuiji, N. Ito, S. Kusunoki, T. Tsubata, A Guillain-Barré syndrome-associated SIGLEC10 rare variant impairs its recognition of gangliosides. Journal of Autoimmunity 116, 102571 (2021).

85. T. Tsubata, The ligand interactions of B cell Siglecs are involved in the prevention of autoimmunity to sialylated self-antigens and in the quality control of signaling-competent B cells. International immunology 35, 461–473 (2023).

86. D. Raïch-Regué, P. Resa-Infante, M. Gallemí, F. Laguia, X. Muñiz-Trabudua, J. Muñoz-Basagoiti, D. Perez-Zsolt, J. Chojnacki, S. Benet, B. Clotet, J. Martinez-Picado, N. Izquierdo-Useros, Role of Siglecs in viral infections: A double-edged sword interaction. Mol Aspects Med 90, 101113 (2023).

87. M. S. Macauley, J. C. Paulson, Siglecs induce tolerance to cell surface antigens by BIM-dependent deletion of the antigen-reactive B cells. Journal of immunology (Baltimore, Md. : 1950) 193, 4312–4321 (2014).

88. N. Hashimoto, S. Ito, A. Harazono, A. Tsuchida, Y. Mouri, A. Yamamoto, T. Okajima, Y. Ohmi, K. Furukawa, Y. Kudo, N. Kawasaki, K. Furukawa, Bidirectional signals generated by Siglec-7 and its crucial ligand tri-sialylated T to escape of cancer cells from immune surveillance. iScience 27, 111139 (2024).

89. Y. Zheng, X. Ma, D. Su, Y. Zhang, L. Yu, F. Jiang, X. Zhou, Y. Feng, F. Ma, The Roles of Siglec7 and Siglec9 on Natural Killer Cells in Virus Infection and Tumour Progression. J Immunol Res 2020, 6243819 (2020).

90. D. Perez-Zsolt, J. Muñoz-Basagoiti, J. Rodon, M. Elosua-Bayes, D. Raïch-Regué, C. Risco, M. Sachse, M. Pino, S. Gumber, M. Paiardini, J. Chojnacki, I. Erkizia, X. Muñiz-Trabudua, E. Ballana, E. Riveira-Muñoz, M. Noguera-Julian, R. Paredes, B. Trinité, F. Tarrés-Freixas, I. Blanco, V. Guallar, J. Carrillo, J. Blanco, A. Telenti, H. Heyn, J. Segalés, B. Clotet, J. Martinez-Picado, J. Vergara-Alert, N. Izquierdo-Useros, SARS-CoV-2 interaction with Siglec-1 mediates trans-infection by dendritic cells. Cellular & Molecular Immunology 18, 2676–2678 (2021).

91. H. H. Chou, H. Takematsu, S. Diaz, J. Iber, E. Nickerson, K. L. Wright, E. A. Muchmore, D. L. Nelson, S. T. Warren, A. Varki, A mutation in human CMP-sialic acid hydroxylase occurred after the Homo-Pan divergence. Proc Natl Acad Sci U S A 95, 11751–11756 (1998).

92. N. Kimura, K. Ohmori, K. Miyazaki, M. Izawa, Y. Matsuzaki, Y. Yasuda, H. Takematsu, Y. Kozutsumi, A. Moriyama, R. Kannagi, Human B-lymphocytes express alpha2-6-sialylated 6-sulfo-N-acetyllactosamine serving as a preferred ligand for CD22/Siglec-2. The Journal of biological chemistry 282, 32200–32207 (2007).

93. T. Hayakawa, Y. Satta, P. Gagneux, A. Varki, N. Takahata, Alu-mediated inactivation of the human CMP- N-acetylneuraminic acid hydroxylase gene. Proc Natl Acad Sci U S A 98, 11399–11404 (2001).

94. S. N. Henriques, L. Oliveira, R. F. Santos, A. M. Carmo, CD6-mediated inhibition of T cell activation via modulation of Ras. Cell Commun Signal 20, 184 (2022).

95. Z. Ning, K. Liu, H. Xiong, Roles of BTLA in Immunity and Immune Disorders. Frontiers in Immunology Volume 12 - 2021, (2021).

96. X. Xu, B. Hou, A. Fulzele, T. Masubuchi, Y. Zhao, Z. Wu, Y. Hu, Y. Jiang, Y. Ma, H. Wang, E. J. Bennett, G. Fu, E. Hui, PD-1 and BTLA regulate T cell signaling differentially and only partially through SHP1 and SHP2. Journal of Cell Biology 219, e201905085 (2020).

97. B. Li, M.-C. Zhong, C. C. Galindo, J. Dou, J. Qian, Z. Tang, D. Davidson, A. Veillette, SLAMF6 as a drug-targetable suppressor of T cell immunity against cancer. Nature, (2026).

98. M. A. Mintz, J. H. Felce, M. Y. Chou, V. Mayya, Y. Xu, J. W. Shui, J. An, Z. Li, A. Marson, T. Okada, C. F. Ware, M. Kronenberg, M. L. Dustin, J. G. Cyster, The HVEM-BTLA Axis Restrains T Cell Help to Germinal Center B Cells and Functions as a Cell-Extrinsic Suppressor in Lymphomagenesis. Immunity 51, 310–323.e317 (2019).

99. A. Cardani-Boulton, F. Lin, C. C. Bergmann, CD6 regulates CD4 T follicular helper cell differentiation and humoral immunity during murine coronavirus infection. J Virol 99, e0186424 (2025).

100. D. Hao, Y. Wang, L. Li, G. Qian, J. Liu, M. Li, Y. Zhang, R. Zhou, D. Yan, SHP-1 suppresses the antiviral innate immune response by targeting TRAF3. FASEB journal : official publication of the Federation of American Societies for Experimental Biology 34, 12392–12405 (2020).

101. B. S. Hostager, G. A. Bishop, Cutting edge: contrasting roles of TNF receptor-associated factor 2 (TRAF2) and TRAF3 in CD40-activated B lymphocyte differentiation. Journal of immunology (Baltimore, Md. : 1950) 162, 6307–6311 (1999).

102. P. Xie, B. S. Hostager, G. A. Bishop, Requirement for TRAF3 in signaling by LMP1 but not CD40 in B lymphocytes. J Exp Med 199, 661–671 (2004).

103. H. Qi, J. L. Cannons, F. Klauschen, P. L. Schwartzberg, R. N. Germain, SAP-controlled T-B cell interactions underlie germinal centre formation. Nature 455, 764–769 (2008).

104. N. M. Haynes, C. D. Allen, R. Lesley, K. M. Ansel, N. Killeen, J. G. Cyster, Role of CXCR5 and CCR7 in follicular Th cell positioning and appearance of a programmed cell death gene-1high germinal center-associated subpopulation. Journal of immunology (Baltimore, Md. : 1950) 179, 5099–5108 (2007).

105. C. N. Arnold, D. J. Campbell, M. Lipp, E. C. Butcher, The germinal center response is impaired in the absence of T cell-expressed CXCR5. European journal of immunology 37, 100–109 (2007).

106. J. Shi, S. Hou, Q. Fang, X. Liu, X. Liu, H. Qi, PD-1 Controls Follicular T Helper Cell Positioning and Function. Immunity 49, 264–274.e264 (2018).

107. G. J. Martinez, J. K. Hu, R. M. Pereira, J. S. Crampton, S. Togher, N. Bild, S. Crotty, A. Rao, Cutting Edge: NFAT Transcription Factors Promote the Generation of Follicular Helper T Cells in Response to Acute Viral Infection. J Immunol 196, 2015–2019 (2016).

108. M. Vaeth, M. Eckstein, P. J. Shaw, L. Kozhaya, J. Yang, F. Berberich-Siebelt, R. Clancy, D. Unutmaz, S. Feske, Store-Operated Ca(2+) Entry in Follicular T Cells Controls Humoral Immune Responses and Autoimmunity. Immunity 44, 1350–1364 (2016).

109. J. T. Jacobsen, W. Hu, R. C. TB, S. Solem, A. Galante, Z. Lin, S. J. Allon, L. Mesin, A. M. Bilate, A. Schiepers, A. K. Shalek, A. Y. Rudensky, G. D. Victora, Expression of Foxp3 by T follicular helper cells in end-stage germinal centers. Science 373, (2021).

110. F. Ke, Z. L. Benet, M. P. Maz, J. Liu, A. L. Dent, J. M. Kahlenberg, I. L. Grigorova, Germinal center B cells that acquire nuclear proteins are specifically suppressed by follicular regulatory T cells. Elife 12, (2023).

111. B. J. Laidlaw, Y. Lu, R. A. Amezquita, J. S. Weinstein, J. A. Vander Heiden, N. T. Gupta, S. H. Kleinstein, S. M. Kaech, J. Craft, Interleukin-10 from CD4(+) follicular regulatory T cells promotes the germinal center response. Sci Immunol 2, (2017).

112. J. Jung, J. R. Enterina, D. T. Bui, F. Mozaneh, P.-H. Lin, Nitin, C.-W. Kuo, E. Rodrigues, A. Bhattacherjee, P. Raeisimakiani, G. C. Daskhan, C. D. St. Laurent, K.-H. Khoo, L. K. Mahal, W. F. Zandberg, X. Huang, J. S. Klassen, M. S. Macauley, Carbohydrate Sulfation As a Mechanism for Fine-Tuning Siglec Ligands. ACS Chemical Biology 16, 2673–2689 (2021).

113. E. N. Schmidt, J. Jung, M. S. Macauley, Flow Cytometry-Based Detection of Siglec Ligands. Methods Mol Biol 2657, 181–193 (2023).

114. C. Di Carluccio, L. Cerofolini, M. Moreira, F. Rosu, L. Padilla-Cortés, G. R. Gheorghita, Z. Xu, A. Santra, H. Yu, S. Yokoyama, T. E. Gray, C. D. St Laurent, Y. Manabe, X. Chen, K. Fukase, M. S. Macauley, A. Molinaro, T. Li, B. A. Bensing, R. Marchetti, V. Gabelica, M. Fragai, A. Silipo, Molecular Insights into O-Linked Sialoglycans Recognition by the Siglec-Like SLBR-N (SLBR(UB10712)) of Streptococcus gordonii. ACS central science 10, 447–459 (2024).

115. Y. Hao, S. Hao, E. Andersen-Nissen, W. M. Mauck, 3rd, S. Zheng, A. Butler, M. J. Lee, A. J. Wilk, C. Darby, M. Zager, P. Hoffman, M. Stoeckius, E. Papalexi, E. P. Mimitou, J. Jain, A. Srivastava, T. Stuart, L. M. Fleming, B. Yeung, A. J. Rogers, J. M. McElrath, C. A. Blish, R. Gottardo, P. Smibert, R. Satija, Integrated analysis of multimodal single-cell data. Cell 184, 3573–3587.e3529 (2021).

116. Z. Xie, A. Bailey, M. V. Kuleshov, D. J. B. Clarke, J. E. Evangelista, S. L. Jenkins, A. Lachmann, M. L. Wojciechowicz, E. Kropiwnicki, K. M. Jagodnik, M. Jeon, A. Ma’ayan, Gene Set Knowledge Discovery with Enrichr. Curr Protoc 1, e90 (2021).

117. L. Pirayeshfard, S. Luo, J. M. Githaka, A. Saini, N. Touret, I. S. Goping, O. Julien, Comparing the BAD Protein Interactomes in 2D and 3D Cell Culture Using Proximity Labeling. Journal of proteome research 23, 3433–3443 (2024).

118. T. S. P. Heng, M. W. Painter, K. Elpek, V. Lukacs-Kornek, N. Mauermann, S. J. Turley, D. Koller, F. S. Kim, A. J. Wagers, N. Asinovski, S. Davis, M. Fassett, M. Feuerer, D. H. D. Gray, S. Haxhinasto, J. A. Hill, G. Hyatt, C. Laplace, K. Leatherbee, D. Mathis, C. Benoist, R. Jianu, D. H. Laidlaw, J. A. Best, J. Knell, A. W. Goldrath, J. Jarjoura, J. C. Sun, Y. Zhu, L. L. Lanier, A. Ergun, Z. Li, J. J. Collins, S. A. Shinton, R. R. Hardy, R. Friedline, K. Sylvia, J. Kang, C. The Immunological Genome Project, The Immunological Genome Project: networks of gene expression in immune cells. Nature Immunology 9, 1091–1094 (2008).

