## Supplementary Figures for "Siglec-G on B cells restrains the germinal center response by controlling T cell help during positive selection"

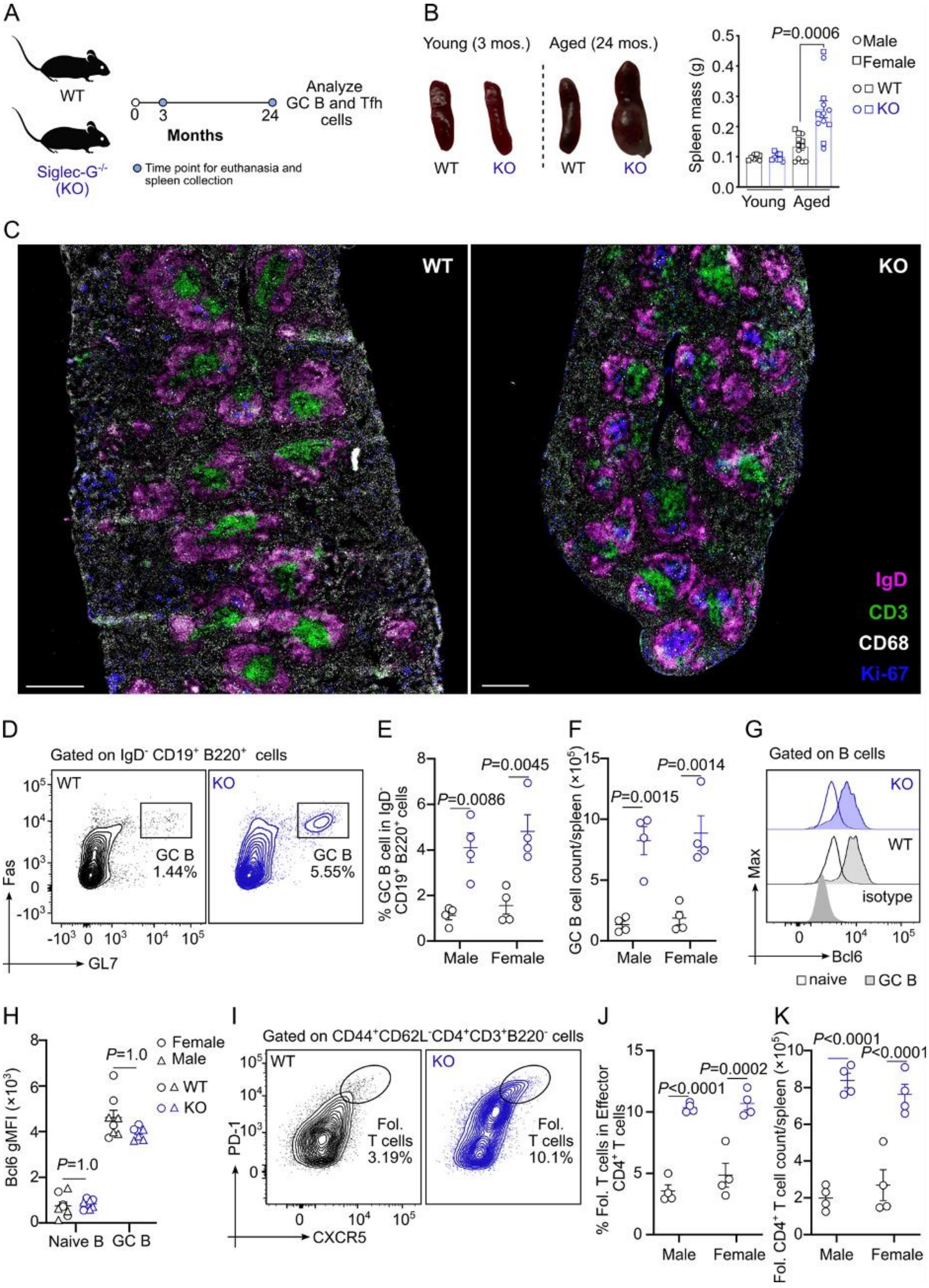

**Supplementary Figure 1. Systemic knockout of Siglec-G results in splenomegaly and spontaneous GCs in mice**. (**A**) Experimental timeline for monitoring and cellular analysis of spleens from WT and Siglec-$G^{KO}$ (KO) mice (**B**) Images and quantification of splenic mass from young and aged WT and KO mice. (**C**) Representative immunofluorescence images of spleens from 3 months old WT and KO mice. The scale bar represents 200 μM. (**D-F**) Flow cytometric plots (**D**) and quantifications of percent GC B cells (**E**) and absolute GC B cell numbers (**F**) from spleens of naive 3 months old WT and KO mice. (**G,H**) Flow cytometric plots (**G**) and quantification of transcription factor, Bcl6, expression levels (**H**) in GC B cells from spleens of naive 3 months old WT and KO mice. (**I-K**) Flow cytometric plots (**I**) and quantifications of percent follicular $CD4^+$ T cells (**E**) and absolute follicular $CD4^+$ T cell numbers (**F**) from spleens of naive 3 months old WT and KO mice. Data plots are presented as mean±SEM. Statistical analysis was performed using Kruskal-Wallis H test with post-hoc Dunn's multiple comparisons test for B, and two-way ANOVA with post-hoc Tukey's multiple comparisons test for E, F, H, J, and K.

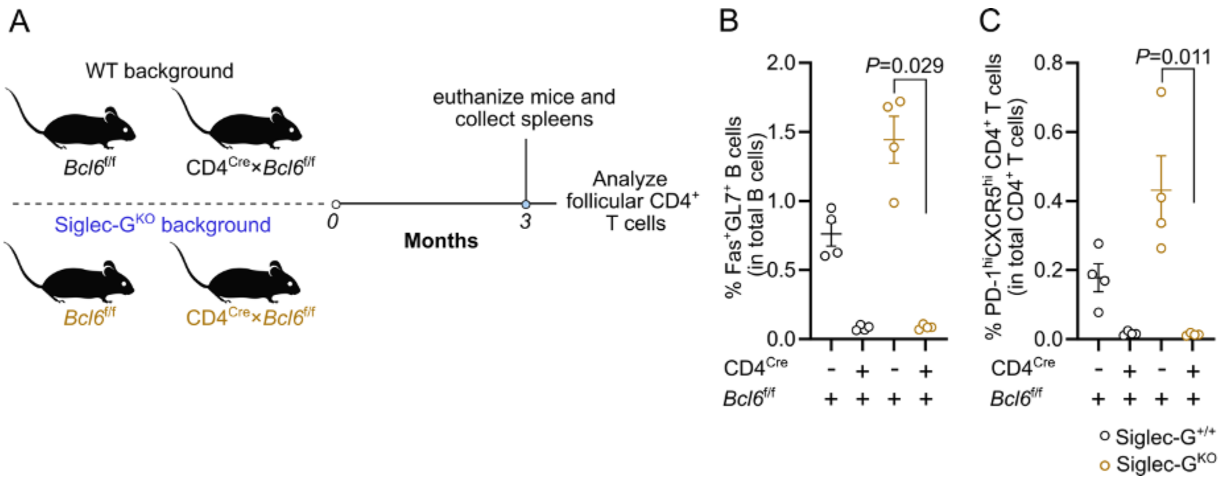


**Supplementary Figure 2. Spontaneously formed GC B cells in Siglec-G$^{KO}$ mice is Tfh cell-dependent (related to Figure 1)**. (**A**) Experimental scheme for tracking spontaneously formed GC B cells and follicular CD4$^+$ T cells from spleens of *Bcl6*$^{f/f}$ and CD4$^{Cre}$×*Bcl6*$^{f/f}$ Siglec-G$^{+/+}$ B6 mice, as well as spleens from *Bcl6*$^{f/f}$ and CD4$^{Cre}$×*Bcl6*$^{f/f}$ Siglec-G$^{KO}$ B6 mice. (**B,C**) Quantification of percent GC B cells (**B**) and percent follicular CD4$^+$ T cells (**C**) in total B cells and CD4$^+$ T cells, respectively. Data plots are presented as mean±SEM. Statistical analysis was performed using Kruskal-Wallis *H* test with post-hoc Dunn's multiple comparison test.

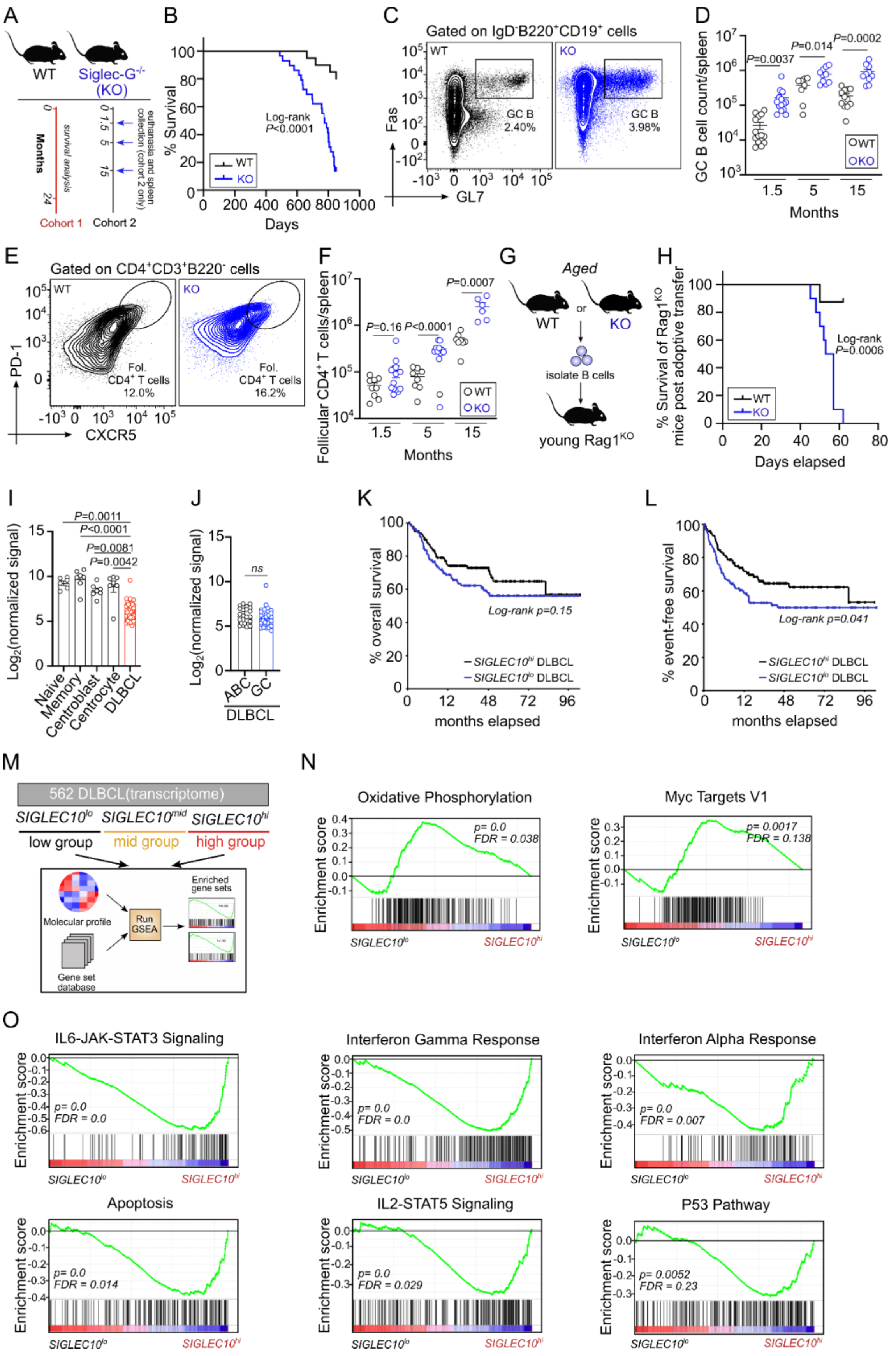
A
WT
Siglec-G-/- (KO)
Months
survival analysis
euthanasia and spleen collection (cohort 2 only)
Cohort 1
Cohort 2
B
% Survival
Days
Log-rank P<0.0001
WT
KO
C
Gated on IgD-B220+CD19+ cells
WT
KO
Fas
GL7
GC B 2.40%
GC B 3.98%
D
GC B cell count/spleen
P=0.0037
P=0.014
P=0.0002
Months
WT
KO
E
Gated on CD4+CD3+B220- cells
PD-1
CXCR5
Fol. CD4+ T cells 12.0%
Fol. CD4+ T cells 16.2%
F
Follicular CD4+ T cells/spleen
P=0.16
P<0.0001
P=0.0007
Months
G
Aged
WT
or
KO
isolate B cells
young Rag1KO
H
% Survival of Rag1KO mice post adoptive transfer
Log-rank P=0.0006
Days elapsed
I
Log2(normalized signal)
P=0.0011
P<0.0001
P=0.0081
P=0.0042
Naive
Memory
Centroblast
Centrocyte
DLBCL
J
Log2(normalized signal)
ns
ABC
GC
DLBCL
K
% overall survival
Log-rank p=0.15
SIGLEC10hi DLBCL
SIGLEC10lo DLBCL
months elapsed
L
% event-free survival
Log-rank p=0.041
SIGLEC10hi DLBCL
SIGLEC10lo DLBCL
months elapsed
M
562 DLBCL(transcriptome)
SIGLEC10lo
SIGLEC10mid
SIGLEC10hi
low group
mid group
high group
Molecular profile
Run GSEA
Enriched gene sets
Gene set database
N
Oxidative Phosphorylation
p= 0.0
FDR = 0.038
Myc Targets V1
p= 0.0017
FDR = 0.138
Enrichment score
SIGLEC10lo
SIGLEC10hi
O
IL6-JAK-STAT3 Signaling
p= 0.0
FDR = 0.0
Interferon Gamma Response
p= 0.0
FDR = 0.0
Interferon Alpha Response
p= 0.0
FDR = 0.007
Apoptosis
p= 0.0
FDR = 0.014
IL2-STAT5 Signaling
p= 0.0
FDR = 0.029
P53 Pathway
p= 0.0052
FDR = 0.23
Enrichment score
SIGLEC10lo
SIGLEC10hi

**Supplementary Figure 3. SIGLEC10 gene in human diffuse large B cell lymphoma (DLBCL) tumours.** (**A**) Experimental timeline for monitoring and cellular analysis of spleens from WT and Siglec-$G^{KO}$ mice. A group of WT and Siglec-$G^{KO}$ B6 mice were either aged for survival analysis (cohort 1) or sacrificed at defined time points for flow cytometric analysis of splenic B cell and T cell compartments (cohort 2). (**B**) Kaplan-Meier plot of % survival of aged WT and Siglec-$G^{KO}$ mice. Log-rank method was used to compare survival rates between WT and Siglec-$G^{KO}$ mice (**C,D**) Flow cytometric plots (**C**) and quantification of total splenic GC B cell numbers in WT and Siglec-$G^{KO}$ mice (**D**). (**E,F**) Flow cytometric plots (**E**) and quantification of total splenic GC follicular $CD4^+$ T cell numbers in WT and Siglec-$G^{KO}$ mice (**F**). (**G**) Experimental scheme for B cell adoptive transfer into $Rag1^{-/-}$ mice using splenic B cells from aged WT and Siglec-$G^{KO}$ B6 mice. (H) Kaplan-Meier plot showing the survival of recipient mice. Percentage survival of $Rag1^{-/-}$ mice following adoptive transfer of B cells from aged WT and Siglec-$G^{KO}$ donors. (**I**) Quantification of normalized 130 *SIGLEC10* gene count in DLBCL tumors and healthy naive B cells, MBCs, centroblasts (DZ GC), and centrocytes (LZ GC), using a publicly available dataset (accession number: GSE56315) (**J**) Comparison of normalized *SIGLEC10* expression between ABC and GC subgroups of DLBCL tumors in (**I**). (**K,L**) Kaplan-Meier plots of overall (**K**) and event-free (**L**) survival rates of patients with DLBCL tumors expressing low ($SIGLEC10^{lo}$) or high ($SIGLEC10^{hi}$) levels of *SIGLEC10* gene. Log-rank method was used to statistically compare survival rates between the two groups. (**M**) Scheme for the analysis of gene signatures associated in either $SIGLEC10^{lo}$ or $SIGLEC10^{hi}$ DLBCL tumors. Transcriptome data from 562 DLBCL tumors were stratified based on *SIGLEC10* gene expression level. DLBCL in the lower and upper tertiles were to use to identify differentially expressed genes (DEGs) and performed gene set enrichment analysis using annotated Hallmark database as gene signature reference. (**N**) Gene signatures significantly associated with $SIGLEC10^{lo}$ DLBCL. (**O**) Gene signatures significantly associated with $SIGLEC10^{hi}$ DLBCL.

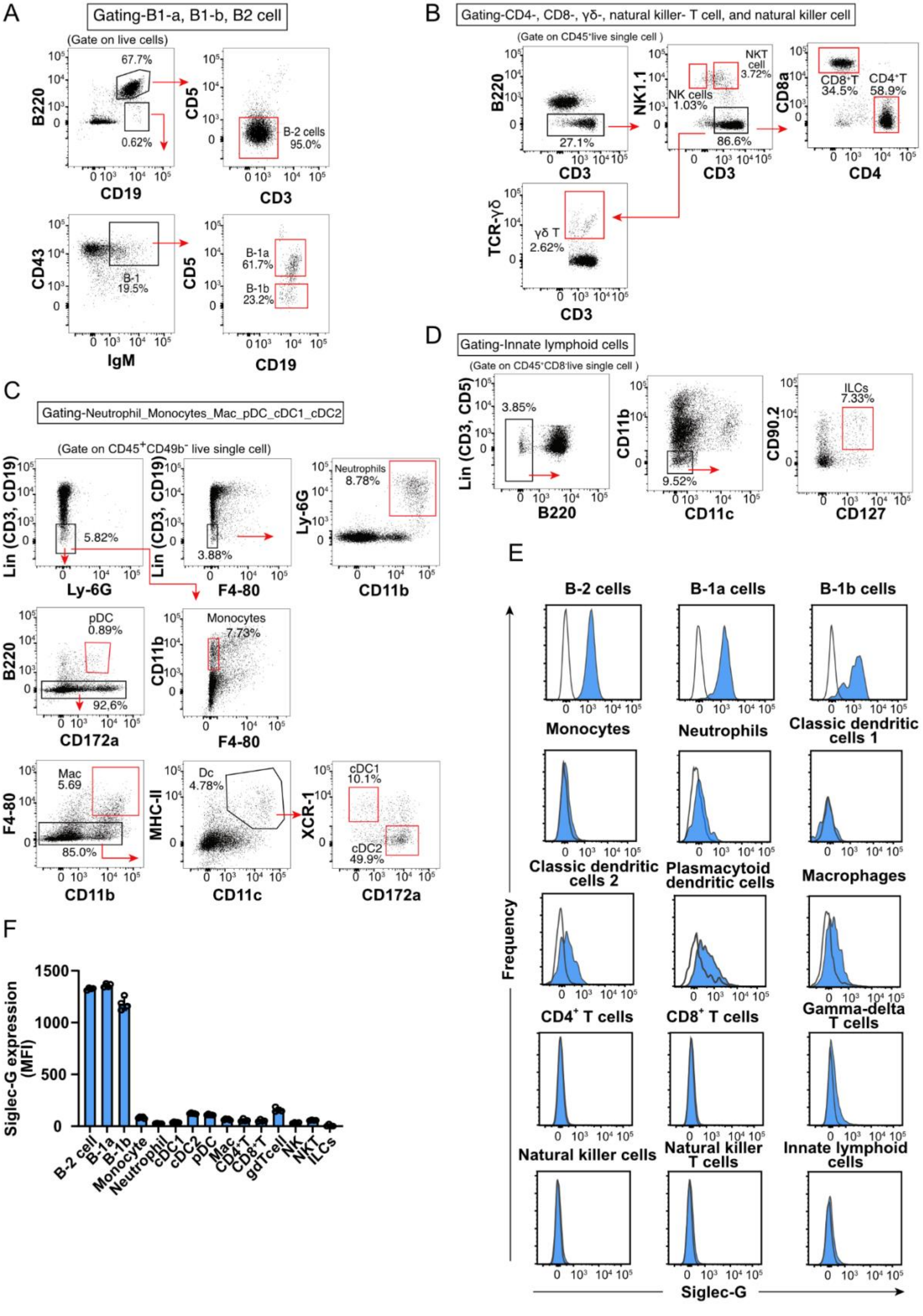
A
Gating-B1-a, B1-b, B2 cell
(Gate on live cells)
B220
CD19
CD5
CD3
B-2 cells
95.0%
67.7%
0.62%
CD43
IgM
B-1
19.5%
B-1a
61.7%
B-1b
23.2%
B
Gating-CD4-, CD8-, γδ-, natural killer- T cell, and natural killer cell
27.1%
NK1.1
NK cells
1.03%
NKT cell
3.72%
86.6%
CD8a
CD4
CD8⁺T
34.5%
CD4⁺T
58.9%
TCR-γδ
γδ T
2.62%
C
Gating-Neutrophil_Monocytes_Mac_pDC_cDC1_cDC2
Lin (CD3, CD19)
Ly-6G
5.82%
F4-80
3.88%
CD11b
Neutrophils
8.78%
CD172a
pDC
0.89%
92.6%
Monocytes
7.73%
Mac
5.69
85.0%
MHC-II
CD11c
Dc
4.78%
XCR-1
cDC1
10.1%
cDC2
49.9%
D
Gating-Innate lymphoid cells
Lin (CD3, CD5)
3.85%
9.52%
CD90.2
CD127
ILCs
7.33%
E
B-2 cells
B-1a cells
B-1b cells
Monocytes
Neutrophils
Classic dendritic cells 1
Classic dendritic cells 2
Plasmacytoid dendritic cells
Macrophages
CD4⁺ T cells
CD8⁺ T cells
Gamma-delta T cells
Natural killer cells
Natural killer T cells
Innate lymphoid cells
Frequency
Siglec-G
F
Siglec-G expression (MFI)
B-2 cell
B-1a
B-1b
Monocyte
Neutrophil
cDC1
cDC2
pDC
Mac
CD4⁺T
CD8⁺T
gdTcell
NK
NKT
ILCs

**Supplementary Figure 4. Siglec-G expression in different immune cell types in naive mouse spleen. (A-D**) Flow cytometric gating strategy for detecting Siglec-G expression on different subsets of B cells (**A**), T and NK cells (**B**), myeloid cells (**C**), and innate lymphoid cells (ILCs) (**D**). (**E**,**F**) Flow cytometric histograms (**E**) and quantification (**F**) of Siglec-G levels on various immune cells.

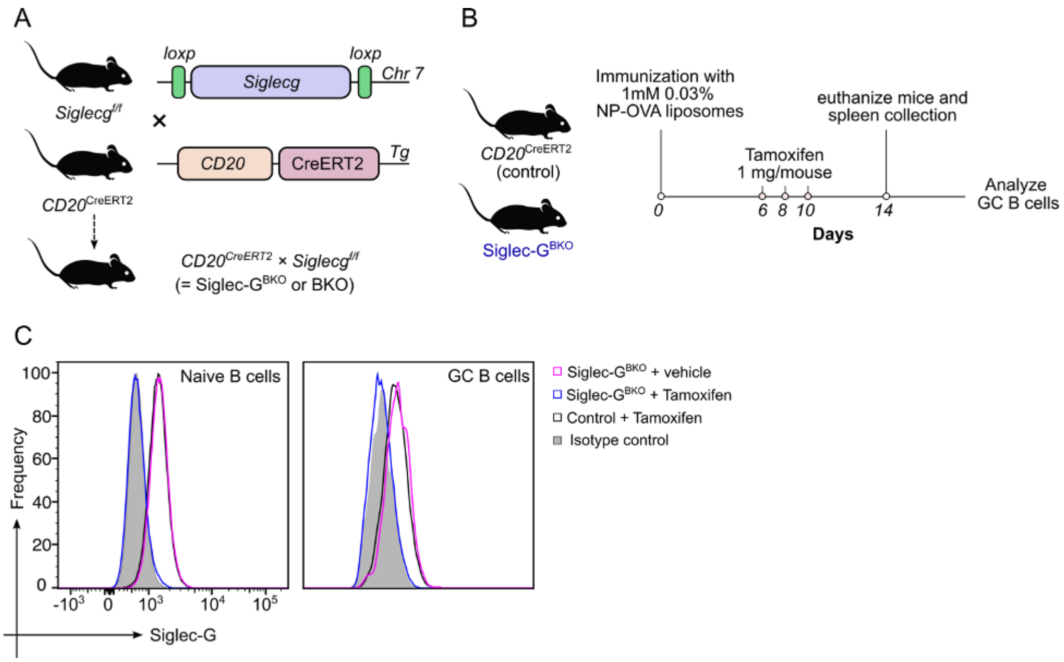


**Supplementary Figure 5. Characterization of a conditional Siglec-G knockout mouse model**. (**A**) Scheme on the generation of a tamoxifen drug-inducible and B cell-specific conditional knockout (Siglec-G$^{BKO}$) mice. (**B**) Experimental scheme for the conditional deletion of Siglec-G on B cells following TD antigen immunization. (**C**) Flow cytometric histograms of Siglec-G expression levels on splenic B cells from immunized control and Siglec-G$^{BKO}$ mice.

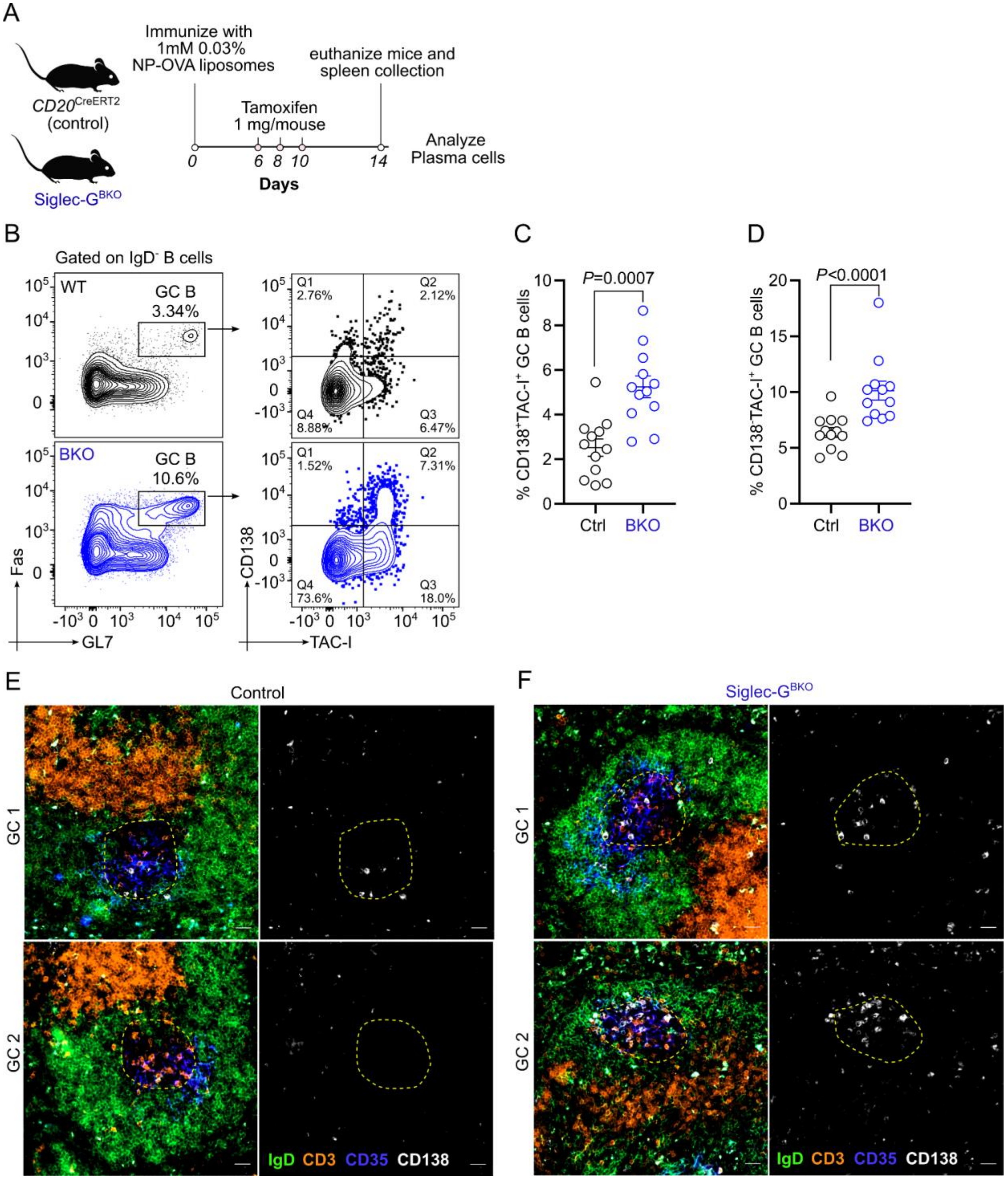


**Supplementary Figure 6. GC-derived plasmablasts are expanded in immunized mice lacking Siglec-G on B cells.** (**A**) Experimental scheme for immunization of control and Siglec-G$^{BKO}$ mice with NP-OVA liposomes. (**B**) Flow cytometric gating strategy for the identification of TACI$^{+}$CD138$^{+}$ plasmablasts within the GC B cell compartment. (**C,D**) Quantifications of the percentage of TACI$^{+}$CD138$^{+}$ plasmablasts on Day 14 (**C**) and Day 21 (**D**) post-immunization. (**E**) Representative IF images of the GC area and represent of GC-derived plasmablasts within the GC.

Data plots are presented as mean±SEM. Statistical analysis for C and D was performed using Kruskal Wallis test.

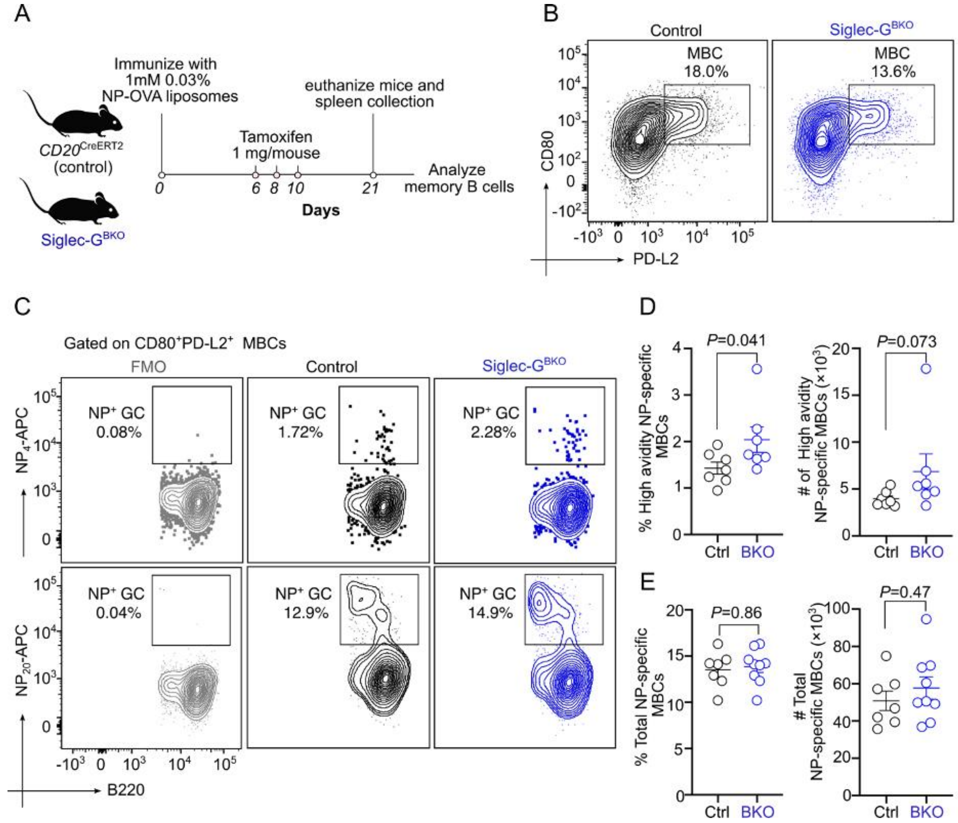


**Supplementary Figure 7. Loss of Siglec-G on B cells does not impact the production of antigen-specific MBCs.** (**A**) Experimental scheme for immunization of control and Siglec-$G^{BKO}$ mice with NP-OVA liposomes. (**B**) Flow cytometric gating strategy for the detection of $CD80^{+}$PD-$L2^{+}$ MBCs. (**C-E**) Flow cytometric plots (**C**) and quantifications of percentages and absolute numbers of high avidity ($NP_4$-$APC^{+}$) (**D**) and total ($NP_{20}$-$APC^{+}$) (**E**) NP-specific GC B cells after day 21 post-immunization. Data plots are presented as mean±SEM. Statistical analysis was performed using Mann-Whitney *U* test.

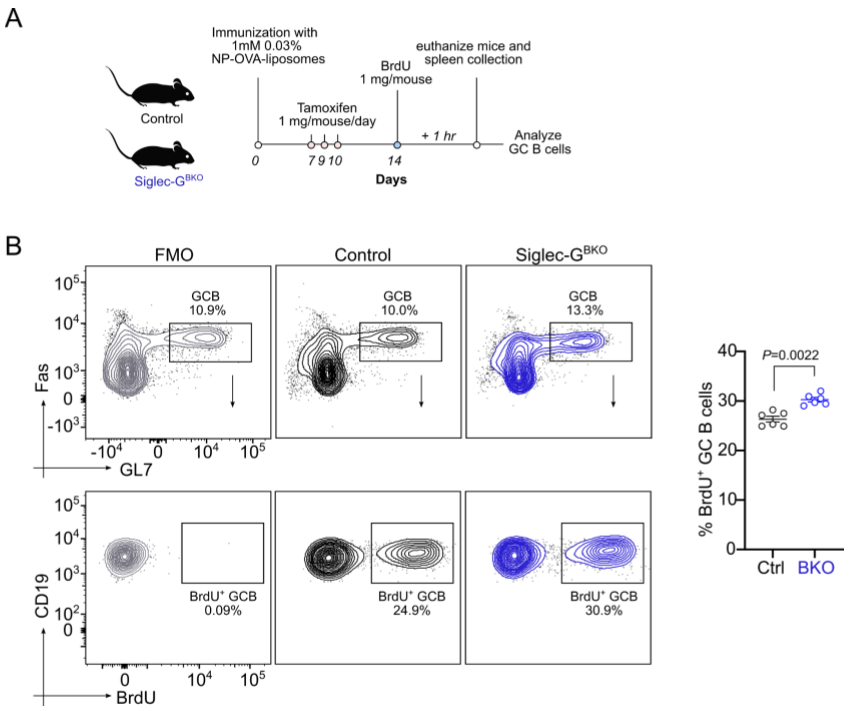


**Supplementary Figure 8. Siglec-G controls GC B cell proliferation.** (**A**) Experimental scheme quantification of proliferating GC B cells using BrdU. Day 14 after NP-OVA liposome immunization, control and Siglec-$G^{BKO}$ mice were treated with 1 mg BrdU/mouse. One hour later, mice were euthanized, and spleens were harvested to analyze for dividing GC B cells. (**B**) Flow cytometric and quantification plots of $BrdU^+$ GC B cells. Data plots are presented as mean±SEM. Statistical analysis was performed using Mann-Whitney *U* test.

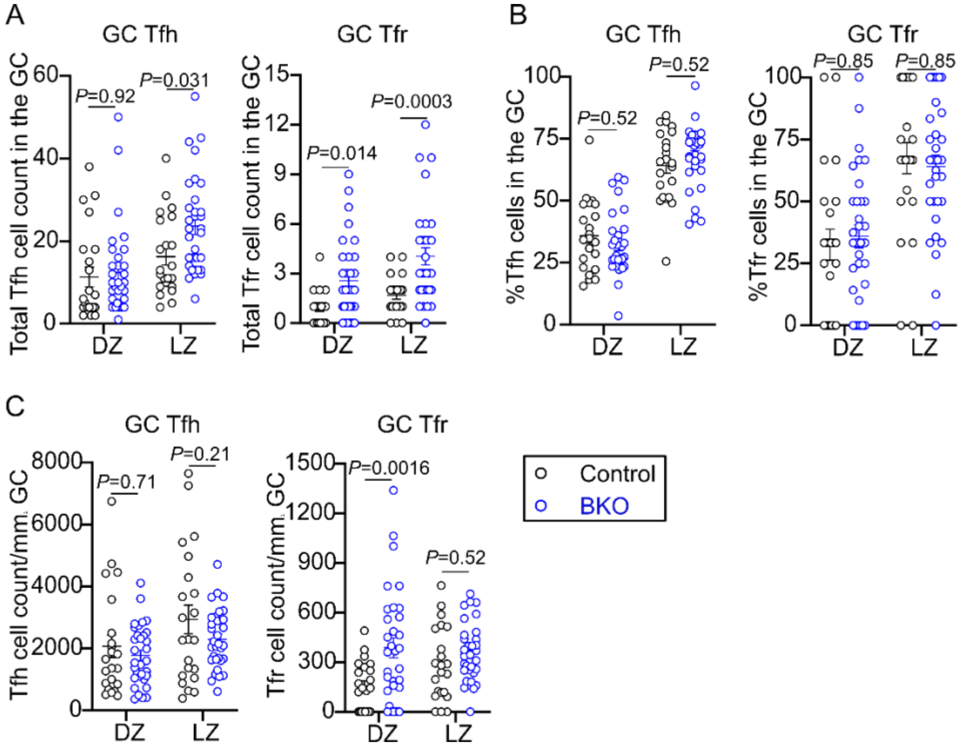


**Supplementary Figure 9. Loss of Siglec-G on B cells does not impact Tfh and Tfr distribution in the GC.** (**A**) Quantifications of absolute Tfh ($Foxp3^{-}CD3^{+}$) and Tfr ($Foxp3^{+}CD3^{+}$) cell count in the GC from spleens of immunized control and Siglec-$G^{BKO}$ mice on Day 14 post-immunization. (**B**) Quantifications of percent Tfh cells and Tfr cells in the GC from spleens of immunized control and Siglec-$G^{BKO}$ mice. (**C**) Quantification of Tfh and Tfr cell count/$mm^2$ GC from spleens of immunized control and Siglec-$G^{BKO}$ mice. Data plots are presented as mean±SEM. Statistical analysis was performed using two-way ANOVA with post-hoc Sidak's multiple comparison's test.

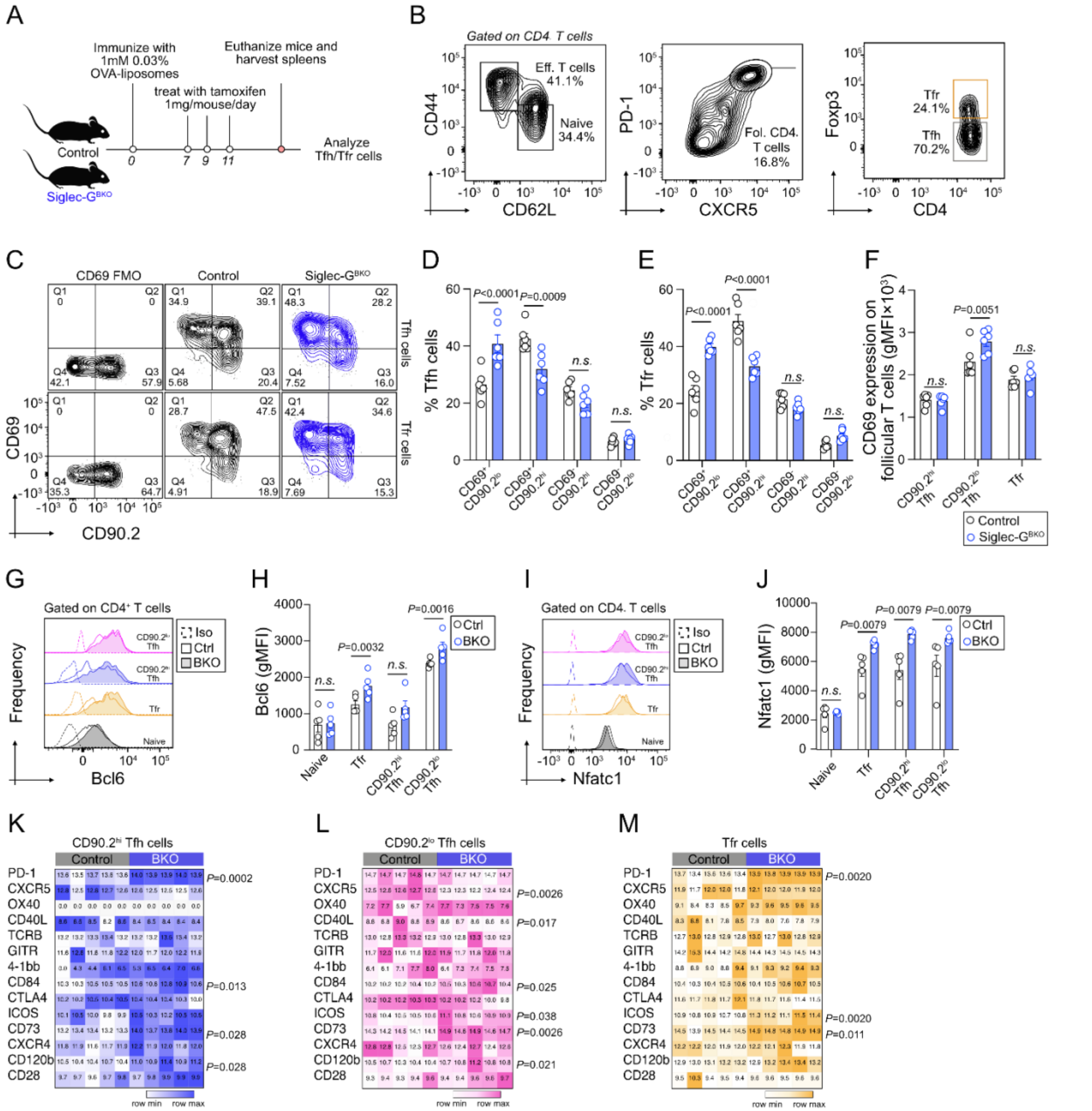


**Supplementary Figure 10. Siglec-G on GC B cells restricts follicular CD4$^+$ T cell activation.** (**A**) Experimental scheme for immunization of control and Siglec-G$^{BKO}$ mice with OVA liposomes. (**B**) Flow cytometric gating strategy for the detection of Tfh and Tfr. (**C-E**) Flow cytometric plots (**C**) and quantifications of percentages (**D,E**) of CD69$^+$ or CD69$^-$ on CD90.2$^+$ and CD90.2$^-$ Tfh. (**F**) The quantification plot of CD69 expression on CD90.2$^{hi}$ Tfh, CD90.2$^{lo}$ Tfh, and Tfr. (**G,H**) Flow cytometric histograms (**G**) and quantification (**H**) of Bcl6 levels in naive and follicular CD4$^+$ T cells. (**I,J**) Flow cytometric histograms (**I**) and quantification (**J**) of Nfatc1 levels in naive and follicular CD4$^+$ T cells. (**K-M**) Heatmaps of gMFIs of key receptors expressed on follicular CD4$^+$ T cells. Data plots are presented as mean±SEM. Statistical analysis was performed using two-way

ANOVA with post-hoc Sidak's multiple comparison's test for D, E, F, H, and J, and unpaired *t* tests for K, L, and M.

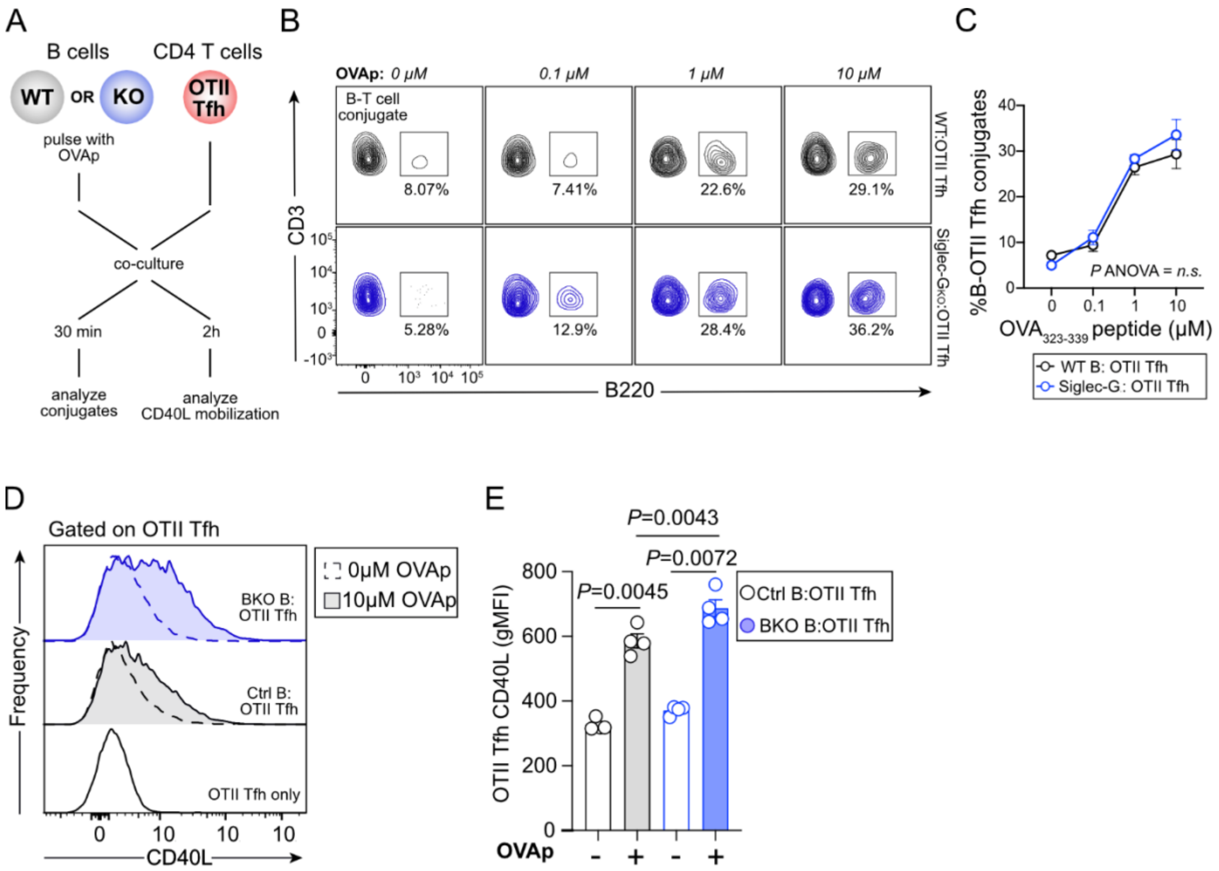


**Supplementary Figure 11. Siglec-G regulates CD40L surface mobilization.** (**A**) For the B-T conjugation assay, purified B cells from control (Ctrl) and full-body Siglec-G knockout (KO) mice were pulsed with varying concentrations of $OVA_{323-339}$ peptide (0 to 10 μM) for 30 min at 37°C, followed by a 30-minute co-culture. For the CD40L mobilization test, purified B cells from Ctrl and B cell-specific knockout (BKO) mice were pulsed with peptide for 2 hours at 37°C, followed by a 2-hour co-culture. In both assays, the pulsed B cells were co-cultured at a 3:1 (B:T) ratio with purified $CD4^{+}$ T cells isolated from OT-II transplanted and immunized B6-$CD45.1^{+}$ mice.(**B**) Flow cytometric gating strategy for the stable B-T cell conjugates identified as double-positive events ($B220^{+}$, $CD3^{+}$). (**C**) Quantification of the percentage of B-T cell conjugates formed between the Tfh cells and the respective B cell genotypes. (**D,E**) Flow cytometric histograms (**D**) and quantification (**E**) of surface CD40L levels on OT-II Tfh cells co-cultured with either Ctrl or BKO B cells. Data plots are presented as mean±SEM. Statistical analysis was performed using one-way ANOVA followed by Tukey's test for multiple comparisons.

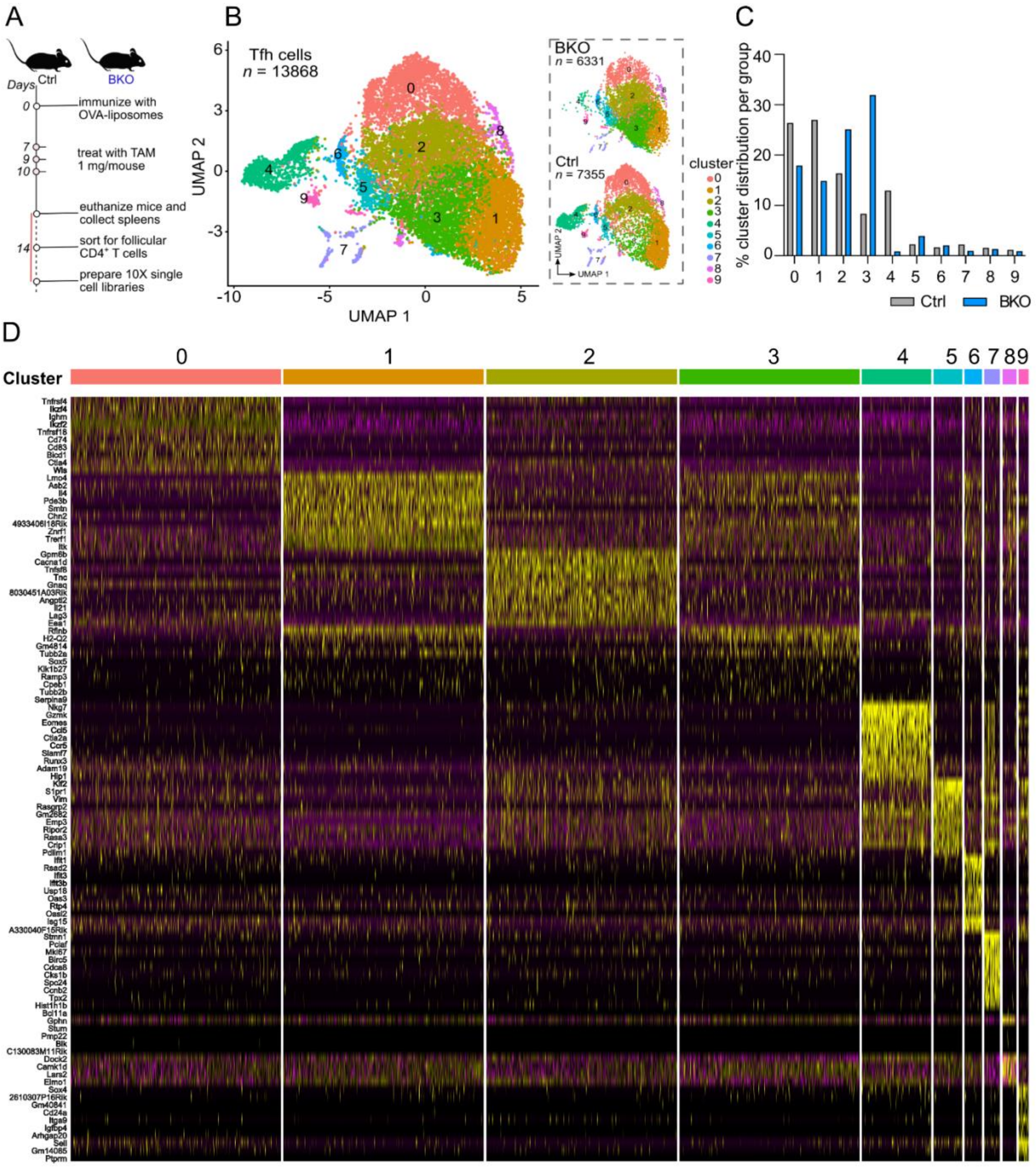


**Supplementary Figure 12. Single cell RNA sequencing of Tfh cells.** (**A**) Experimental workflow for control and Siglec-G$^{BKO}$ mice were immunized with OVA-liposomes following the Tamoxifen treatment (TAM) on days 7, 9, and 10 to induce B cell-specific deletion. Follicular CD4$^{+}$ T cells were sorted from spleens at day 14 and processed for single-cell RNA sequencing. (**B**) Transcriptional heterogeneity of Tfh cells in UMAP visualization of 13,868 Tfh cells colored by cluster (left) and split by genotype of control (Ctrl) and Siglec-G$^{BKO}$(BKO) (right panel). (**C**) Siglec-G$^{BKO}$-driven shift in Tfh cluster distribution with quantification of the percentage distribution of Tfh cells across clusters 0–9. (**D**) Heatmap displaying the top 10 differentially expressed genes identified in each Tfh Seurat cluster. Relative gene expression is row-scaled, with yellow indicating maximum relative expression and dark purple indicating minimum relative expression.

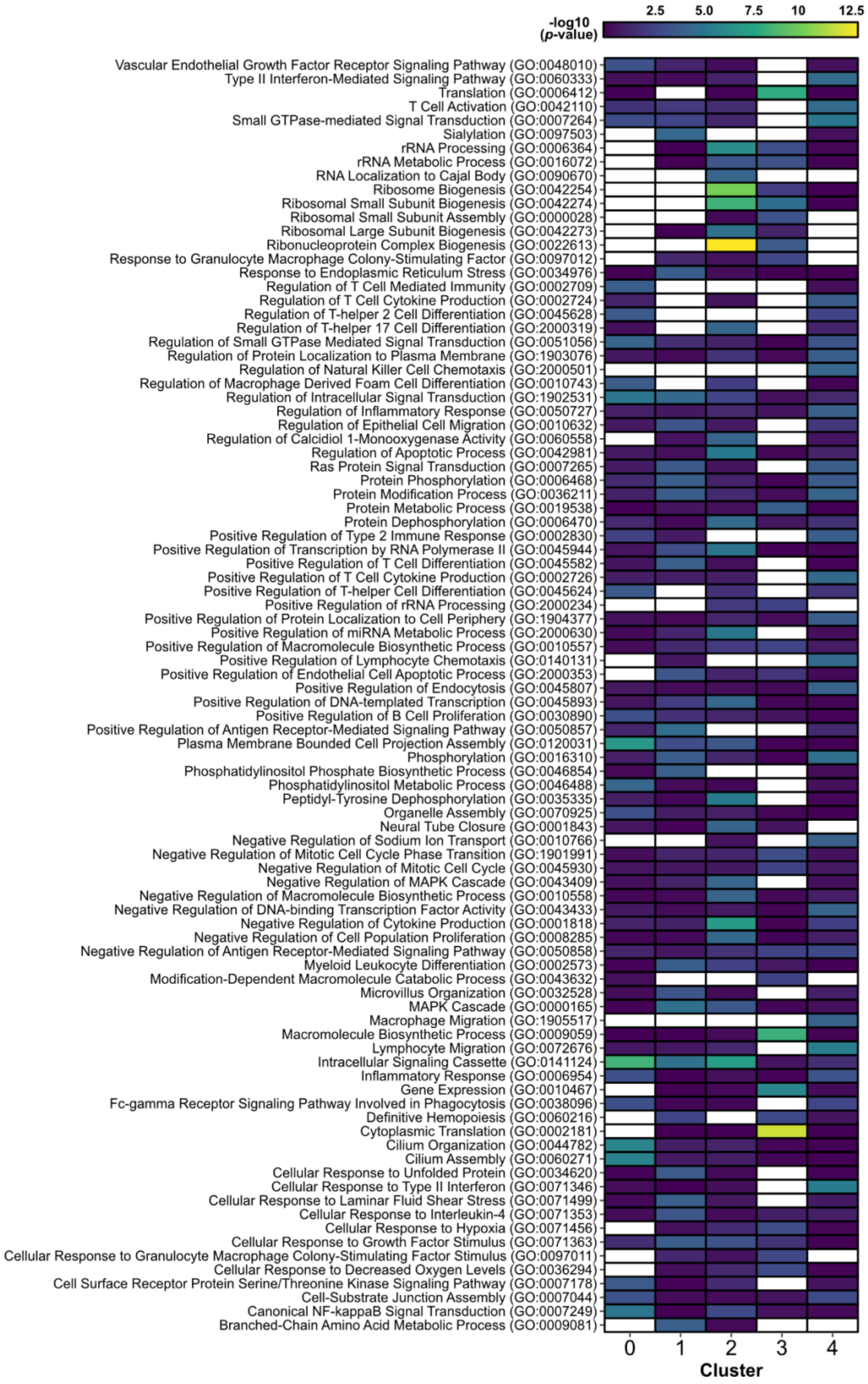
-log10 (p-value)
2.5
5.0
7.5
10
12.5
Vascular Endothelial Growth Factor Receptor Signaling Pathway (GO:0048010)
Type II Interferon-Mediated Signaling Pathway (GO:0060333)
Translation (GO:0006412)
T Cell Activation (GO:0042110)
Small GTPase-mediated Signal Transduction (GO:0007264)
Sialylation (GO:0097503)
rRNA Processing (GO:0006364)
rRNA Metabolic Process (GO:0016072)
RNA Localization to Cajal Body (GO:0090670)
Ribosome Biogenesis (GO:0042254)
Ribosomal Small Subunit Biogenesis (GO:0042274)
Ribosomal Small Subunit Assembly (GO:0000028)
Ribosomal Large Subunit Biogenesis (GO:0042273)
Ribonucleoprotein Complex Biogenesis (GO:0022613)
Response to Granulocyte Macrophage Colony-Stimulating Factor (GO:0097012)
Response to Endoplasmic Reticulum Stress (GO:0034976)
Regulation of T Cell Mediated Immunity (GO:0002709)
Regulation of T Cell Cytokine Production (GO:0002724)
Regulation of T-helper 2 Cell Differentiation (GO:0045628)
Regulation of T-helper 17 Cell Differentiation (GO:2000319)
Regulation of Small GTPase Mediated Signal Transduction (GO:0051056)
Regulation of Protein Localization to Plasma Membrane (GO:1903076)
Regulation of Natural Killer Cell Chemotaxis (GO:2000501)
Regulation of Macrophage Derived Foam Cell Differentiation (GO:0010743)
Regulation of Intracellular Signal Transduction (GO:1902531)
Regulation of Inflammatory Response (GO:0050727)
Regulation of Epithelial Cell Migration (GO:0010632)
Regulation of Calcidiol 1-Monooxygenase Activity (GO:0060558)
Regulation of Apoptotic Process (GO:0042981)
Ras Protein Signal Transduction (GO:0007265)
Protein Phosphorylation (GO:0006468)
Protein Modification Process (GO:0036211)
Protein Metabolic Process (GO:0019538)
Protein Dephosphorylation (GO:0006470)
Positive Regulation of Type 2 Immune Response (GO:0002830)
Positive Regulation of Transcription by RNA Polymerase II (GO:0045944)
Positive Regulation of T Cell Differentiation (GO:0045582)
Positive Regulation of T Cell Cytokine Production (GO:0002726)
Positive Regulation of T-helper Cell Differentiation (GO:0045624)
Positive Regulation of rRNA Processing (GO:2000234)
Positive Regulation of Protein Localization to Cell Periphery (GO:1904377)
Positive Regulation of miRNA Metabolic Process (GO:2000630)
Positive Regulation of Macromolecule Biosynthetic Process (GO:0010557)
Positive Regulation of Lymphocyte Chemotaxis (GO:0140131)
Positive Regulation of Endothelial Cell Apoptotic Process (GO:2000353)
Positive Regulation of Endocytosis (GO:0045807)
Positive Regulation of DNA-templated Transcription (GO:0045893)
Positive Regulation of B Cell Proliferation (GO:0030890)
Positive Regulation of Antigen Receptor-Mediated Signaling Pathway (GO:0050857)
Plasma Membrane Bounded Cell Projection Assembly (GO:0120031)
Phosphorylation (GO:0016310)
Phosphatidylinositol Phosphate Biosynthetic Process (GO:0046854)
Phosphatidylinositol Metabolic Process (GO:0046488)
Peptidyl-Tyrosine Dephosphorylation (GO:0035335)
Organelle Assembly (GO:0070925)
Neural Tube Closure (GO:0001843)
Negative Regulation of Sodium Ion Transport (GO:0010766)
Negative Regulation of Mitotic Cell Cycle Phase Transition (GO:1901991)
Negative Regulation of Mitotic Cell Cycle (GO:0045930)
Negative Regulation of MAPK Cascade (GO:0043409)
Negative Regulation of Macromolecule Biosynthetic Process (GO:0010558)
Negative Regulation of DNA-binding Transcription Factor Activity (GO:0043433)
Negative Regulation of Cytokine Production (GO:0001818)
Negative Regulation of Cell Population Proliferation (GO:0008285)
Negative Regulation of Antigen Receptor-Mediated Signaling Pathway (GO:0050858)
Myeloid Leukocyte Differentiation (GO:0002573)
Modification-Dependent Macromolecule Catabolic Process (GO:0043632)
Microvillus Organization (GO:0032528)
MAPK Cascade (GO:0000165)
Macrophage Migration (GO:1905517)
Macromolecule Biosynthetic Process (GO:0009059)
Lymphocyte Migration (GO:0072676)
Intracellular Signaling Cassette (GO:0141124)
Inflammatory Response (GO:0006954)
Gene Expression (GO:0010467)
Fc-gamma Receptor Signaling Pathway Involved in Phagocytosis (GO:0038096)
Definitive Hemopoiesis (GO:0060216)
Cytoplasmic Translation (GO:0002181)
Cilium Organization (GO:0044782)
Cilium Assembly (GO:0060271)
Cellular Response to Unfolded Protein (GO:0034620)
Cellular Response to Type II Interferon (GO:0071346)
Cellular Response to Laminar Fluid Shear Stress (GO:0071499)
Cellular Response to Interleukin-4 (GO:0071353)
Cellular Response to Hypoxia (GO:0071456)
Cellular Response to Growth Factor Stimulus (GO:0071363)
Cellular Response to Granulocyte Macrophage Colony-Stimulating Factor Stimulus (GO:0097011)
Cellular Response to Decreased Oxygen Levels (GO:0036294)
Cell Surface Receptor Protein Serine/Threonine Kinase Signaling Pathway (GO:0007178)
Cell-Substrate Junction Assembly (GO:0007044)
Canonical NF-kappaB Signal Transduction (GO:0007249)
Branched-Chain Amino Acid Metabolic Process (GO:0009081)
0
1
2
3
4
Cluster

**Supplementary Figure 13. Gene Ontology (GO) enrichment analysis reveals distinct functional specializations within Tfh cell sub-clusters.** Heatmap of significantly enriched GO biological processes associated with the differentially expressed genes upregulated in each of the 5 main Tfh cluster, as described in Figure S12B. The color gradient corresponds to the statistical significance of the GO term in each cluster. White tiles represent GO terms that did not meet the threshold for significant enrichment.

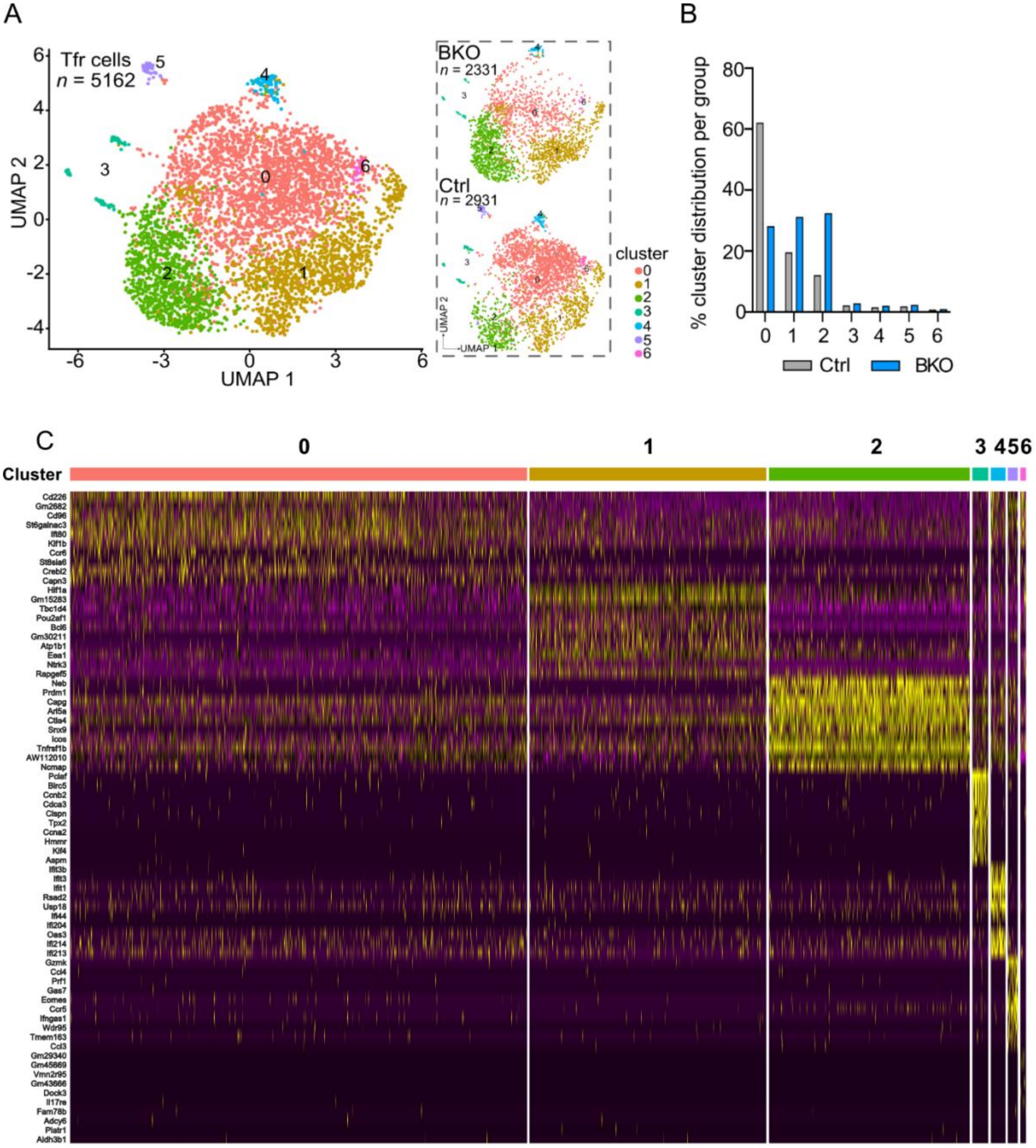


**Supplementary Figure 14. Single cell RNA sequencing of Tfr cells.** (**A**) Transcriptional landscape of Tfr cells in UMAP visualization of 5,162 Tfr cells colored by cluster (left) and split by genotype (right). (**B**) Percentage of Tfr cells per cluster for Ctrl and BKO groups. (**C**) Heatmap illustrating the top differentially expressed marker genes defining distinct Seurat clusters within the Tfr cell compartment from WT and Siglec-G$^{BKO}$ mice. For panel, relative gene expression is row-scaled, with yellow indicating maximum relative expression (upregulation) and dark purple indicating minimum relative expression (downregulation).

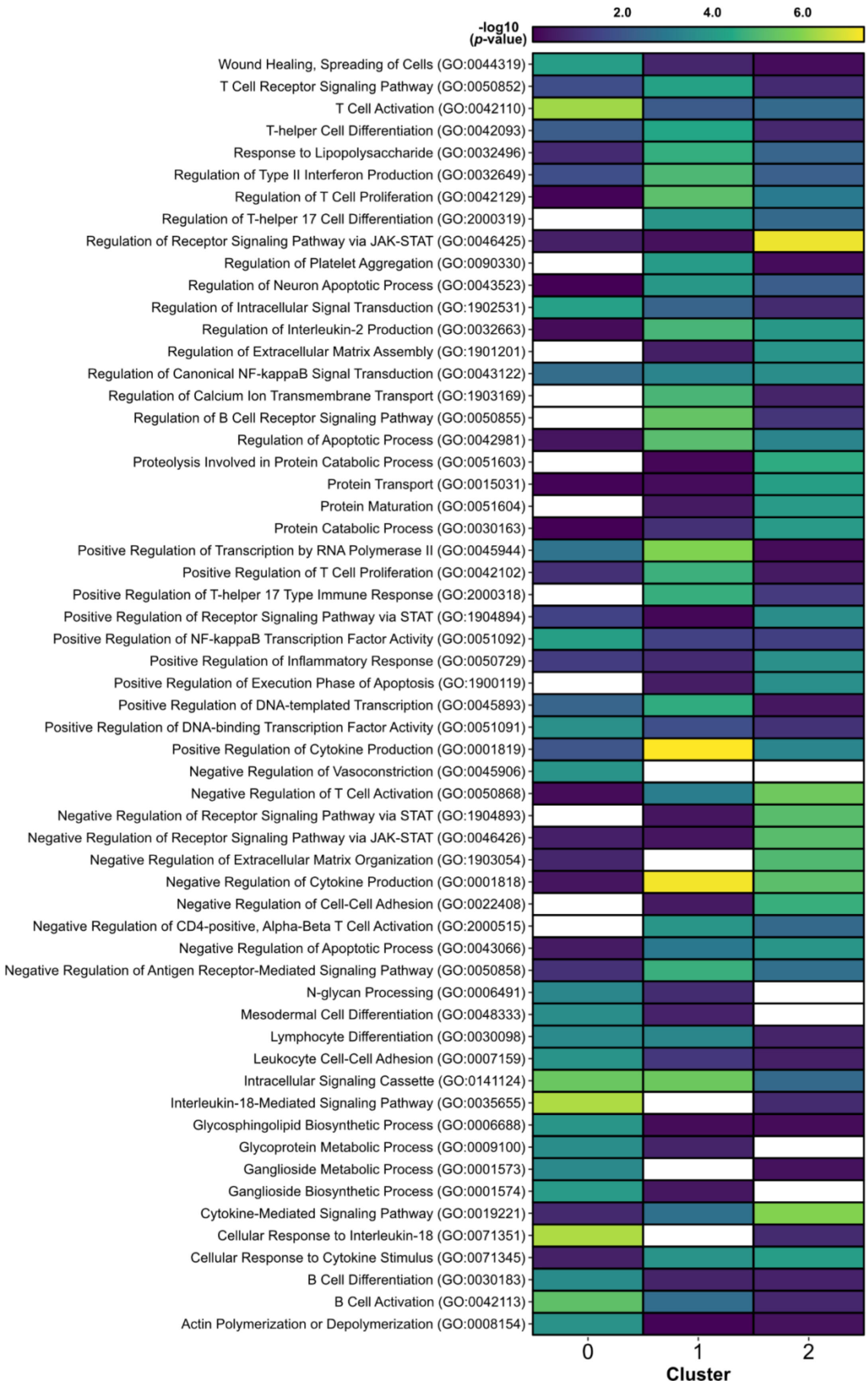
-log10 (p-value)
2.0
4.0
6.0
Wound Healing, Spreading of Cells (GO:0044319)
T Cell Receptor Signaling Pathway (GO:0050852)
T Cell Activation (GO:0042110)
T-helper Cell Differentiation (GO:0042093)
Response to Lipopolysaccharide (GO:0032496)
Regulation of Type II Interferon Production (GO:0032649)
Regulation of T Cell Proliferation (GO:0042129)
Regulation of T-helper 17 Cell Differentiation (GO:2000319)
Regulation of Receptor Signaling Pathway via JAK-STAT (GO:0046425)
Regulation of Platelet Aggregation (GO:0090330)
Regulation of Neuron Apoptotic Process (GO:0043523)
Regulation of Intracellular Signal Transduction (GO:1902531)
Regulation of Interleukin-2 Production (GO:0032663)
Regulation of Extracellular Matrix Assembly (GO:1901201)
Regulation of Canonical NF-kappaB Signal Transduction (GO:0043122)
Regulation of Calcium Ion Transmembrane Transport (GO:1903169)
Regulation of B Cell Receptor Signaling Pathway (GO:0050855)
Regulation of Apoptotic Process (GO:0042981)
Proteolysis Involved in Protein Catabolic Process (GO:0051603)
Protein Transport (GO:0015031)
Protein Maturation (GO:0051604)
Protein Catabolic Process (GO:0030163)
Positive Regulation of Transcription by RNA Polymerase II (GO:0045944)
Positive Regulation of T Cell Proliferation (GO:0042102)
Positive Regulation of T-helper 17 Type Immune Response (GO:2000318)
Positive Regulation of Receptor Signaling Pathway via STAT (GO:1904894)
Positive Regulation of NF-kappaB Transcription Factor Activity (GO:0051092)
Positive Regulation of Inflammatory Response (GO:0050729)
Positive Regulation of Execution Phase of Apoptosis (GO:1900119)
Positive Regulation of DNA-templated Transcription (GO:0045893)
Positive Regulation of DNA-binding Transcription Factor Activity (GO:0051091)
Positive Regulation of Cytokine Production (GO:0001819)
Negative Regulation of Vasoconstriction (GO:0045906)
Negative Regulation of T Cell Activation (GO:0050868)
Negative Regulation of Receptor Signaling Pathway via STAT (GO:1904893)
Negative Regulation of Receptor Signaling Pathway via JAK-STAT (GO:0046426)
Negative Regulation of Extracellular Matrix Organization (GO:1903054)
Negative Regulation of Cytokine Production (GO:0001818)
Negative Regulation of Cell-Cell Adhesion (GO:0022408)
Negative Regulation of CD4-positive, Alpha-Beta T Cell Activation (GO:2000515)
Negative Regulation of Apoptotic Process (GO:0043066)
Negative Regulation of Antigen Receptor-Mediated Signaling Pathway (GO:0050858)
N-glycan Processing (GO:0006491)
Mesodermal Cell Differentiation (GO:0048333)
Lymphocyte Differentiation (GO:0030098)
Leukocyte Cell-Cell Adhesion (GO:0007159)
Intracellular Signaling Cassette (GO:0141124)
Interleukin-18-Mediated Signaling Pathway (GO:0035655)
Glycosphingolipid Biosynthetic Process (GO:0006688)
Glycoprotein Metabolic Process (GO:0009100)
Ganglioside Metabolic Process (GO:0001573)
Ganglioside Biosynthetic Process (GO:0001574)
Cytokine-Mediated Signaling Pathway (GO:0019221)
Cellular Response to Interleukin-18 (GO:0071351)
Cellular Response to Cytokine Stimulus (GO:0071345)
B Cell Differentiation (GO:0030183)
B Cell Activation (GO:0042113)
Actin Polymerization or Depolymerization (GO:0008154)
0
1
2
Cluster

**Supplementary Figure 15. Gene Ontology (GO) enrichment analysis reveals distinct functional specializations within Tfr cell sub-clusters.** Heatmap of significantly enriched GO biological processes associated with the differentially expressed genes upregulated in each of the 3 main Tfr cluster, as described in Figure S14A. The color gradient corresponds to the statistical significance of the GO term in each cluster. White tiles represent GO terms that did not meet the threshold for significant enrichment.

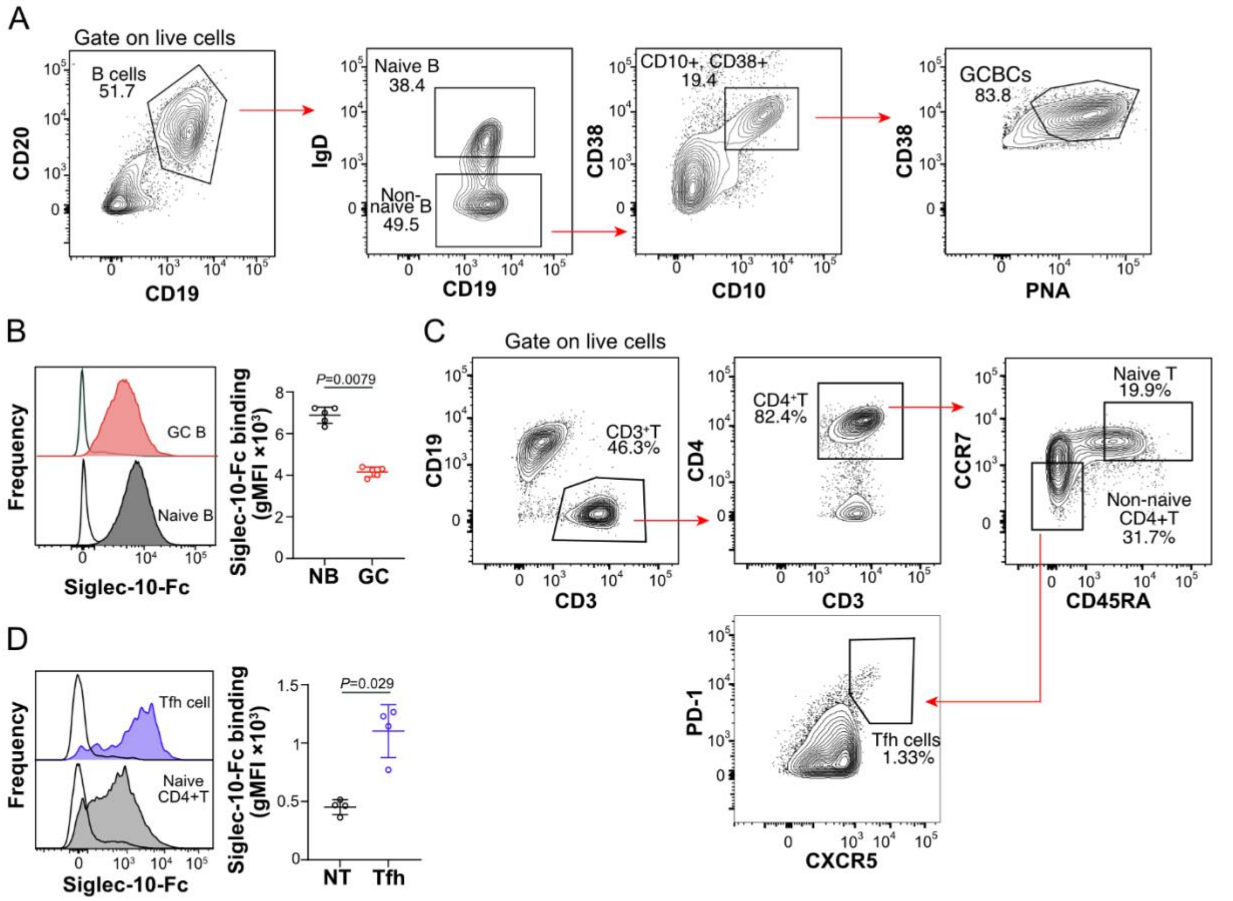


**Supplementary Figure 16. Human GC B cells downregulate ligands for Siglec-10.** (**A**) Flow cytometric gating strategy used to identify naive and GC B cells from human tonsils. (**B**) The flow cytometry histograms and quantification of Siglec-10-Fc binding between human naive B cells and GC B cells. (**C**) Flow cytometric gating strategy used to identify Tfh cells from human tonsils. (**D**) Representative flow cytometry histogram and quantification of Siglec-10-Fc binding on human Tfh cells. Data plots are presented as mean±SEM. Statistical analysis in B and D was performed using Mann-Whitney *U* test.

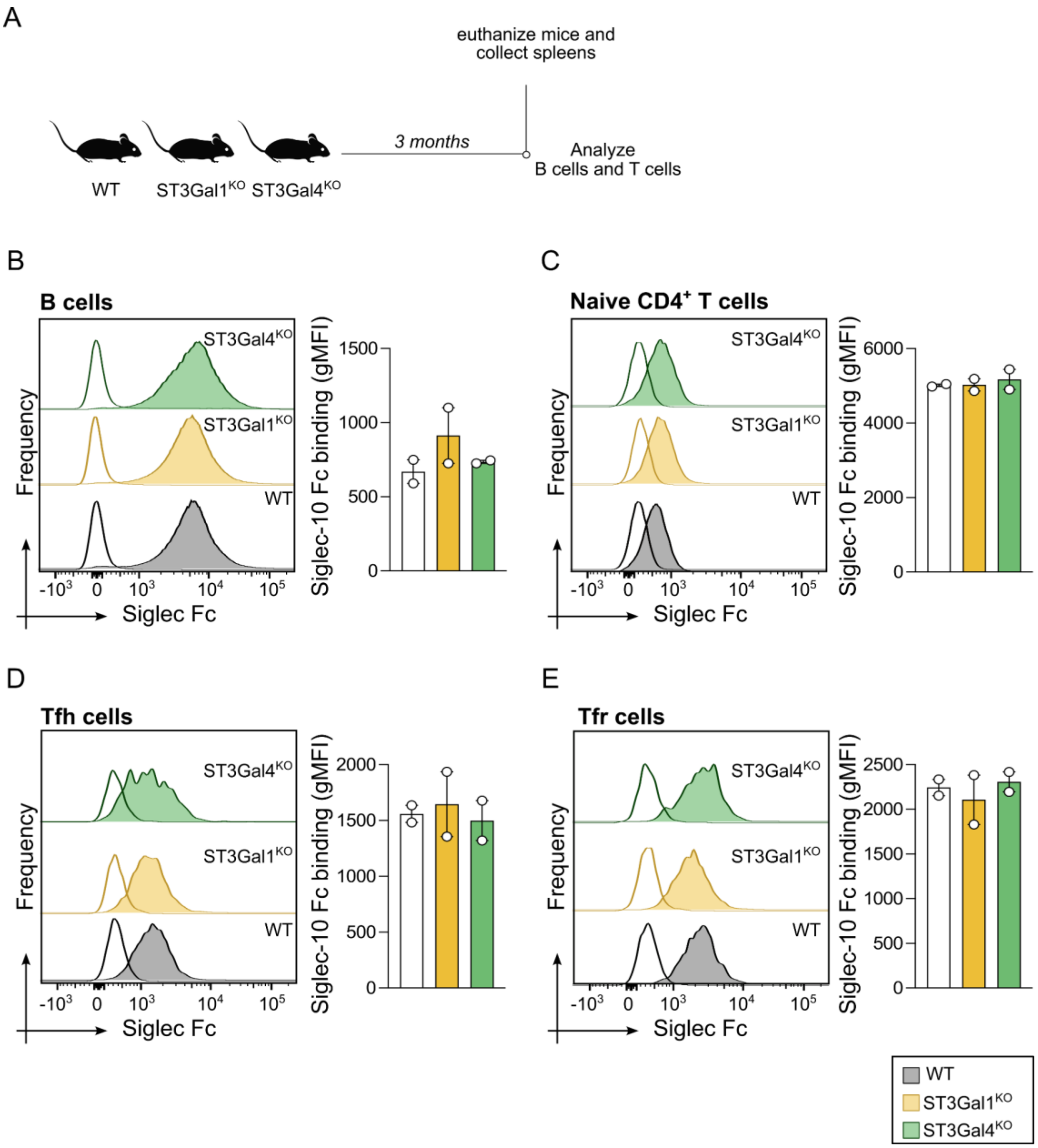


**Supplementary Figure 17. Siglec-10 ligands on mouse B and T cells from a single knockout of the ST3Gal1 and ST3Gal4 sialyltransferases.** (**A**) Experimental scheme for the Siglec-10-Fc staining on sialyltransferase knockout mice. (**B-E**) Siglec10-Fc staining on splenocytes from ST3Gal1$^{KO}$, ST3Gal4$^{KO}$, and WT mice. Flow cytometry histogram (*left*) and quantification (*right*) are shown for B cells (**B**), Naive CD4$^{+}$T cells (**C**), Tfh cells (**D**), and Tfr cells (**E**).

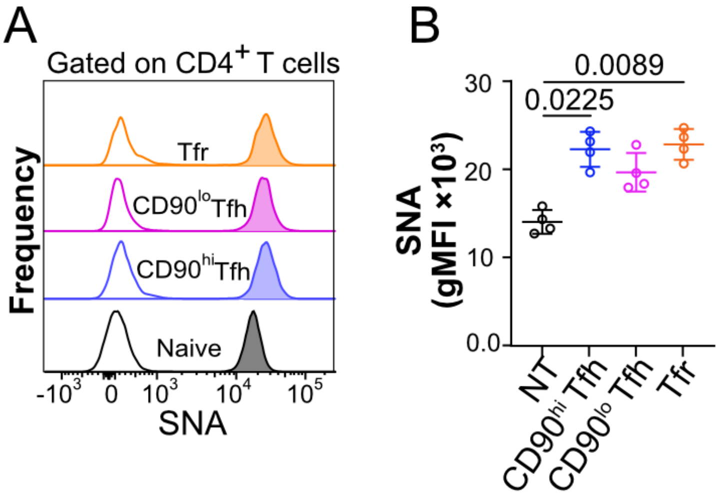


**Supplementary Figure 18. Follicular CD4$^+$ T cells upregulate α2-6 linked sialic acid glycans**. (**A,B**)Flow cytometry histograms (**A**) and quantification (**B**) of *Sambucus nigra* (SNA) lectin staining on naive CD4$^+$ T and follicular CD4$^+$ T cells. Data plots are presented as mean±SEM. Statistical analysis was performed using a Kruskal-Wallis test with post-hoc Tukey's test for multiple comparisons.

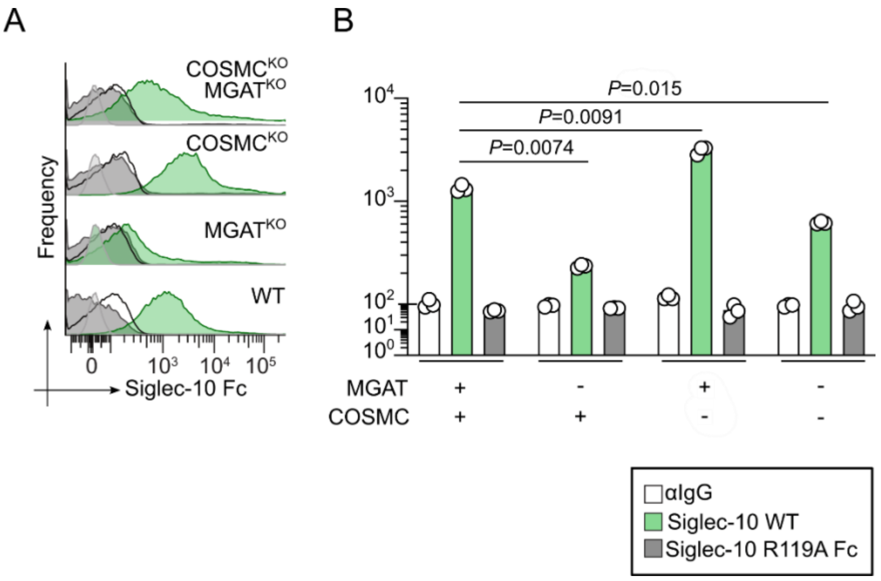


**Supplementary Figure 19. Siglec-10 binds preferentially to sialylated *N*-glycans.** (**A**) Flow histograms of Siglec-10-Fc binding on WT, $MGAT^{KO}$, $COSMC^{KO}$ and $MGAT^{KO}\times COSMC^{KO}$ U937 cell lines. (**B**) Quantification of Siglec-10-Fc binding (MFI) on WT, $MGAT^{KO}$, $COSMC^{KO}$ and $MGAT^{KO}\times COSMC^{KO}$ U937 cell lines. Data plots are presented as mean±SEM. Statistical analysis was performed using Mann-Whitney *U* test.

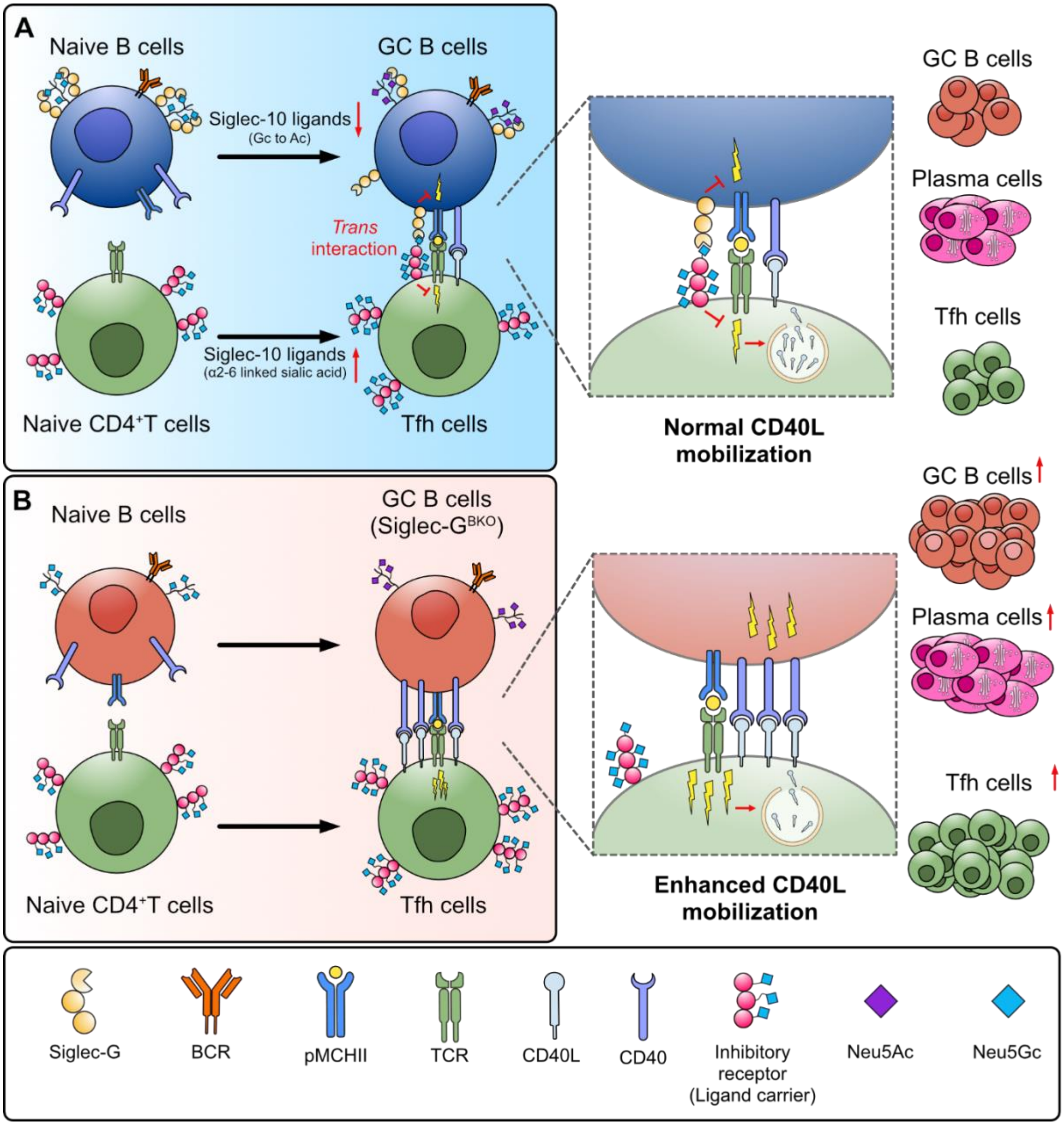


**Supplementary Figure 20. Glycan remodeling in the GC promotes *trans* interactions of Siglec-G/10 to modulate B-T cell interactions.** (**A**) Intrinsic glycan remodeling on GC B cells and Tfh cells facilitates Siglec-10 *trans* interactions. GC B cells downregulate *cis* ligands for Siglec-10 partly due to switch from Neu5Gc to Neu5Ac sialic acid. Concurrently, Tfh cells upregulate Siglec-G/10 glycan ligands by increasing α2-6 linked sialylation. Engagement of Siglec-G/10 on GC B cells and its *trans* ligands on Tfh cells during contact likely results in the recruitment of immunomodulatory co-receptors close to the TCR complex, thereby fine-tuning Tfh activation strength and GC output. (**B**) Loss of Siglec-G/10 on B cells drives an enhanced GC reaction. Without Siglec-G/10, bidirectional signaling between GC B cells and Tfh cells is augmented leading increase GC B cell numbers, improved GC localization of Tfh cells, and higher PC output. Some illustration from NIAID NIH BioArt Source.

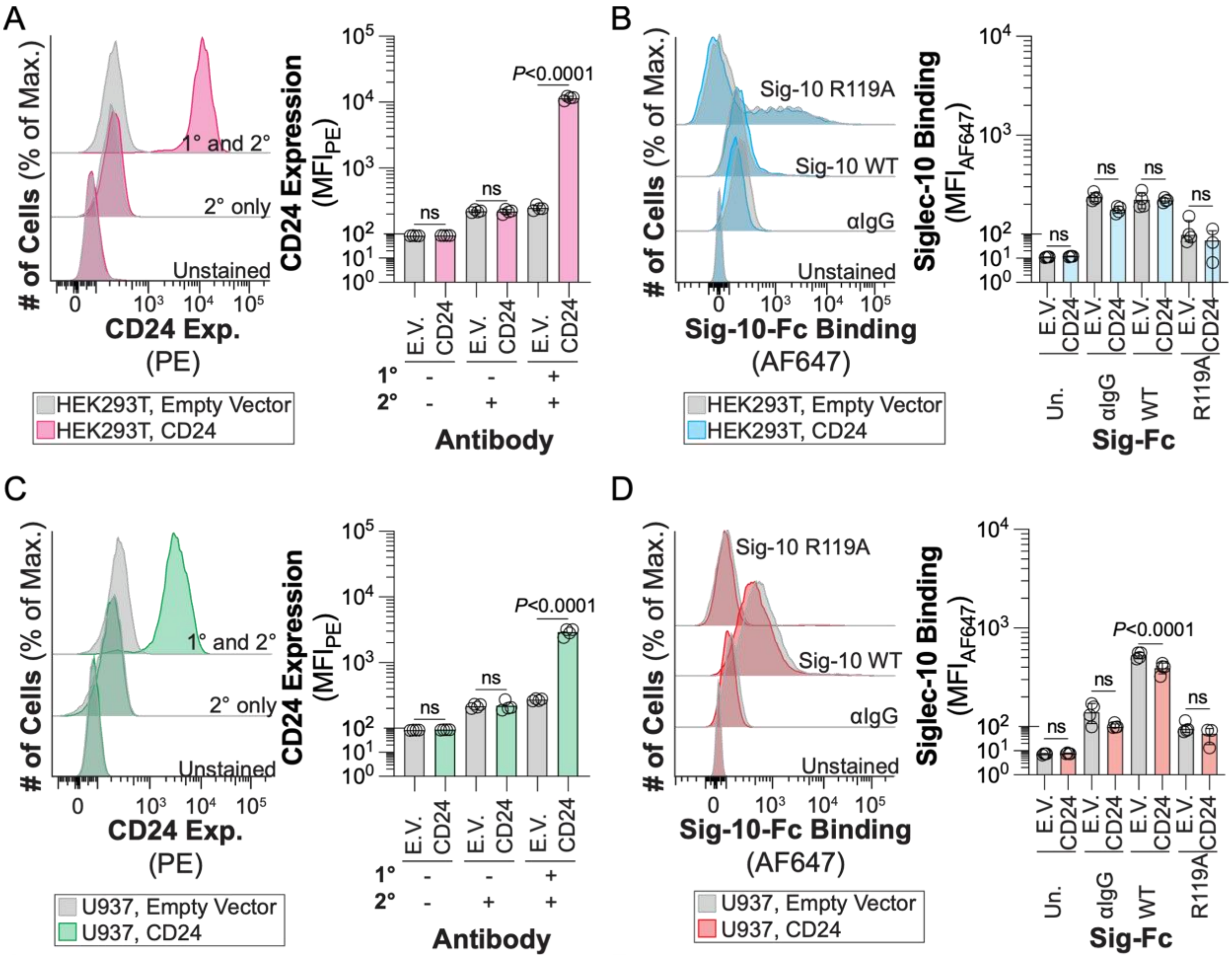


**Supplementary Figure 21. Siglec-10-Fc binding is independent of CD24 expression in HEK293T and U937 cells.** (**A,B**) Flow cytometric analysis quantifying CD24 expression and Siglec-10-Fc staining on HEK293T cells transfected with either an empty vector (E.V.) or a CD24 overexpression vector (CD24). (**C,D**) Flow cytometric analysis quantifying CD24 expression and Siglec-10-Fc staining on U937 cells transduced with E.V. and CD24. Data plots are presented as mean±SEM. Statistical analysis was performed using one-way ANOVA followed by Tukey's test for multiple comparisons.

**Table S1. Differential gene expression analysis identifying cluster-specific marker genes for follicular T cell subsets.**

See attached Excel file

**Table S2. Enriched proteins identified by APEX2-based proximity labeling assay.** The table summarizes the 69 candidate interacting proteins of Siglec-10 on follicular CD4$^+$ T cells identified by APEX2-based proximity labeling and quantitative mass spectrometry.

| | Accession | Gene Symbol | log2 (FC) | Abundance Ratio *P*-Value: (Sample) / (Control) | | Accession | Gene Symbol | log2 (FC) | Abundance Ratio *P*-Value: (Sample) / (Control) |
|---|---|---|---|---|---|---|---|---|---|
| | | | | | 35 | Q8QZY6 | Tspan14 | 2.47 | 0.0006 |
| | | | | | 36 | O09126 | Sema4d | 2.46 | 0.0038 |
| 1 | A0A075B5J4 | Trbc2 | 4.35 | 0.0001 | 37 | Q91V08 | Clec2d | 2.43 | 0.0001 |
| 2 | P47774 | Ccr7 | 4.07 | 0.0213 | 38 | B1ASL3 | Tnfrsf4 | 2.42 | 0.0008 |
| 3 | P97333 | Nrp1 | 3.80 | 0.0447 | 39 | A0A075B5J5 | Trbv31 | 2.42 | 0.0029 |
| 4 | P04212 | TVB4 | 3.61 | 0.0137 | 40 | Q80V42 | Cpm | 2.40 | 0.0016 |
| 5 | P41272 | Cd27 | 3.56 | 0.0003 | 41 | E9QNY6 | Btla | 2.38 | 0.0013 |
| 6 | P97370 | Atp1b3 | 3.44 | 0.0028 | 42 | Q01965 | Ly9 | 2.37 | 0.0006 |
| 7 | P35330 | Icam2 | 3.37 | 0.0003 | 43 | P01831 | Thy1 | 2.36 | 0.0030 |
| 8 | P01887 | B2m | 3.33 | 0.0000 | 44 | P26011 | Itgb7 | 2.30 | 0.0032 |
| 9 | B1B507 | Sell | 3.31 | 0.0000 | 45 | Q62312 | Tgfbr2 | 2.25 | 0.0028 |
| 10 | Q61003 | Cd6 | 3.16 | 0.0002 | 46 | Q9DBX3 | Susd2 | 2.24 | 0.0022 |
| 11 | Q3UXS0 | Scamp3 | 3.11 | 0.0363 | 47 | P14206 | Rpsa | 2.18 | 0.0008 |
| 12 | Q9Z127 | Slc7a5 | 3.11 | 0.0022 | 48 | P10852 | Slc3a2 | 2.15 | 0.0158 |
| 13 | Q00651 | Itga4 | 3.11 | 0.0001 | 49 | Q62351 | Tfrc | 2.07 | 0.0002 |
| 14 | Q9ET39 | Slamf6 | 3.10 | 0.0034 | 50 | P11688 | Itga5 | 1.94 | 0.0022 |
| 15 | F8WJA1 | Fxyd5 | 3.09 | 0.0007 | 51 | Q64697 | Ptprcap | 1.82 | 0.0402 |
| 16 | P06800 | Ptprc | 3.06 | 0.0002 | 52 | O88713 | Klrg1 | 1.74 | 0.0036 |
| 17 | P09055 | Itgb1 | 3.02 | 0.0006 | 53 | Q61503 | Nt5e | 1.73 | 0.0160 |
| 18 | P31041 | Cd28 | 3.01 | 0.0002 | 54 | A2AW86 | Ly75 | 1.67 | 0.0056 |
| 19 | Q64735 | Cr1l | 2.99 | 0.0091 | 55 | P04214 | TVB6 | 1.65 | 0.0003 |
| 20 | A2APM2 | Cd44 | 2.95 | 0.0002 | 56 | G3UZP7 | H2-D1 | 1.58 | 0.0117 |
| 21 | D3Z627 | Itgal | 2.90 | 0.0002 | 57 | P22646 | Cd3e | 1.58 | 0.0078 |
| 22 | B1ARB3 | Pecam1 | 2.87 | 0.0013 | 58 | P01901 | H2-K1 | 1.55 | 0.0015 |
| 23 | P14094 | Atp1b1 | 2.82 | 0.0001 | 59 | B1ASL6 | Tnfrsf18 | 1.52 | 0.0294 |
| 24 | P43406 | Itgav | 2.81 | 0.0002 | 60 | P06332 | Cd4 | 1.45 | 0.0130 |
| 25 | P11835 | Itgb2 | 2.80 | 0.0002 | 61 | Q921F4 | Hnrnpll | 1.32 | 0.0040 |
| 26 | A0A171EBK7 | Vsir | 2.77 | 0.0160 | 62 | Q6PFB2 | Rcc1 | 1.29 | 0.0009 |
| 27 | P21995 | Emb | 2.75 | 0.0009 | 63 | P13379 | Cd5 | 1.20 | 0.0348 |
| 28 | P18181 | Cd48 | 2.74 | 0.0009 | 64 | Q9Z2A9 | Ggt5 | 1.17 | 0.0072 |
| 29 | A0A0R4J090 | Cd1d1 | 2.67 | 0.0017 | 65 | Q9D0L7 | Armc10 | 1.12 | 0.0344 |
| 30 | P08920 | Cd2 | 2.67 | 0.0148 | 66 | P29699 | Ahsg | 1.11 | 0.0085 |
| 31 | O35598 | Adam10 | 2.63 | 0.0008 | 67 | Q3V3E1 | Ubash3a | 0.97 | 0.0036 |
| 32 | E9QJS7 | Adgre5 | 2.54 | 0.0013 | 68 | Q8C0Z1 | Fam234a | 0.91 | 0.0145 |
| 33 | Q8C503 | Sit1 | 2.49 | 0.0009 | 69 | G3X9T7 | Lgals9 | 0.82 | 0.0376 |
| 34 | Q8VDN2 | Atp1a1 | 2.48 | 0.0001 | | | | | |
